# Hypertension-Induced Neurovascular and Cognitive Dysfunction at Single-Cell Resolution

**DOI:** 10.1101/2025.04.14.648770

**Authors:** Samantha M. Schaeffer, Anthony G. Pacholko, Monica M. Santisteban, Sung Ji Ahn, Gianfranco Racchumi, Gang Wang, Laibaik Park, Giuseppe Faraco, Josef Anrather, Costantino Iadecola

## Abstract

Arterial hypertension is a leading cause of cognitive impairment, attributed to cerebrovascular insufficiency, blood-brain barrier disruption, and white matter damage. However, the molecular mechanisms by which hypertension affects brain cells remain unclear. Using scRNA-seq in a mouse model of hypertension induced by angiotensin II, we mapped neocortical transcriptomic changes before (3 days) and after (42 days) onset of neurovascular and cognitive deficits. Surprisingly, evidence of endothelial transport disruption and senescence, stalled oligodendrocyte differentiation, interneuronal hypofunction and network imbalance emerged after just 3 days. By 42 days, when cognitive impairment becomes apparent, deficits in myelination and axonal conduction, as well as neuronal mitochondrial dysfunction developed. These findings reveal a previously unrecognized early vulnerability of endothelial cells, interneurons, and oligodendrocytes, and provide the molecular bases for the subsequent neurovascular dysfunction and cognitive impairment in hypertension. In addition, the data constitute a valuable resource for future mechanistic studies and therapeutic target validation.

## INTRODUCTION

Arterial hypertension (HTN) is estimated to affect 1.28 billion people worldwide and is one of the major contributors to the global disease burden^1^. HTN has devastating effects on the brain that are key drivers of the associated morbidity and mortality^2^. HTN is the leading risk factor for stroke and vascular cognitive impairment^2^, but also for Alzheimer’s disease (AD), the major cause of cognitive decline in the elderly^2,3^. Anti-hypertensive pharmacotherapy has greatly reduced cardiovascular mortality, including stroke^4^, but the benefit on cognitive impairment remain less clear^5^. Indeed, the risk of dementia in HTN has increased dramatically over the past 2 decades^6^, highlighting the need to gain a deeper understanding of how HTN affects brain function^2^.

The brain’s structural and functional integrity depends on the blood flow-dependent delivery of O_2_ and glucose, commensurate to the local energy demands of brain activity (neurovascular coupling), and on the homeostasis of its internal milieu safeguarded by the blood brain barrier (BBB)^7,8^. These critical functions are carried out through the close interaction between vascular cells (endothelial and mural cells), glia, and neurons, forming a functional unit known as the neurovascular unit (NVU)^7,8^. In humans as in animal models, HTN has profound effects on neurovascular cells, causing endothelial dysfunction, disruption of the BBB, and suppression of neurovascular coupling^9–16^. In turn, these interrelated pathogenic processes lead to an increased susceptibility of the brain to dysfunction and damage^2^. The subcortical white matter is particularly vulnerable to the damaging effects of HTN, and white matter disruption is the leading cause of cognitive decline in individuals with high blood pressure (BP)^17–20^. However, the cellular and molecular events underlying the NVU impairment in HTN, and their impact on neuronal networks activity and white matter integrity remain poorly understood.

Administration of angiotensin II (AngII), an octapeptide central to the pathobiology of human HTN^21^, reproduces the damaging neurovascular and cognitive effects of HTN in several animal models^22^. Notably, administration of low doses of AngII in mice over a period of weeks elicits a slow-developing rise in BP over several days, known as “slow pressor” HTN^22–24^. In this model, circulating AngII does not elevate BP by constricting peripheral resistance vessels directly, since the dose is too low to induce acute vasoconstriction^21,25^. Rather, a key factor mediating the HTN is activation of AngII type 1 receptors (AT1R) in the subfornical organ, one of the circumventricular organs, leading to sympathetic activation, hormonal release, and renal oxidative stress (neurohumoral dysfunction)^23,24,26^. Since neurohumoral dysfunction and oxidative stress are key features of essential hypertension, the most common form of hypertension in humans^27^, the slow pressor model is frequently used to investigate HTN-induced and organ damage^22^. Accordingly, this model reproduces selected neurovascular and cognitive alterations observed in hypertensive individuals albeit on a more compressed time scale^9,28–30^. For instance, endothelial dysfunction, neurovascular uncoupling, and BBB leakage emerge within 1–2 weeks of AngII administration, preceding the onset of cognitive deficits, which typically appear around the fourth week^14,31–33^. In contrast, administration of pressor doses of AngII induces acute hypertension and rapid-onset neurovascular dysfunction^15,16,34^. Thus, the temporal delay between AngII infusion and the onset of neurovascular and cognitive impairment in the slow pressor model makes it particularly well suited for investigating early molecular changes in NVU cells that may underlie and drive subsequent pathogenic outcomes.

The recent application of single-cell, single-nuclei RNA sequencing (scRNA-seq) to the study of the neurovasculature and associated cells has unveiled a remarkable molecular heterogeneity of cerebrovascular cells and shed light on the molecular bases for interactions with other cells in the NVU^8,35–39^. Given the profound impact of HTN on neural and vascular function, scRNA-seq could offer an assumption-free assessment of the cell-type-specific transcriptomic changes associated with the resulting neurovascular and cognitive dysfunction. Yet, no studies to date have been performed to fill such knowledge gap. To explore this, we employed scRNA-seq to examine the transcriptomic changes induced by HTN on NVU cells in mice treated with AngII at two time points: before (3 days; 3D) and after (42 days; 42D) the full development of neurovascular and cognitive dysfunction. Our data revealed profound transcriptional alterations reflecting endothelial senescence at the capillary-venular transition, BBB alterations, disruption of oligodendrocyte maturation, and dysfunction of inhibitory interneurons, all of which, surprisingly, emerged after just 3D of AngII infusion. By 42 days, when cognitive impairment was observed, deficits in myelination and axonal conduction emerged, accompanied by neuronal mitochondrial dysfunction. The data uncover the molecular bases of a previously unrecognized early vulnerability of endothelial cells at the capillary-venule transition, interneurons, and oligodendrocytes to AngII-induced hypertension (AngII HTN), which precedes neurovascular and BBB dysfunction. They also highlight the molecular changes underlying the subsequent white matter and neural network impairment that contribute to cognitive deficits in AngII HTN. Finally, the data serve as a valuable resource for future mechanistic studies and for validation of potential therapeutic targets identified through the current molecular investigations. To this end, a portal (https://anratherlab.shinyapps.io/angii_brain/) has been established for user-friendly access to our data.

## RESULTS

### SINGLE-CELL TRANSCRIPTOMIC PROFILING OF NEOCORTICAL CELLS IN ANGII HTN

We induced “slow pressor” HTN in 10-week-old C57BL/6J male mice through infusion of low concentrations of AngII (600 ng/kg/min) using subcutaneously (*s.c.*) implanted osmotic mini pumps (n = 6-8/group). Mice receiving vehicle (saline) served as controls. As previously reported^14^, AngII treatment elicited a gradual increase in systolic BP beginning at 3 days (3D) and continuing through the full 42 days (42D) of infusion (Fig.1A), when cognitive impairment was observed in the novel object recognition test (Fig. 1B). We used young male mice because female mice are resistant to slow pressor AngII HTN^40^ and aging has confounding effects on the neurovasculature^41^ that would confound the interpretation of the results.

**Figure 1.**
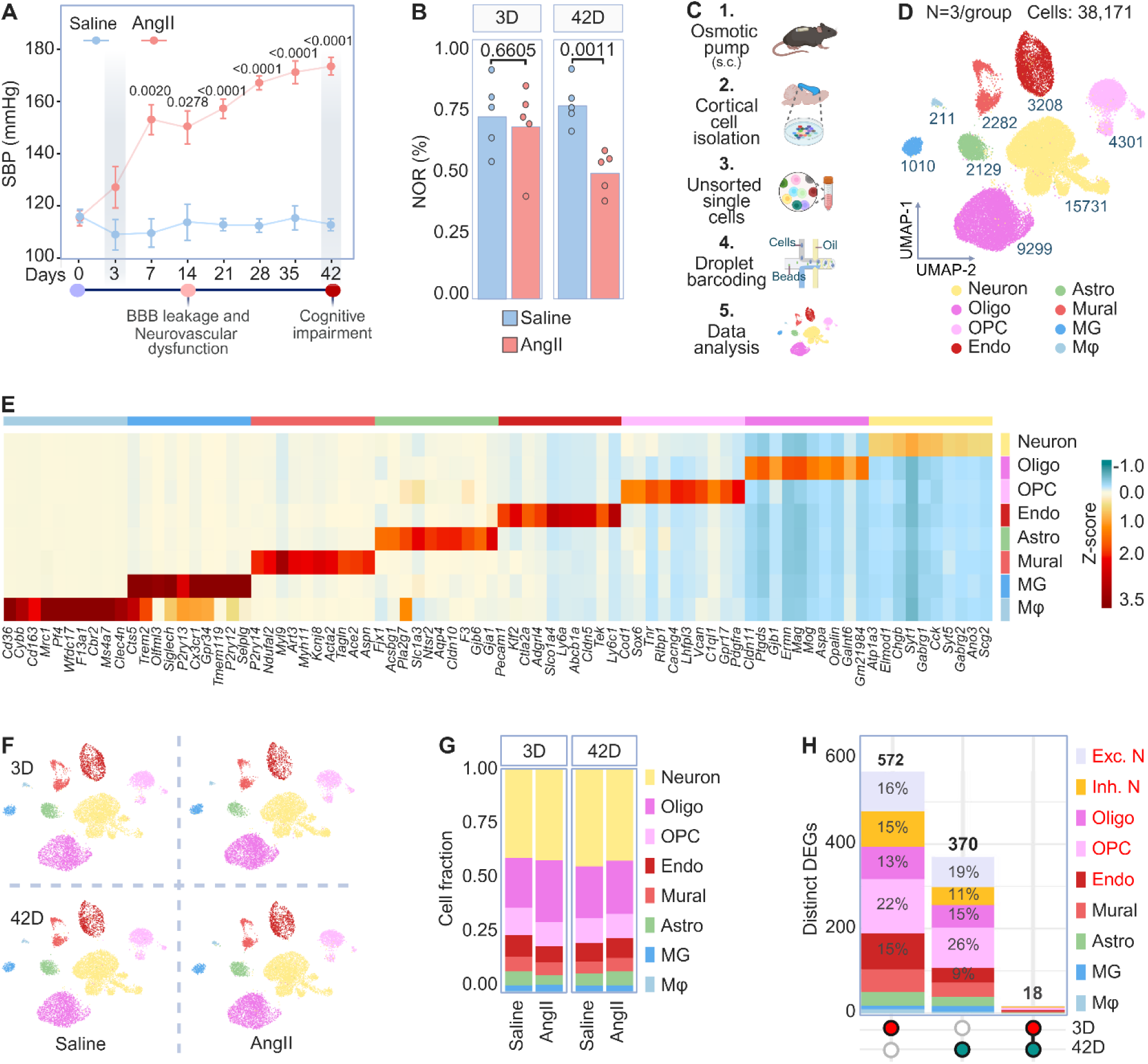
Single-cell transcriptomic profiling of mouse cortical cells in early- and late-stage AngII HTN. **(A)** Line graph depicting the gradual elevation in BP elicited by slow pressor AngII infusion. Measurements performed by tail-cuff plethysmography. The onset of BBB, neurovascular, and cognitive dysfunction is indicated based on refs^13,14,33^. N = 6-8/group. Data represents mean ± SEM. Statistical analysis performed using repeated measures two-way ANOVA with Šidák’s correction. **(B)** Bar graph showing recognition memory assessed by the novel object recognition test (NOR) in saline- and AngII-treated mice after 3 days (left) or 42 days (right). \*\**p*<0.01; n = 5/group. Data represents mean ± SEM. Statistical analysis performed using Student’s *t*-test for single contrast of intergroup differences. **(C)** Schematic overview of the Drop-Seq scRNAseq pipeline used to analyze brain cortical cells isolated from mice treated for 3 or 42 days with saline or AngII via subcutaneously implanted osmotic minipumps. **(D)** Uniform Manifold Approximation and Projection (UMAP) plot depicting color-coded cell clusters identified in merged 3-day and 42-day saline and AngII single-cell transcriptomes: neurons, oligodendrocytes (Oligo), oligodendrocyte precursors (OPC), endothelial cells (Endo), mural cells (Mural), astrocytes (Astro), microglia (MG), and macrophages (Mφ). **(E)** Heatmap displaying the expression of the top ten upregulated genes in each general cluster. Scale bar represents z-score of the average log gene expression. **(F)** UMAP of overlaid time points and treatments representing overlapping clusters. **(G)** Bar graph showing the relative frequencies of each cluster across time points and treatments. **(H)** UpSet plot depicting differentially expressed genes (DEG) by cell type at day 3, day 42, and both day 3 and 42 in response to AngII treatment. On the x axis, red circles indicate DEG at day 3 (left), blue circles DEG at day 42 (middle), and both red and blue circles DEG common to both time points (right). Percentages reflect the proportion of DEG for each cell type at each time point.

To delineate the transcriptomic changes at the single cell level preceding or concurrent with the full development of neurovascular and cognitive impairment, we performed high throughput scRNA-seq of neocortical cells from mice treated with AngII for 3D or 42D (n=3 biological replicates per group, 2 pooled brains per replicate). We used cortical tissue because the neurovascular effects of AngII HTN are well described in the cortex^13–16,33^ and for the relevance of cortical circuits to cognitive dysfunction^18–20^. Dissected cortices were processed via gentle papain digestion and single cell transcriptome libraries were prepared using the Drop-seq platform (Fig.1C)^42–44^. After quality control, we obtained 38,171 single cell transcriptomes. Visualization in Uniform Manifold Approximation and Projection (UMAP) space separated cells into 9 distinct classes (Fig.1D). Using established marker genes and unsupervised cell-type annotation (Fig.1E), clusters were identified as neurons, mature oligodendorcytes, oligodendrocyte precursor cells (OPC), endothelial cells (EC), astrocytes (Astro), microglia (MG), mural cells (smooth muscle cells and pericytes), and macrophages (Mφ). Separate UMAP representations indicated that neither treatment nor timepoint affected the frequency or distribution of the major cell types or their subclusters (Fig.1F-G). Analysis of the differentially expressed genes (DEG) between AngII and saline treated mice revealed that the majority of DEG were timepoint and cell-type specific (Fig.1H). The most pronounced transcriptional responses occurred in EC, oligodendrocyte-lineage cells, and neurons, particularly at the 3D timepoint. Less pronounced changes were seen in mural cells (Fig. S4), astrocytes (Fig. S5), and microglia (Fig. S6), and cell counts for Mφ did not reach the threshold for further analysis. Accordingly, transcriptomic changes in EC, oligodendrocytes and neurons will be described first.

### ENDOTHELIAL CELL CLUSTERS ALIGN ALONG AN ARTERIAL-VENOUS AXIS

To characterize cerebral EC transcriptomics in AngII HTN, we performed fine classification of EC subtypes within the cerebrovascular tree^39,42^. Unbiased sub-clustering of 3,232 ECs revealed 4 continuous populations consistent with the gradual molecular and phenotypic transitions which occur along the arterial-venous axis (Fig. 2A, Fig. S1A), as previously described^39^. Subtypes were defined by automated-hierarchical annotations and established expression patterns of zonation genes (Fig. S1B-C), revealing: 1) arteriolar (aEC; *Bmx*), 2) arteriolar-capillary (aCapEC; *Tm4sf1, Rgcc*), 3) capillary-venular (vCapEC; *Rgcc, Car4*), and 4) venular (vEC; *Car4*, *Slc38a5*) segments^45,46^. Based on *Vcam1* expression, which correlates with vessel size^47^, we distinguished aEC and vEC from capillary EC and putatively classified them as pre- and post-capillary, respectivel*y.* Consistent with recent *in silico* reports employing lineage tracing on EC transcriptomes^38,48^, EC subclusters were further grouped into arteriolar and venular ‘zones’ based on the increased expression of *Car4* in vCapEC and vEC populations. RNAscope in naïve mice revealed that (a) *Car4* mRNA colocalizes with the venous marker *Slc38a5*^46^ (Fig. S1D), (b) is mainly expressed in CD31^+^ microvessels with a lumen diameter of 7-10 µm (Fig. S1E-F), and (c) is not observed in smooth muscle associated vessels (αActa2+) (Fig. S1G), validating its use as a marker of ECs at the transition between capillary and venous microvasculature (vCapEC). These observations define the endothelial clusters of interest and their microvascular zonation.

**Figure 2.**
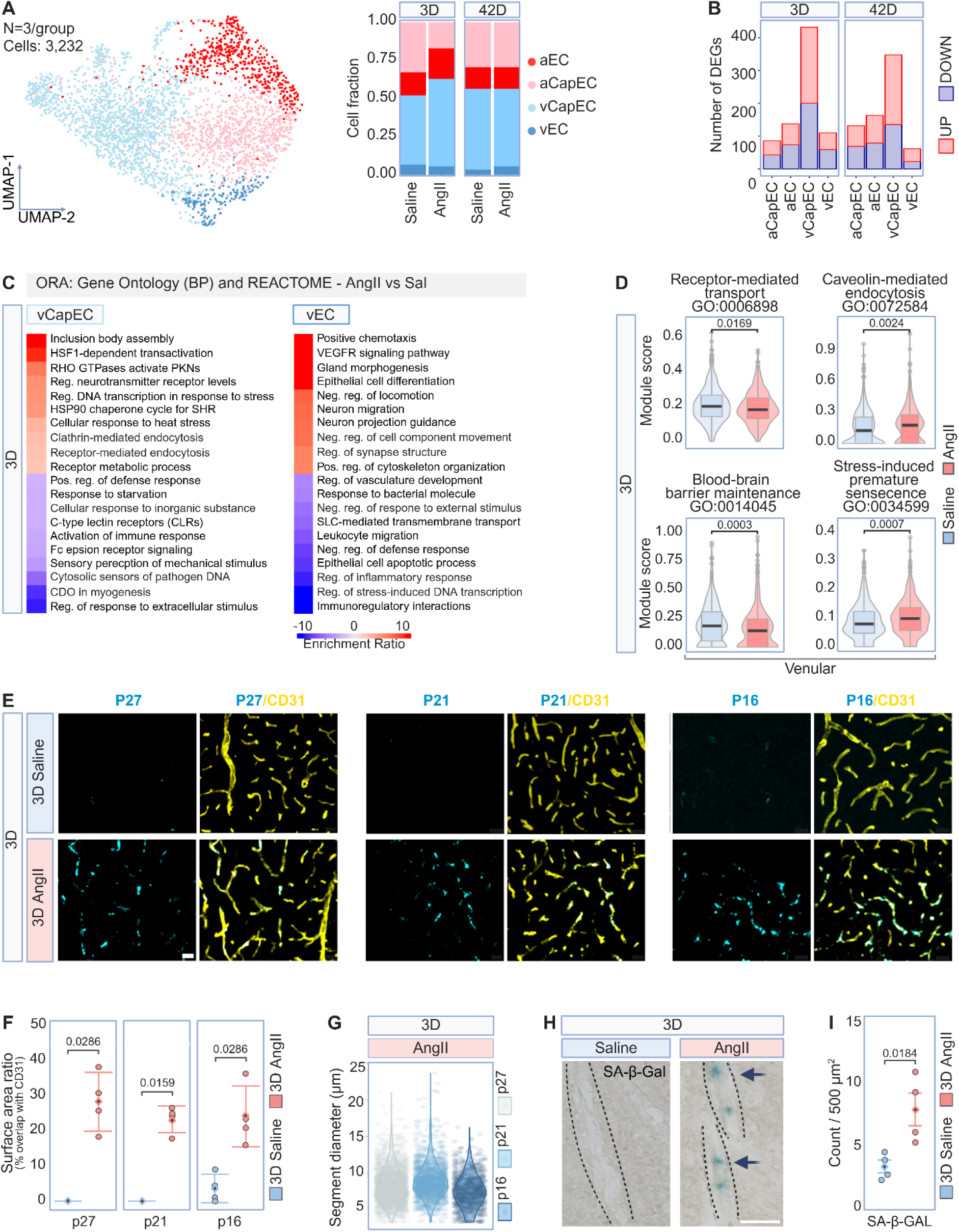
AngII HTN induces senescence in venular endothelial cells. **(A)** Left: UMAP representation of 3,232 endothelial cells showing the 4 subtypes identified in merged single cell transcriptomes: arterial (aEC), arterial capillary (aCapEC), venous (vEC), and venous capillary (vCapEC). Right: Bar graph showing the relative frequencies within each sub-cluster across time points and treatments. Also see Figure S1. **(B)** Bar graph depicting the number of significant upregulated and downregulated differentially expressed genes (DEG) from each subcluster in response to 3 days or 42 days of AngII. **(C)** Heatmaps representing enriched biological pathways after 3D of AngII treatment in endothelial cell subtypes determined through overrepresentation analysis (ORA) of upregulated and downregulated DEG (*p*<0.05, logFC>2). Pathways are derived from the Gene Ontology Biological Process and Reactome libraries. Scale bar represents the enrichment score of genes within a given pathway. **(D)** Violin and box plots showing the results of module score analyses for select gene sets at the 3D timepoint. *p*<0.05. Boxplots represent the median ± IQR. Statistical analysis performed using the Wilcoxon test, AngII versus saline. **(E)** Immunostaining showing colocalization of senescence markers p27^Kip1^ (cyan), p21^WAF1/Clip1^ (cyan), and p16^INK4A^ (cyan) with microvascular CD31^+^ endothelial cells (yellow) in the cortex of mice treated for 3 days with saline or AngII. Scale bar = 20 µm. **(F)** Dot plots depicting the quantification of senescence marker colocalization with CD31^+^ endothelial cells. N = 4 brains/group. Data represents mean ± SEM. Statistical analysis performed using the Wilcoxon test, saline versus AngII. **(G)** Violin plots depicting senescence marker expression stratified by vessel diameter. n = 4 brains/group. **(H)** Light microscopic images of senescence-associated β-galactosidase (SA-β-gal) staining in cortical vessels derived from mice treated for 3 days with saline or AngII. Scale bar = 20 µm. **(I)** Dot plots of SA-β-gal quantification in the cortex. Data represents mean ± SEM, with each individual data point representing the average count of SA-β-gal positive cells found per 500 um^2^ section. N = 4 brains/group, 5 sections/brain. Statistical analysis performed using an unpaired, two-tailed t-test.

### THE CAPILLARY-VENOUS ENDOTHELIUM IS A KEY TARGET OF ANGII HTN

Next, we investigated whether AngII HTN exhibits segment-specific transcriptomic effects. First, DEG were calculated for each EC subtype in 3D and 42D AngII treatment groups relative to their corresponding saline controls. EC within the venular zone exhibited significantly more DEGs than those within the arteriolar zone (Fig.2B), highlighting the exquisite sensitivity of the venous microvasculature to the effects of AngII HTN^14^. A substantial portion of the DEG emerged after just 3D of AngII treatment (Fig.2B), with over-representation analysis (ORA) and supervised module scoring revealing several aging-related transcriptional phenotypes (Fig.2 C-D). As described in the aging endothelium^41,49^, genes involved in energy homeostasis (*Slc38a5, Slc1a2, Cox6a1, Sesn1, Suclg1*) were downregulated in the venular axis concurrent with a reciprocal increase in those pertinent to oxidative fatty acid metabolism (*Acot7, Oxct1, Pnpla8*) (Fig.2C). Additionally, 3D of treatment with AngII upregulated the expression of genes in vCapEC and vEC indicative of transport-related aberrations (*Cav1, Picalm, Nedd4, Ttr, Atp1a2*) coincident with downregulation of those involved in endocytic cargo-recognition (*Itsn2, Ubc, Lgals9, Sgk1, Psap,, Diaph1*) (Fig. 2C). Subsequent module scoring confirmed that these changes reflect a shift in endothelial transport from ligand-receptor interactions mediated by solute carriers, to non-specific caveolar transcytosis (Fig. 2D), a phenotype reported in the aging microvasculature^50^. Furthermore, AngII suppressed the expression of β-catenin/Wnt signaling genes (*Fzd4, Plcb1, Psma1*) in vEC and vCapEC, which are required for maintenance of the BBB^51^, and upregulated *Nedd4* concurrent with suppression of *Mfsd2a*, indicating an increase in transcytosis associated with impaired BBB integrity^14,52,53^. Module scoring confirmed a focal decrease in gene sets involved in BBB maintenance (Fig.2D). Although the permeability of the blood-brain barrier (BBB) to small molecules is not yet increased at 3D in this model^13^, these observations suggest that aberrations in endothelial transport mechanisms occur early in slow pressor AngII-HTN, providing the molecular basis for the BBB dysfunction observed later on (14 days)^14^.

### ANGII HTN INDUCES A SENESCENCE PHENOTYPE IN CAPILLARY-VENULAR ENDOTHELIAL CELLS

AngII treatment for 3D induced enrichment of genes involved in cellular senescence^54^, particularly in vEC and vCapEC (*Cdkn1b, H2afv, Stip1, Atf3, Nck1, Arpc3, Klf2, Dll4, Pip5k1c, Tab2, Cyld, Nedd4, Hsbp1, Hist1h2bc, H3f3a*) (Fig.2C). Oxidative stress is a potent driver of cellular senescence^54^ and, accordingly, module scoring revealed that cellular senescence in vEC and vCapEC was coupled to oxidative stress-induced premature senescence (SIPS), a reactive oxygen species (ROS) driven induction of p38 MAPK signaling^55,56^ (Fig.2D). Markers of cellular senescence include upregulation of cyclin-dependent kinase inhibitors (i.e. p27^Kip1^, p21^WAF1/Cip1^, p16^INK4A^), suggesting cell cycle arrest^57^, and the induction of senescent-associated β-galactosidase activity (SA-β-gal), reflecting enlarged lysosomal compartments, decreased functional capacity, and metabolic dysfunction^57^. Therefore, we next confirmed the presence of these indicators *ex vivo*^57^. In mice treated for 3D with AngII, p27^Kip1^(+29 ± 8%), p21^WAF1/Cip1^ (+24 ± 4%), and p16^INK4A^ (+24 ± 9%) immunoreactivity was enriched in CD31^+^ microvessels (Fig.2E-F) less than 20 µm in diameter (Fig.2G), and histochemical staining for SA-β-gal (Fig.2H) revealed elevated enzymatic activity in cells localized to the venous microvasculature (Fig.2I).

To assess whether HTN-induced endothelial senescence could be reversed by lowering BP, we induced AngII slow pressor HTN and, once the BP was elevated (day 5), treated mice with the AT1R antagonist losartan (600mg/L; in the drinking water)^58,59^. Losartan reversed the HTN within 1 day (Fig. 3A) and attenuated senescence markers in cerebral endothelial cells assessed at day 8 (Fig. 3B-C). To determine whether the losartan-induced attenuation of endothelial senescence was due to inhibition of AngII signaling rather than a reversal of the mechanical effects of elevated blood pressure, HTN was induced using phenylephrine, a peripheral vasoconstrictor that does not cross the blood-brain barrier^33^. We found that phenylephrine elevated BP (Fig. 3D) but did not induce endothelial senescence (Fig. 3E-F). Conversely, lower doses of AngII that do not increased BP (non-pressor dose; 200 ng/kg/min)^33^, induced senescence markers in cerebral endothelial cells (Fig. 3E-F). These observations demonstrate that endothelial senescence is downstream of AT1R signaling and independent of the elevation in BP.

**Figure 3.**
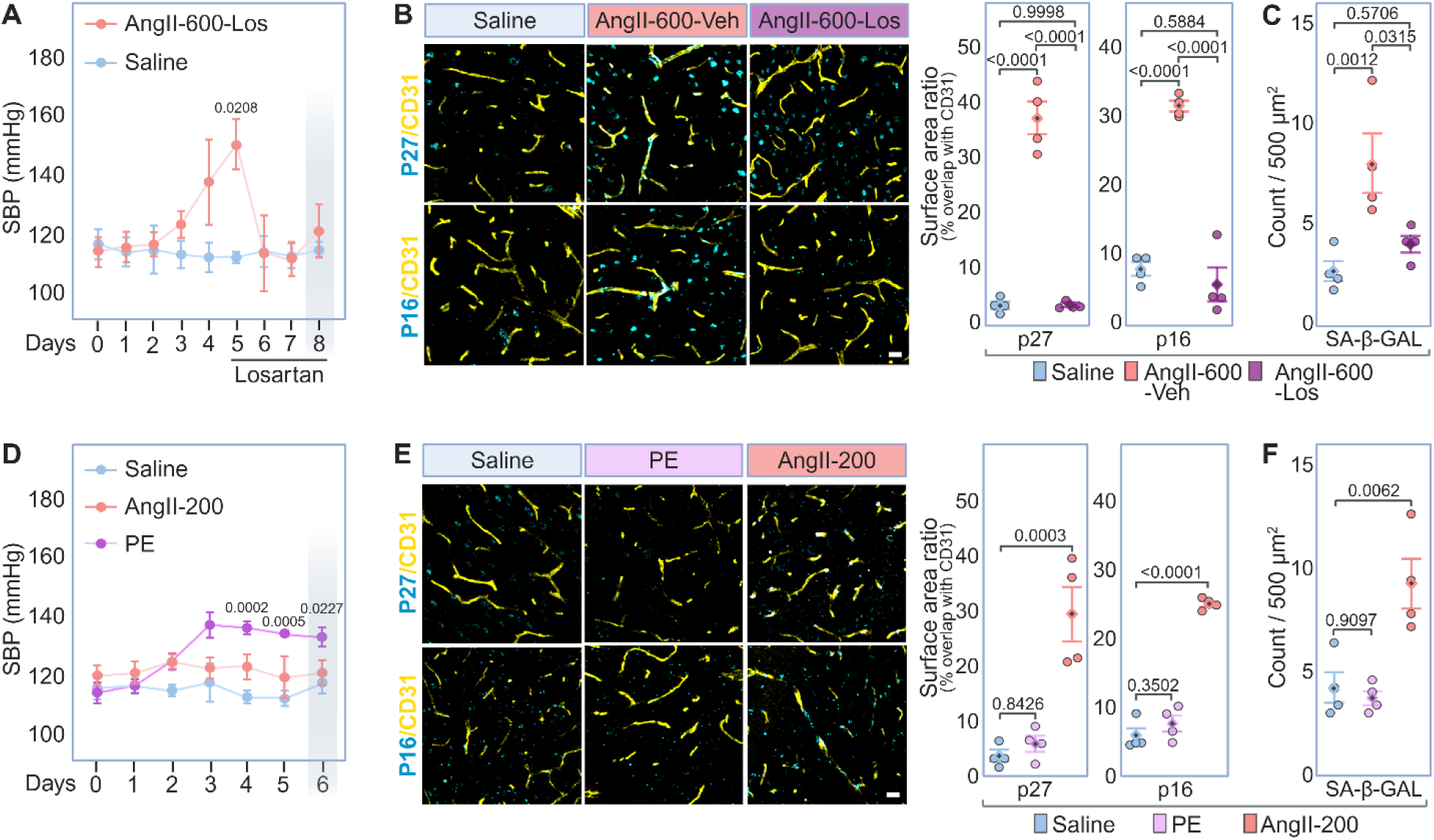
Senescence in endothelial cells is induced by AngII independently of the elevated SBP and is reversed by the AT1R blocker losartan. **(A)** Line graph depicting the gradual elevation in BP elicited by slow pressor AngII infusion followed by subsequent attenuation by losartan. Losartan was administered in the drinking water from day 5 through 8. Measurements performed daily by tail-cuff plethysmography. N = 5/group. Data represents mean ± SEM. Statistical analysis performed using repeated measures two-way ANOVA with Šidák’s correction. **(B)** Left: Immunostaining showing colocalization of senescence markers p27^Kip1^ (cyan) and p16^INK4A^ (cyan) with the endothelial cells marker CD31^+^ (yellow) in the cortex of mice treated with saline, slow-pressor AngII, or slow-pressor AngII with or without Losartan. N = 4 brains/group. Scale bar = 20 µm. Right: Dot plots depicting the quantification of senescence marker colocalization with CD31^+^ endothelial cells. N = 4 brains/group. Data represents mean ± SEM. Statistical analysis performed using one-way ANOVA with Tukey’s test. **(C)** Dot plots of SA-β-gal quantification in the cortex. Data represents mean ± SEM, with each individual data points representing the average count of SA-β-gal positive cells found per 500 µm^2^ section. N = 4 brains/group, 5 sections/brain. Statistical analysis performed using one-way ANOVA with Tukey’s test. **(D)** Line graph depicting BP during administration of saline, a non-pressor dose of AngII and a pressor dose of the peripheral vasoconstrictor phenylephrine. Measurements performed by tail-cuff plethysmography. N = 5/group. Data represents mean ± SEM. Statistical analysis performed using repeated measures two-way ANOVA with Šidák’s correction. **(E)** Left: Immunostaining showing colocalization of p27^Kip1^ (cyan) and p16^INK4A^ (cyan) with microvascular CD31^+^ endothelial cells (yellow) in the cortex of mice treated with saline, phenylephrine or non-pressor AngII. N = 4 brains/group. Scale bar = 20 µm. Right: Dot plots depicting the quantification of senescence marker colocalization with CD31^+^ endothelial cells. N = 4 brains/group. Data represents mean ± SEM. Statistical analysis performed using one-way ANOVA with Dunnet’s test. **(F)** Dot plots of SA-β-gal quantification in the cortex. Data represents mean ± SEM, with each individual data points representing the average count of SA-β-gal positive cells found per 500 um^2^ section. N = 4 brains/group, 5 sections/brain. Statistical analysis performed using one-way ANOVA with Dunnet’s test.

### TRANSCRIPTOMIC CHANGES IN ARTERIOLAR ENDOTHELIUM SUGGEST REMODELING

After 42D of AngII treatment significant alterations emerged in arteriolar segments. In aEC and aCapEC, genes involved in remodeling (*Cav1, Adam15, Itgb1, Phactr4, Ecm2, Cdc42, Jun, Junb, CD81, Rhob, Myh9, Fam107a*), and collagen metabolism (*Adam15, Serpinh1, Itgb1, Ppib*) were upregulated, while genes associated with redox homeostasis were downregulated (*Txnl1, Tmx3, Tmx4, Txnrd1*) (Fig. 4A). Subsequent module scoring confirmed an increase in blood vessel remodeling and collagen biosynthesis related genes (Fig. 4B). Histological studies confirmed the increase in collagen IV in the microvasculature, consistent with vascular remodeling (Fig. 4C-D). On the venular side, 42D of AngII upregulated expression of *Vcam1* and *Vcam1*-mediated immune cell transmigration pathway genes (*i.e.*, *Esam*, *Ctnna1, Sema3c, Pak1, Tab2, Jak1, Vamp2, Ecscr*), reflecting an aging-like phenotype implicated in leukocyte tethering, endothelial activation, and inflammation^60^.

**Figure 4.**
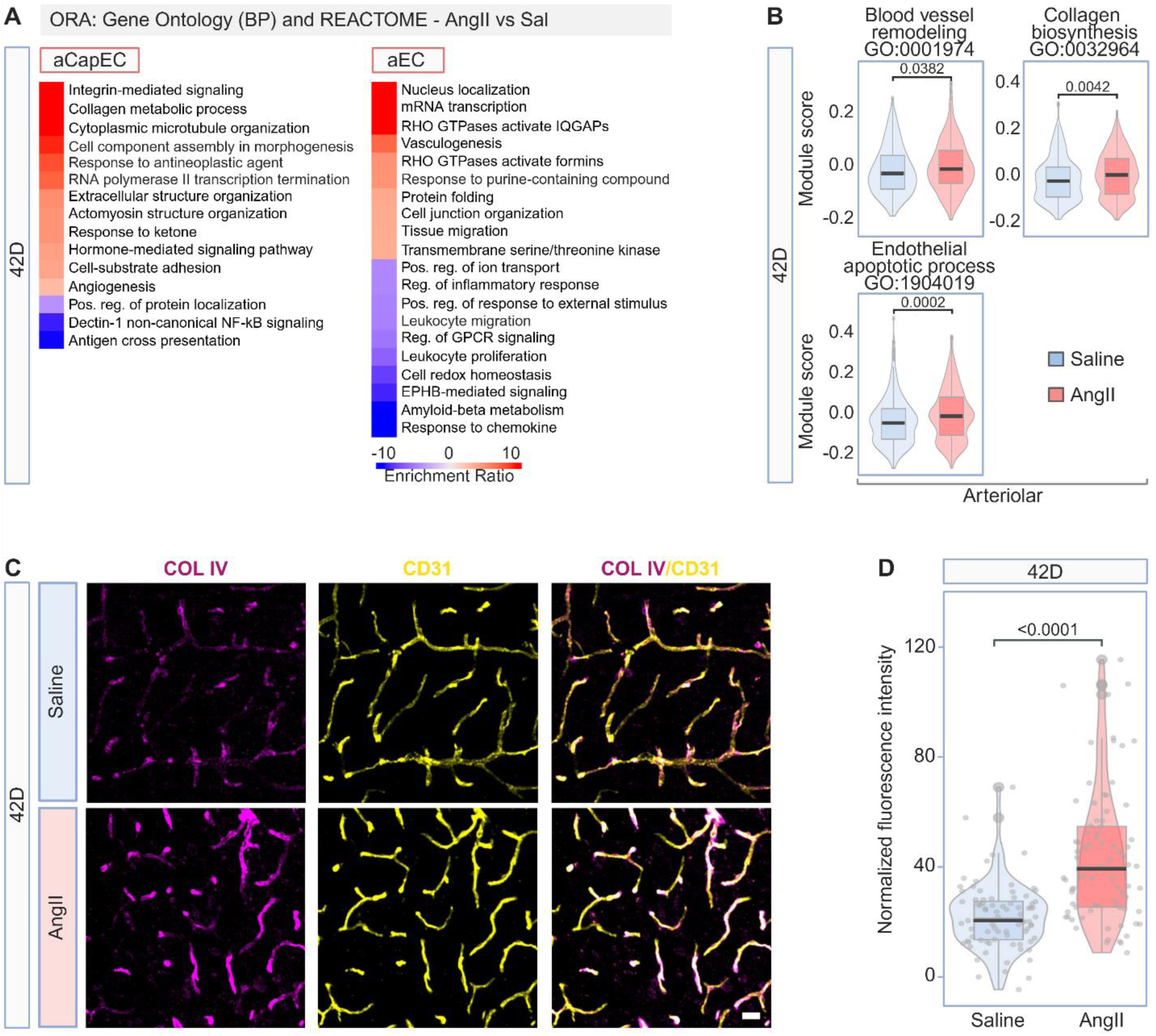
AngII HTN promotes remodeling in arteriolar endothelial cells. **(A)** Heatmaps representing enriched biological pathways after 42 days of AngII treatment in endothelial cell subtypes determined through ORA of upregulated and downregulated DEG (*p*<0.05, logFC>2). Pathways derived from the Gene Ontology Biological Process and Reactome libraries. Scale bar represents the enrichment score of genes within a given pathway. **(B)** Violin and box plots showing the results of module score analyses for select gene sets at the 42-day timepoint. *p*<0.05. Boxplots represent the median ± IQR. Statistical analysis performed using Wilcoxon test, AngII versus saline. **(C)** Immunostaining showing colocalization of collagen IV (COL IV, magenta) with microvascular CD31^+^ endothelial cells (yellow) in the cortex of mice treated with saline or slow-pressor AngII for 42 days N = 4 brains/group. Scale bar = 20 µm. **(D)** Combined violin and jitter plot depicting the quantification of COL IV expression in CD31^+^ cerebral microvessels. N = 4 brains/group. Data represents mean ± SEM. Statistical analysis performed using the Mann-Whitney test. Dots represent the COL IV fluorescence signal in individual vessel segments.

Collectively, these observations indicate that, at the transcriptional level, pathological changes consequent to AngII HTN begin in the venular endothelium with a pattern of premature vascular aging and senescence-related changes. Over time, changes are also observed in the arteriolar endothelium, where remodeling responses emerge.

### TRANSCRIPTOMIC CHANGES INDICATE STALLED OLIGODENDROCYTE DEVELOPMENT IN ANG HTN

The clustering of 13,600 oligodendrocytes revealed 5 distinct populations: cycling progenitors (CyP), oligodendrocyte progenitor cells (OPC), differentiation-committed oligodendrocyte precursors (COP), newly formed oligodendrocytes (NFOL), and mature oligodendrocytes (MOL), as previously described^61,62^ (Fig. S2A). Cells were further manually assigned to 3 clusters reflecting the continuous maturation process: OPC, pre-oligodendrocytes (pre-OL), and MOL (Fig. 5A, Fig. S2A-D). Notably, OPC were disproportionately affected at both timepoints (Fig. 5B), suggesting an early and sustained vulnerability to AngII HTN.

**Figure 5.**
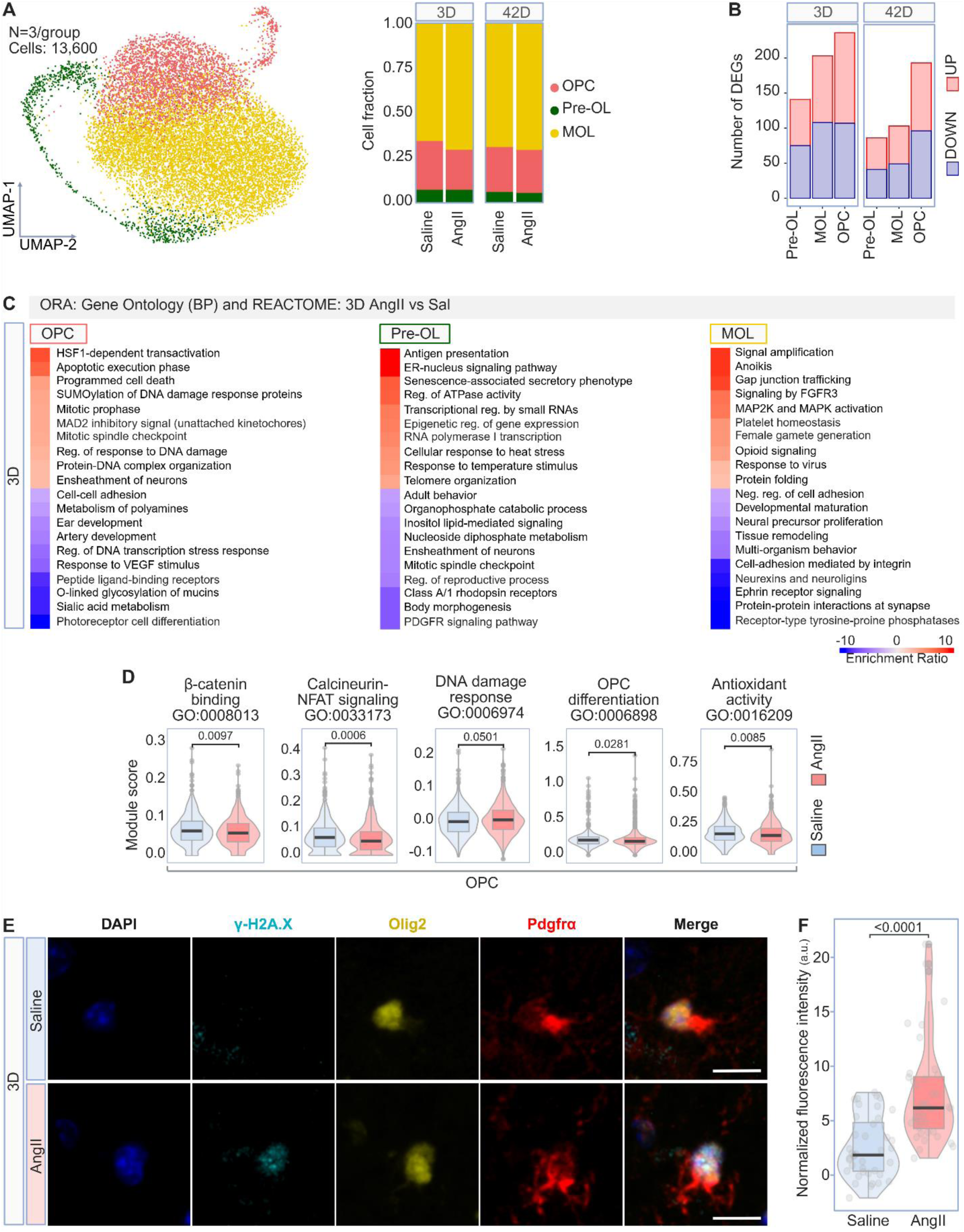
Oligodendrocyte precursor cells are affected early in AngII HTN. **(A)** Left: UMAP depicting the organization of 13,600 oligodendrocyte-lineage cells into 3 subtypes characterized by degree of maturation: oligodendrocyte precursors (OPC), pre-myelinating oligodendrocytes (Pre-OL), and mature oligodendrocyte (MOL). Right: Bar graph showing the relative frequencies within each sub-cluster across time points and treatments. Also see Figure S2. **(B)** Bar graph depicting the number of significant upregulated and downregulated differentially expressed genes (DEG) in each subtype in response to 3 days or 42 days of AngII. **(C)** Heatmaps representing the enriched biological pathways after 3 days of AngII treatment in oligodendrocyte-lineage cell subtypes determined through overrepresentation analysis (ORA) of upregulated and downregulated DEG (*p*<0.05, logFC>2). Pathways are derived from the Gene Ontology Biological Process and Reactome libraries. Scale bar represents the enrichment score of genes within a given pathway. **(D)** Violin and box plots showing the results of module score analyses for select gene sets. *p*<0.05. Boxplots represent the median ± IQR. Statistical analysis performed using the Wilcoxon test, AngII versus saline. **(E)** Immunostaining showing colocalization of the double-strand DNA break marker γ-H2A.X (cyan) with OPC identified by co-expression of Olig2 (yellow) and Pdgfrα (red) in the cortex of mice treated for 3 days with saline or slow-pressor AngII. Cell nuclei are stained with 4′,6-diamidino-2-phenylindole (DAPI, blue). N = 4 brains/group. Scale bar = 10 µm. **(F)** Combined violin and jitter plot depicting the quantification of γ-H2A.X fluorescence intensity in OPC nuclei. N = 4 brains/group. Data represents mean ± SEM. Statistical analysis performed using the Mann-Whitney test. Dots represent individual cells.

Owing to their high metabolic demands and relative paucity of antioxidant enzymes, OPC are selectively vulnerable to oxidative stress^63,64^. Accordingly, in OPC, treatment with AngII for 3D upregulated genes associated with DNA damage (*Lmnb1, Casp3, Hmgb2, Hist1h1e, Seh1l, Nup37, Nsmce4a, Sumo1, Smc5, Fbxo5, Kmt5a, Mad2l2, Atad5, Mcl1, Dhx9)* and, consistent with the inhibitory effect of oxidative damage on OPC differentiation^65^, pathways pertaining to cell maturation were downregulated (*e.g.*, photoreceptor cell differentiation, artery development, connective tissue development, urogenital system development) (Fig.5C). While these pathways are not specific to OPC differentiation, they are populated by genes involved in regulation of the proliferation, migration, and maturation of OPC (*Notch1, Bbs4, Cep290, Robo1, Prrx1, Jun, Egr1, Plce1, Kcnma1, Src, Wwtr1, Ptprt, Plxnb3, Bmp1, Shroom2*)^66^. Subsequent module scoring confirmed that AngII HTN upregulated DNA damage response-associated genes in OPC, while those genes involved in β-catenin binding, Nfat/calcineurin signaling, antioxidant capacity, and cell differentiation, which are essential for OPC differentiation, myelin production and antioxidant defense^65,67,68^, were downregulated (Fig.5D). To provide complementary evidence of DNA damage in OPC, we used the DNA damage marker γ-H2A.X^69^. OPC, identified by co-labeling with PDGFRα and Olig2^70^ (Fig. 5E), were found to express γ-H2A.X (Fig. 5F), confirming the transcriptomic findings at the cellular level.

### NODAL DISRUPTION AND SLOWED AXONAL CONDUCTANCE

After 3D of AngII treatment, stress response genes were upregulated in pre-OL (*Hikeshi, H2afz, H3f3a, Hspa2, Ube2e1, Hsph1, Hspa5, Dynll2*) concurrent with downregulation of those associated with neuronal ensheathment (*Cldn11, Mal, Ndrg1, Trf, Nkx6-2, Dag1*) (Fig.5C). Similarly, neurexin-neuroligin signaling was suppressed in MOL (*Nrxn1, Shank1, Lrrtm3, Nlgn3, Lrrtm1*) (Fig.5C). These interactions are required for growth of the myelin sheath^71^, suggesting that processes involved in myelin formation could be disrupted early in AngII HTN and may lead to subsequent deficits in myelin structure. Accordingly, in MOL, 42D of AngII treatment induced a significant downregulation of genes associated with fatty acid metabolism (*Decr1*, *Hacd1*, *Cyp2j12, Pde8a, Nudt, Eno1, Pde7a, Enpp1*), a key process in myelin synthesis^72–74^, while genes involved in bone morphogenetic protein signaling (*Hes5*, *Ryr2*, *Vim*), particularly the powerful myelination inhibitor *Hes5*^75,76^, were upregulated (Fig.6A). Additionally, genes pertinent to the cell cycle (*Smc2, Psme1, H2afv, Pcna, Cep57, Mcm2, Cdc7, Jak2, Nup85, Spc25, Ube2c, Pole3, Sgo2a, Hmmr, Ywhah*) and DNA synthesis (*Psme1, Pcna, Mcm2, Cdc7, Ube2c, Pole3*) were markedly suppressed, suggesting an attenuation of OPC proliferation (Fig.6A).

**Figure 6.**
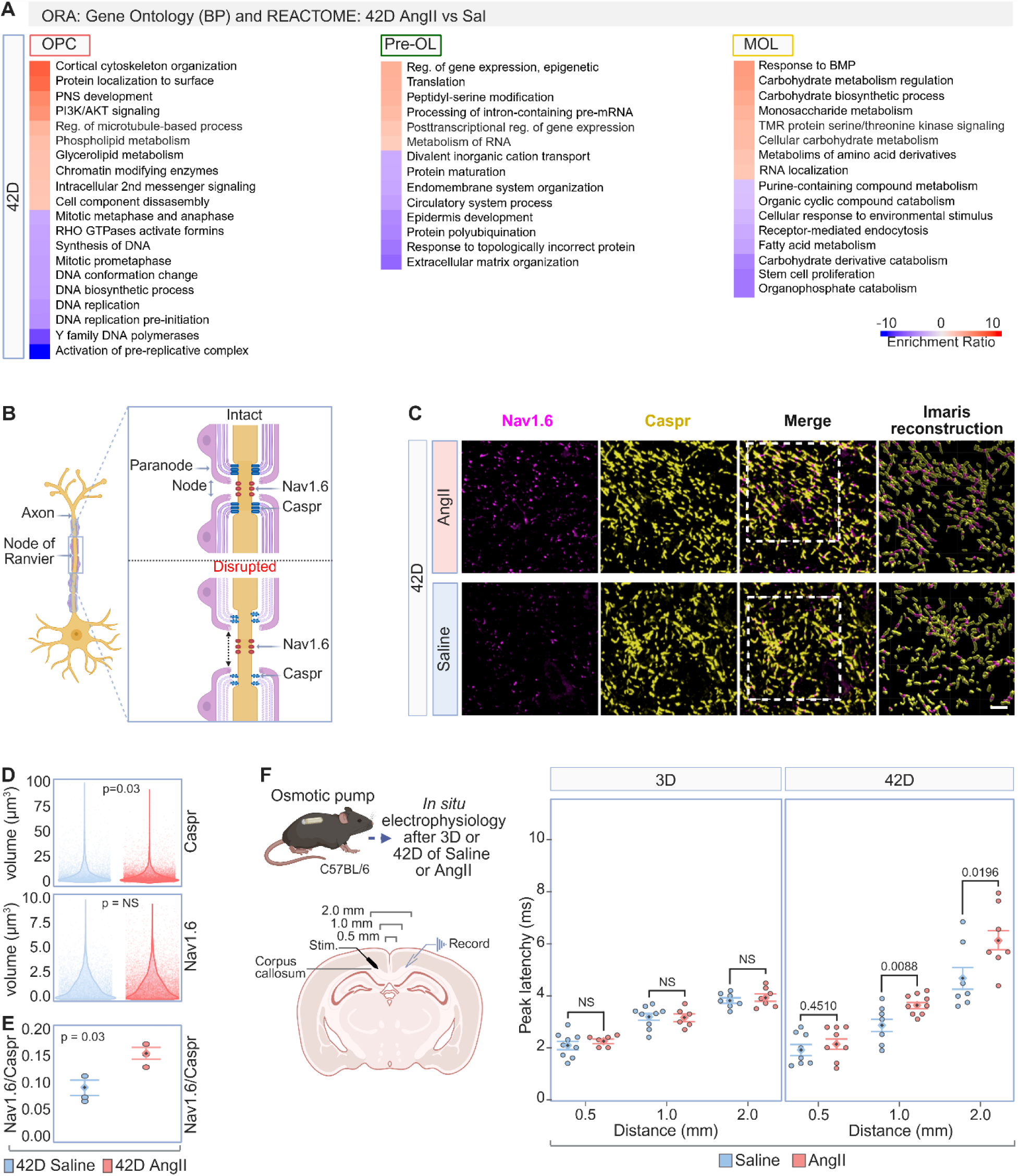
AngII HTN disrupts nodal structure and axonal conductance. **(A)** Heatmaps representing the enriched biological pathways after 42 days of AngII treatment in oligodendrocyte-lineage cell subtypes determined through ORA of upregulated and downregulated DEG (*p*<0.05, logFC>2). Pathways derived from the Gene Ontology Biological Process and Reactome libraries. Scale bar represents the enrichment score of genes within a pathway. **(B)** Diagram showing the increased exposure of N_av_1.6 channels along the nodes of Ranvier in response to AngII-induced HTN, reflecting a disruption of nodal structure. **(C)** Representative IF confocal images and Imaris 3D reconstruction of the colocalization of N_av_1.6 channels (magenta) with Caspr (yellow) in layers 4/5 of the cortex. Scale bar = 20 µm. **(D)** Combined violin and jitter plots depicting the mean volume of Caspr segments and N_av_1.6 nodes. **(E)** Dot plot showing the ratio of Caspr and N_av_1.6 mean volumes. *p*<0.05. Data represents mean ± SEM. Statistical analysis performed using the Wilcoxon test, AngII versus saline. **(F)** Left: Schematic outlining the design of the trans-callosal communication recording protocol. Right: Dot plot depicting the delay between an evoked stimulus and recorded field response, represented by peak latency, as a function of the inter-electrode distance in the corpus callosum. *p*<0.05; n = 4 brains/group. Data represent mean ± SEM. Statistical analysis performed using two-way ANOVA with Šidák’s correction.

Considering that OPC are involved in the process of myelination, including axonal ensheathment and the functional organization of the axon^77^, we next sought to determine whether these transcriptional changes in oligodendrocytes translate into a functional disruption of axons. Since adult born OPC in P60 mouse brains have a cell division cycle between 10 and 30 days^78^, any ensuing effects of the early alterations to OPC function should be evident at the 42D timepoint. The integrity of the node of Ranvier was tested *ex vivo* after 42D of AngII HTN using immunofluorescence to quantify the spatial relationship of voltage-gated sodium channels (N_av_1.6), which are densely clustered within the unmyelinated segment between paranodes, and the paranodal protein Caspr, which interacts with Contactin1 to anchor the outer loops of the myelin sheath to the axonal membrane^77^ (fig. 6B). AngII HTN diminished Caspr volume along axonal tracts, resulting in looser association with Na_v_1.6 and disruption of the overall nodal structure (Fig. 6C-E). These nodal aberrations were associated with an increased latency in transcallosal impulse conduction in brain slices after 42D, but not 3D, of AngII HTN (Fig. 6F), as observed in other models of axonal disruption^79,80^. Therefore, AngII HTN disrupts nodal structure and impairs transcallosal conduction, consistent with the transcriptomic evidence of altered OPC maturation and myelination.

### NEURONAL TRANSCRIPTOMICS SUGGESTS NETWORK DYSFUNCTION EARLY IN ANGII HTN

Unbiased clustering of 13,080 neurons revealed 10 subtypes (Fig. 7A, Fig. S3A), classified as inhibitory or excitatory based on the expression of *Slc17a7* and *Gad1/2*, respectively (Fig. S3B). Integration and transfer of cell-type labels from the Saunders neuronal database aided the identification of equivalent neuronal classes across datasets^42^ (Fig. S3A-C). In interneurons the number of DEG was greatest after 3D of AngII and reduced at 42D, while in excitatory neurons DEG were comparable between 3D and 42D (Fig. 7B).

**Figure 7.**
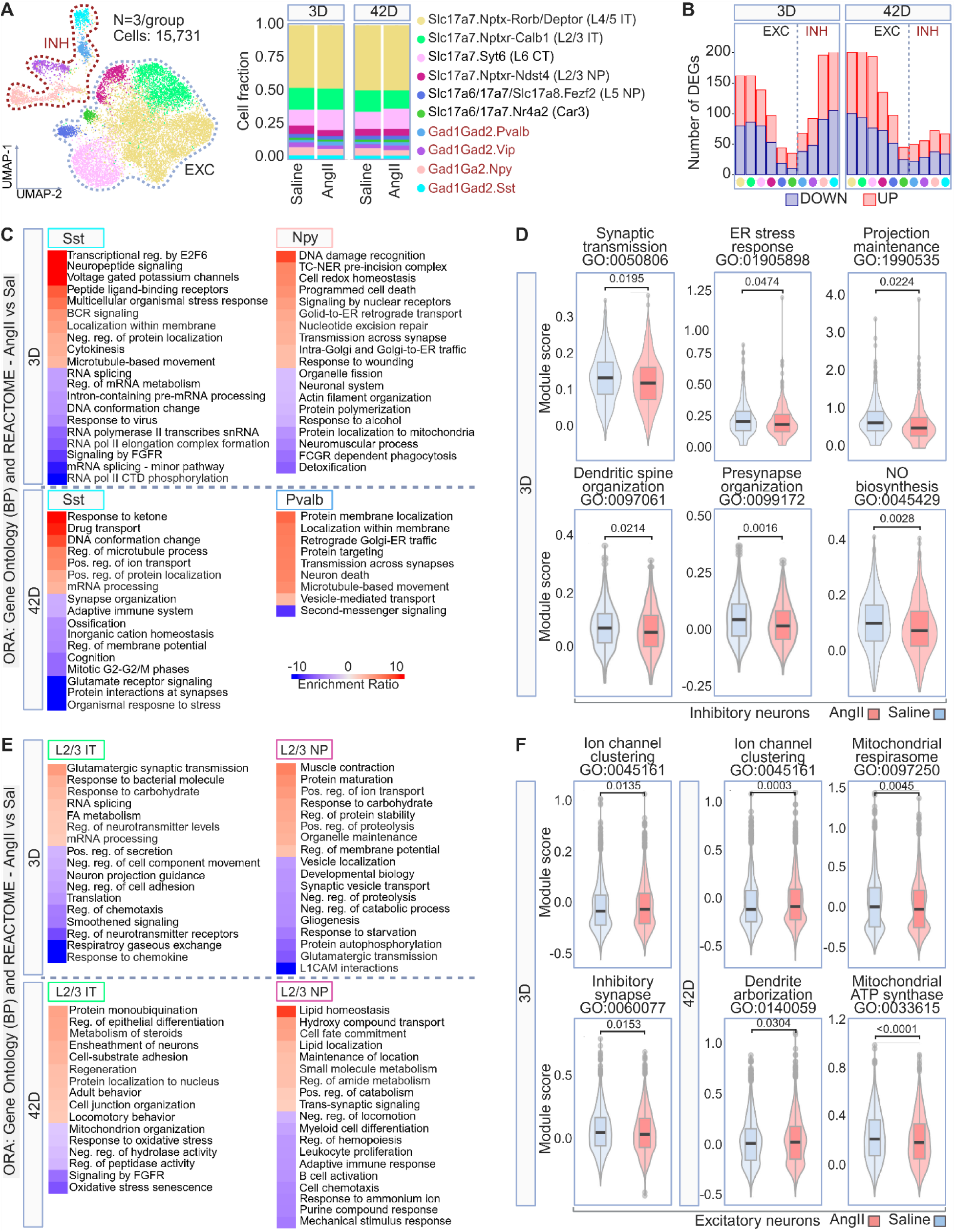
Neurons are affected early in AngII HTN. **(A)** Left: UMAP analysis of 15,731 neurons identified 4 inhibitory interneuron subtypes and 6 excitatory neuron subtypes in merged single cell transcriptomes: Sst (Gad1Gad2.Sst), Npy (Gad1Gad2.Npy), Vip (Gad1Gad2.Vip), Pvalb (Gad1Gad2.Pvalb), L4/5 IT (Slc17a7.Nptxr-Rorb/Deptor), L2/3 IT (Slc17a7.Nptxr-Calb1), L6 CT (Slc17a7.Syt6), L2/3 NP (Slc17a7/Nptxr-Ndst4), L5 NP (Slc17a6/17a7/17a8.Fezf2), Car3 (Slc17a6/17a7.Nr4a2). Right: Bar graph showing the relative frequencies within each sub-cluster across time points and treatments. Also see Figure S3. **(B)** Bar graph depicting the number of significant upregulated and downregulated differentially expressed genes (DEG) in each subtype in response to 3 days or 42 days of AngII. **(C)** Heatmaps representing AngII-induced, time-point specific enrichment of biological pathways within inhibitory neuronal subtypes, determined through overrepresentation analysis (ORA) of upregulated and downregulated DEG (*p*<0.05, logFC>2). Pathways are derived from the Gene Ontology Biological Process and Reactome libraries. Scale bar represents the enrichment score of genes within a given pathway. **(D)** Violin and box plots showing the results of module score analyses for select gene sets in inhibitory neurons after 3 days of AngII administration. *p*<0.05. Boxplots represent the median ± IQR. Statistical analysis performed using the Wilcoxon test, AngII versus saline. **(E)** Heatmaps representing the time-point specific enrichment in biological pathways induced by AngII treatment in excitatory neuronal subtypes, determined through ORA of upregulated and downregulated DEG (*p*<0.05, logFC>2). Pathways derived from the Gene Ontology Biological Process and Reactome libraries. Scale bar represents the enrichment score of genes within a given pathway. **(F)** Violin and box plots showing the results of module score analyses for select gene sets after 3 days (left) or 42 days (right) of AngII treatment in excitatory neurons. *p*<0.05. Boxplots represent the median ± IQR. Statistical analysis performed using the Wilcoxon test, AngII versus saline.

In interneurons, changes were most pronounced in the Gad1Gad2.Npy and Gad1Gad2.Sst clusters, reflecting *Npy* and *Sst* neurons, respectively (Fig. 7A). After 3D, AngII induced the upregulation of genes involved in redox and potassium homeostasis (*Ndufb7, Ndufab1, Atp1a3, Kcnb1, Kcna6, Kcnab1)* and cellular stress responses (*Mapk8ip3*, *Ppp3r1, Serp2, Nae1, Os9)* in Npy and Sst interneurons (Fig. 7C). Sst interneurons also exhibited a strong downregulation of genes involved in the synaptic vesicle cycle (*Stxbp1, Atp6v0b, Gdi1, Cltb, Pacsin1, Bcl11a, Dhx36, Vps4a, Wasl, Rdx*) (Fig. 7C), reflecting impairments in endosome processing, vesicle docking, activity-dependent bulk endocytosis, vesicle recycling, and synapse organization^81,82^. Subsequent module scoring confirmed that genes involved in synaptic transmission, neuron projection maintenance, dendritic spine organization, and pre-synaptic organization were downregulated in inhibitory interneurons (Fig. 7D), reinforcing their early impairment in AngII HTN.

Conversely, in excitatory neurons, genes involved in synaptic function were upregulated after 3D of AngII HTN, as anticipated from the diminution of inhibitory influence^83,84^. Relevant pathways included positive regulation of ion transport and regulation of membrane potential in L2/3 NP (*Tesc, Rab3gap, Akap9, Cck, Fgf13*), glutamatergic synapse and regulation of neurotransmitter levels in L2/3 IT (*Unc13c, Unc13b, Ptgs2, Ncs1*), exocytosis and regulation of intracellular transport in L6 CT (*C2cd5, Rap1b, Nlgn1, Nefh, Arhgef2, Rab27b, Bloc1s6*), and synapse organization in Car3 (*Ntrk2, Myo5a, Atp2b2, Nefl, Rock2, Ptprt, Nefm*) (Fig. 7E). Subsequent module scoring revealed an upregulation in genes involved in ion channel clustering, an important determinant of neuronal activity^85,86^. Supporting a reduction in the association of excitatory and inhibitory neurons, module scores for inhibitory and excitatory synapse organization were diminished in excitatory and inhibitory neurons, respectively (Fig. 7F). Taken together, these data indicate an early and sustained disruption of synaptic function in interneurons concurrent with an enhancement of synaptic strength in excitatory neurons, suggesting an excitation-inhibition imbalance.

### EVIDENCE OF NEURONAL HYPERACTIVITY AND METABOLIC DYSFUNCTION EMERGES LATE IN ANGII HTN

After 42D, AngII induced a downregulation of genes involved in protein-protein synaptic interactions (*Gria1, Grm5, Dlgap1, Sparcl1, Mef2c, Tac1*) in Sst interneurons, with the reduction of *Gria1* and *Grm5* suggesting that excitatory contacts remained suppressed (Fig. 7C). Conversely, genes pertaining to synapse organization (*Ephb3, Slit1, Sez6, Tsc1, Lingo2, Dag1, eIF4a1*) and the regulation of synaptic structure (*Dhx36, Ptprt, Dnm3, Atp2b2, Mef2c, Kcnc1*) were elevated in L2/3 IT and Car3 excitatory neurons (Fig. 7E), respectively. Supervised module scoring confirmed that gene-sets concerning synaptic function, including ion channel clustering, dendritic arborization, and AMPA receptor clustering, were increased (Fig. 7F). Notably, this transcriptomic signature, suggestive of increased excitatory neuron activity, coincided with a marked downregulation in genes pertinent to mitochondrial respiration, including those related to mitochondrial function (*Cox14, Romo1, Fmc1, Tmem70, Fam162a, Ssbp1, Dnaja3, Ucqcr10, Ndufa3, Ndufa8, Fis1, Ywhaz, Fbxw7, Mrpl17*) in L5 NP and L2/3 IT, the citric acid cycle and electron transport chain (*Ndufa3, Ndufa8, Ldha, Sdhb, Atp5b, Atyp5j*) in L5 NP and Car3, and precursor metabolite and energy generation (*Uqcrq, Sdhb, Mt3, Cox7a2l*) in Car3 (Fig. 7E). Subsequent module scoring confirmed a focal decrease in genes involved in both mitochondrial respiration and mitochondrial ATP synthase complex function (Fig. 7F). Since elevated synaptic activity requires a corresponding increase in mitochondrial ATP^87^, the discrepancy between heightened synaptic activity and reduced energy production indicated by the transcriptomic data suggests a potentially harmful state where metabolic demands exceed the availability of energy substrates.

### EARLY SUPPRESSION OF GABA TRANSPORTER AND NEURONAL NITRIC OXIDE PRODUCTION

Cortical interneurons expressing the neuronal isoform of NO synthase (*Nos1* or nNOS) have close contacts with cerebral microvessels^88^ and contribute to neurovascular coupling by releasing the potent vasodilator NO^89–91^, a response suppressed in AngII HTN^34^. Recent transcriptomic studies suggest that nNOS is most abundant in a well-defined subset of interneurons which co-express *Sst, Npy, and Chodl*^92,93^. (Fig. S3A-B). ORA revealed suppression of genes related to nitrogen compound catabolism in the Sst cluster after 3D of AngII HTN (*Pde1a, Cnot7, Dhx9, Lrpprc, Ncbp2, Elavl4, Lsm4, Dhx36*) (Table S1). Subsequent supervised module scoring confirmed a concurrent reduction in the expression of genes involved in NO biosynthesis (Fig. 7D), suggesting that NO production is reduced in the Sst cluster alongside the attendant decrease in the catabolism of nitrogenous precursors. In addition, the expression of *Slc32a1*, which is transcriptionally regulated by NO signaling and encodes the presynaptic vesicular GABA transporter (VGAT)^94^, was diminished in the Sst and Npy subclusters (Fig. 8A).

**Figure 8.**
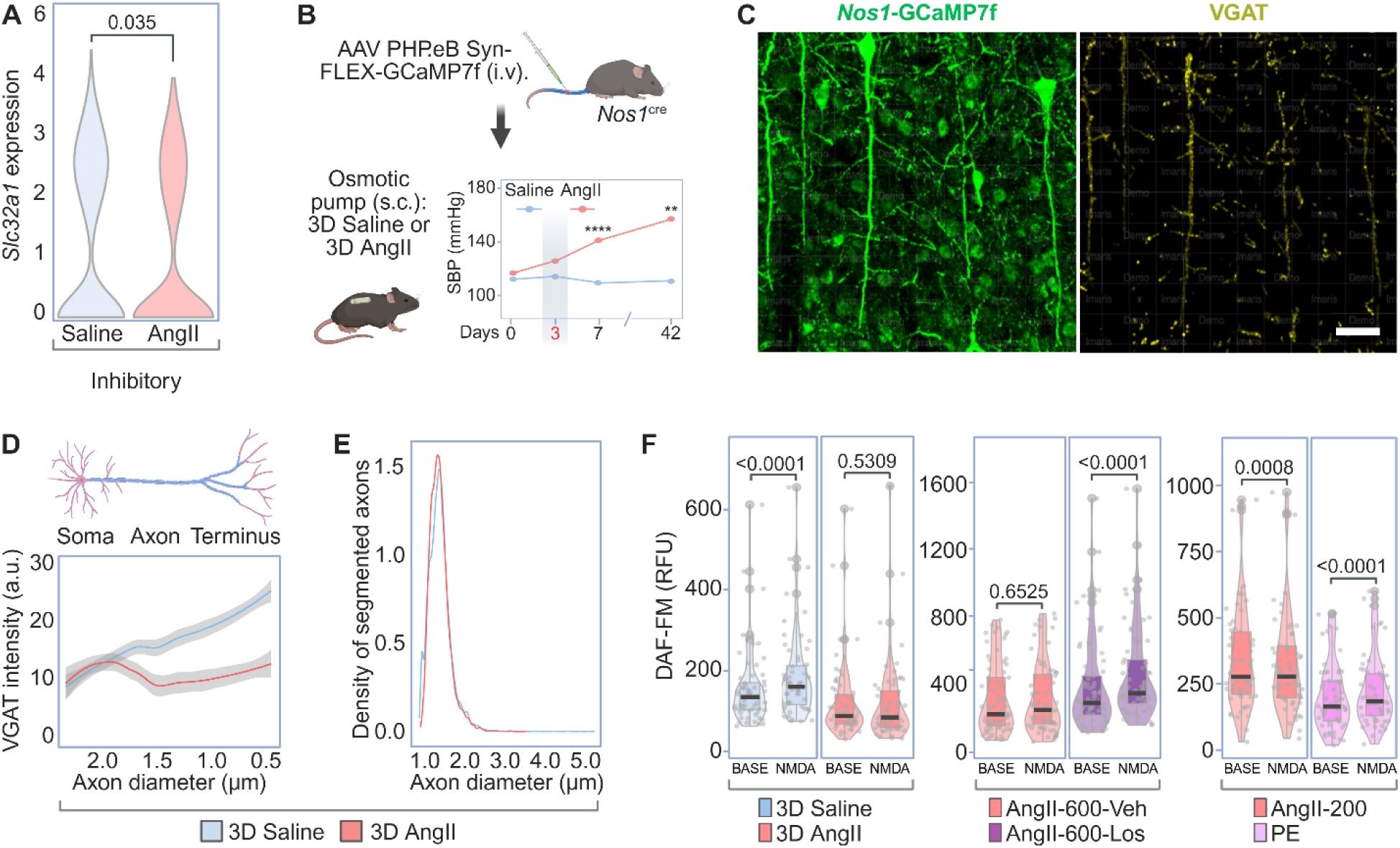
AngII elicits GABAergic deficits and suppression of NO production in interneurons. **(A)** Violin plot showing the expression of *Slc32a1*, which encodes the vesicular GABA transporter (VGAT). *p*<0.05. Statistical analysis performed using the Wilcoxon test, AngII versus saline. **(B)** Top: Diagram representing the generation of Nos1^Cre^-GCaMP7f reporter mice. Bottom: Nos1^Cre^-GCaMP7f mice were treated with AngII or saline for 3 days. **(C)** Representative IF confocal images of VGAT (yellow) in *Nos1*^Cre^ mice expressing Ca^2+^-dependent GCaMP7f (green). Left: IF images showing that the GCaMP7f AAV transduced nNOS neurons with high efficiency. Right: Imaris reconstructed image of the VGAT signal associated with GCaMP7f^+^ neurons. Scale bar = 30 µm. **(D)** Quantification of VGAT signal intensity corresponding to axonal diameter. n = 4 brains/group. Data represent mean ± CI. **(E)** Density plot depicting the quantification of axon segment counts ordered by axon diameter; data generated using Imaris software. n = 4 brains/group. **(F)** Combined violin, box, and jitter plots depicting NO production (captured through DAF-FM fluorescence) at baseline (left) and in response to NMDA (right) in dissociated cortical neurons isolated from mice treated with saline, or slow-pressor AngII, slow-pressor AngII, slow-pressor AngII followed by AngII + losartan, non-pressor AngII, or phenylephrine (PE). *N* = 60 cells/group, 4 mice/group. Statistical analysis performed using a paired, two-tailed t-test. Boxplots represent the median ± IQR. Dots represent individual cells.

To corroborate these findings, we treated Nos1-GFP reporter mice with saline or AngII for 3D to investigate VGAT expression (Fig. 8B-C). VGAT expression was diminished in cortical layers V/VI, particularly in axonal segments approaching 0.5 µm in diameter (Fig. 8D), suggesting early suppression of GABAergic release probability^95^. The number of axon terminals was unaffected (Fig. 8E), implying that AngII influences interneuronal function without synaptic loss. Next, in wild-type mice, we used the NO indicator DAF-FM to assess activity-dependent NO production. In dissociated cortical neurons, we found that 3D of AngII HTN induced a marked reduction in both baseline and NMDA-evoked NO signals (Fig. 8F). Attesting to the involvement of AT1R signaling and not of the BP elevation, the suppression of neuronal NO production was reversed by Losartan, was still present with non-pressor doses of AngII and was not observed if HTN was induced by the vasoconstrictor phenylephrine (Fig. 8F). These findings support our transcriptional data indicating that interneuronal dysfunction leads to attenuation in activity-induced NO production early in AngII HTN and provide the molecular bases of the ensuing attenuation in NO-dependent neurovascular coupling induced by AngII HTN.

### TRANSCRIPTOMIC CHANGES IN MURAL CELLS

Unbiased clustering of 2,282 mural cells revealed two primary subtypes, SMC and PC, which were characterized by expression of *Tagln* and *Acta2* or *Cspg4* and *Pdgfrb*, respectively (Fig. S4A-B). Accounting for cell proportion, the number of transcriptional alterations induced by AngII were roughly equivalent between time points and cell types (Fig. S4C).

In SMC, genes involved in cytoskeletal structure, contraction, and proliferation were both upregulated (*Coro1b, Nme1, Bcas3, Syt1, Pfn1, Vtn, Hspa1a, Snx3*) and downregulated (*Myom1, Kcnab1, Luc71, Akap6, Smarca2, Arp5c, Taok1, Rock1*) after 42 days of treatment with AngII, indicating an alteration to their contractile phenotype. Similar changes were observed after 3D, albeit to a lesser extent (*Msn, Map4, Arp5c, Cnn2, Dstn*) (Fig. S1D). In PC, 3D of AngII HTN upregulated genes pertaining to actomyosin structure organization (*Myo18a, Slmap, Tpm1, Clasp2*) and exocytosis (*Snapin, Kcnb1, Clasp2, Rala, Cask*) (Fig. S4E). Since PC contribute to the BBB maintenance^96^ and may regulate capillary flow^97^, these changes could contribute to the AngII induced BBB alterations and reduced cerebral perfusion.

### TRANSCRIPTOMIC CHANGES IN ASTROCYTES AND MICROGLIA

Unbiased clustering of 2,129 astrocytes and 1,010 microglia did not reveal any subtypes of interest (Fig. S5A-B; Fig. S6A-B). While AngII-induced transcriptional alterations were present (Fig. S5C; Fig. S6C), they were limited in comparison to other cell types in the overall dataset, and functionally relevant changes were sparse in our pathway analyses, supervised module scoring, and manual explorations.

In astrocytes, a small number of genes associated with neuropeptide signaling (*Npy*)^98^ and “gliotransmission” (*Phgdh, Grm3*)^99,100^ were upregulated after 3D of AngII HTN, while those involved in astrocytic iron uptake and transport, a potentially important process in brain iron trafficking, were reduced (*Unc, Atp6v1f, Trfc, Atp6v1g1*)^101^ (Fig. S5D). At 42D, a variety of synaptic genes were upregulated (*Cttnbp2, Dbn1, Rab3a, Mesd, Crkl, Pclo, Arf4, Plppr4, Golga4, Stmn1, Mycbp2, Apod*) (Fig. S5D), including *Apod*, the product of which, Apolipoprotein D, is secreted from astrocytes and functions to improve neuronal survival following oxidative stress^102,103^.

In microglia, at 3D AngII HTN induced a transcriptional signature suggestive of a phagocytic phenotype, albeit limited, which included genes involved in proteolysis (*Rbx1, Akirin2, Asph, Ctsc*), cellular component disassembly (*Smarcc2, Stmn2, Asph, Chchd10*), and the regulation of peptidase activity (*Akirin2, Birc6, Asph*) (Fig. S6D)^104^. Also, a small number of genes pertaining to cytokine production were upregulated (*Tnfrsf13b, Rbx1, Ifnar2*) (Fig. S6D), but not to an extent sufficient to indicate any sort of pronounced inflammatory activation. No notable alterations were observed at 42D.

## DISCUSSION

We utilized scRNA-seq to investigate the cell type-specific alterations elicited by AngII-induced slow-pressor HTN in mice, a model which reproduces key features of the effects of human HTN on the brain^9,21,22^. Several novel findings were revealed. First, major transcriptomic alterations by AngII are observed in endothelial cells, neurons, and oligodendrocyte-lineage cells, emerging after just 3D of treatment and preceding the development of neurovascular impairment. Second, the capillary-venular zone is the primary site of the early pathological effects of AngII HTN on the endothelium. In particular, we found evidence of endothelial senescence and of a shift from specific receptor-mediated transport to non-specific transcytosis, as seen in the aging cerebral endothelium^49,105^. The premature endothelial aging phenotype was corroborated by well-established markers of senescence^106^. HTN-induced pathology was also observed in arteriolar cells at 42D, when endothelial cells exhibited transcriptomic changes suggestive of remodeling, verified by collagen IV immunohistochemistry. Third, oligodendrocyte-lineage cells were affected early in AngII HTN, exhibiting transcriptomic signatures indicative of increased stress responses, reduced proliferation, arrested differentiation and DNA damage in OPC^65–68^, as well as disruption of myelin-associated processes in Pre-OL and MOL^71^, which were still evident at 42D. The structural and functional impact of these OPC and myelination-related changes were validated by demonstrating DNA damage in OPC and by disruption of the nodes of Ranvier resulting in slowing transcallosal impulse conduction. Fourth, contrasting effects on synaptic function-associated genes in inhibitory (downregulation) and excitatory (upregulation) neurons emerged after just 3D of AngII HTN, suggesting a shift in the excitation-inhibition balance toward an elevated excitatory tone. These transcriptomic signatures are still evident at 42D, concurrent with a downregulation of genes associated with mitochondrial function in excitatory neurons indicative of metabolic stress. Interneuronal hypofunction was confirmed *in situ* through a reduction in the GABA transporter VGAT at 3D of AngII and dampening of resting and activity-induced NO production by nNOS-expressing interneurons. Fifth, consistent with the remodeling signature observed in the arteriolar endothelium, at 42D of AngII SMC exhibited expression of genes associated with contractility and cytoskeletal structure.

These neuronal and vascular changes provide a first insight into the molecular bases of the alterations in neurovascular coupling, BBB permeability, neural network dysfunction and white matter damage observed in HTN, both in humans^18,19,28–30^ and in animal models^13,14,107^. The minimal overlap of DEGs between 3D and 42D is noteworthy, as it suggests that distinct transcriptomic programs are activated before and after the onset of neurovascular and cognitive impairments. However, a key question concerns how these changes are initiated. In slow pressor HTN, circulating AngII, in addition to activating AT1R in the subfornical organ and initiating the neurohumoral dysfunction that drives the elevation in blood pressure, engages AT1R on the cerebral endothelium resulting in increased BBB permeability^13,14^. AngII crosses the BBB but remains confined to the perivascular space, wherein it activates AT1R on perivascular macrophages (PVM) leading to increased ROS production and neurovascular dysfunction^13^. Our data support this scenario by showing molecular evidence of increased endothelial transcytosis and oxidative stress in multiple cell types. Unexpectedly, however, these molecular changes occur much earlier than anticipated since they are observed after just 3D of AngII administration, while BP elevation, neurovascular uncoupling and BBB dysfunction are first seen at 7-14 days^13,108^ and cognitive impairment at 42 days^31^. These observations suggest that the transcriptomic and functional changes in neurovascular cells, oligodendrocytes and neurons, are independent of these neurovascular alterations. Therefore, the data reveal major molecular changes in NVU cells that may set the stage for the later development of neurovascular and cognitive dysfunction.

Another key pathogenic event revealed by our analysis is the early development of cerebral endothelial senescence. It is well established that AngII HTN induces senescence in systemic endothelial cells^109,110^, but it remained unclear whether premature aging also occurs in cerebral endothelial cells. Rather, previous studies failed to provide evidence of senescence in whole brain preparations in slow-pressor AngII hypertension^111^, or in isolated brain endothelial cells in a model of genetic hypertension dependent on AngII (BPH mouse)^112^. In contrast, our scRNA-seq analysis demonstrates for the first time a profound senescent phenotype highly restricted to the capillary-venular zone, which could explain why it was previously missed in whole brain analyses. The mechanisms by which AngII HTN induces senescence in a specific microvascular segment remain to be determined. One possibility is that senescence is triggered directly from activation of AT1R via signaling leading to telomere shortening, insufficient endothelial cell renewal, and inflammation, as described in systemic endothelial cells^113,114^. AngII-generated ROS could also induce senescence^110^. However, the observation that premature aging is most prominent in the capillary-venular zone and not in larger microvessels in contact with PVM, the major source of AngII-induced ROS^13^, make this possibility less likely. It is also unlikely that the venular effects are due to increased AT1R expression, since these receptors are not particularly enriched in venules^115,16,39^. However, cerebral venules are enriched in inflammatory mediators which could play a role^8^. Whatever the mechanisms of the zonation of senescence, it is conceivable that the localization to the endothelial cells of capillaries and venules indicates that senescence is involved in the BBB dysfunction of AngII HTN^14^, as reported in the aging mouse brain^116^.

Damage to the white matter is a hallmark of the effects of HTN on the brain and the major cause of HTN-induced cognitive impairment^18–20^. Early transcriptomic changes were observed in oligodendrocytes, indicating stress responses, arrested development in OPC and altered myelination-related process in Pre-OL and MOL. DNA damage in OPC was confirmed with γ-H2A.X. These alterations, which precede vascular dysregulation, provide insights into the early pathogenic events driving the white matter damage induced by HTN, which mainly targets myelinated axons^117^. Given that these effects include suppression of OPC differentiation^65^, on which white matter integrity critically depends^77^, it is likely that early disruption of OPC recruitment and differentiation into mature oligodendrocytes are a contributing factor in the altered nodal architecture observed in this study. In support of this hypothesis, inhibition of oligodendrocyte differentiation has been shown to exacerbate white matter damage and memory impairment, in murine models of AD^118–120^ and disruption of nodal architecture is associated with cognitive impairment in white matter hypoperfusion^121^. The cognitive impact of these transcriptomic changes is observed several weeks later (at 42D), coincident with alterations in nodal structure and axonal conduction. This temporal profile is consistent with human data showing subtle white matter alterations in otherwise healthy young adults^122,123^, which worsen over time and lead to overt white matter disease and cognitive impairment later in life^124–126^.

Early transcriptomic changes were also observed in inhibitory and excitatory neurons, reflecting a shift from inhibitory to excitatory tone. Interneurons are in close association with the cerebral vasculature^127^ and are the largest source of NO during neurovascular coupling^128^. In the present study, transcriptomic and *in situ* evidence of interneuronal hypofunction coincided with suppression of NMDA-induced NO production, raising the possibility that the progressive neurovascular dysfunction developing in AngII HTN^108^ is caused, in part, by the attenuation of NO release from interneurons during neural activity. Concomitant with the interneurons hypofunction, the data also revealed a previously-unappreciated increased in the activity of excitatory neurons which may underlie the excitation/inhibition imbalance and propensity to develop seizures in patients with hypertension^129–131^. In addition, the later metabolic stress observed in neurons at 42D may reflect the longer-term effects of the mismatch between the increased energy requirements imposed by neural activity and the reduced delivery of blood flow.

The mechanisms by which HTN induced alteration in neurons and oligodendrocytes, which, unlike the endothelium, are further away from circulating AngII remain to be elucidated. One possibility is that senescent endothelial cells can adversely affect their neighbors through the senescence-associated secretory phenotype (SASP)^132–134^. The SASP is characterized by pro-inflammatory cytokines, cell cycle regulators, and pro-angiogenic factors, which induces cell dysfunction and premature aging in neighboring cells via autocrine and paracrine signaling^132–134^. Therefore, endothelial SASP could propagate the dysfunction beyond the endothelium targeting interneurons and oligodendrocytes. Considering that the neurovascular dysfunction, BBB disruption and cognitive impairment induced by AngII HTN are reversed by suppressing ROS generation in PVM^13,14^, the role of endothelial senescence in the oligodendrocyte disruption and neuronal dysfunction require additional investigations. Direct effects of AngII on neurons and OPC are unlikely since in this model the BBB is still intact at 3D^13^. Furthermore, although AngII crosses the BBB at 14D, it remains confined to the perivascular space and does not penetrate the brain parenchyma^13^. However, neuronal and glial effects of the activation of central autonomic pathways, as well as hormonal release, induced by circulating AngII through AT1R in the subfornical organ cannot be ruled out.

How applicable are findings from a rodent model based on increasing circulating AngII to human HTN? Although circulating AngII levels are elevated in renovascular or malignant HTN, they are typically normal in the more common essential HTN^21,135^. Nevertheless, AngII plays a central role in the pathophysiology of human HTN, as evidenced by the widespread use and effectiveness of renin-angiotensin system inhibitors as first-line antihypertensive agents^136^. Assuming that AngII signaling mechanisms are conserved between rodents and humans, the molecular changes induced by AngII in rodents may also be relevant to essential HTN—though this warrants further investigation. Future studies should extend these finding to aging mice, including postmenopausal females, who are markedly affected by the adverse effects of HTN^137^. Still, our current study in young mice remains highly relevant, given the rising prevalence of HTN in young adults, where it has been linked to white matter alterations^123^.

In conclusion, we performed the first scRNA-seq study of the effect of AngII HTN on cells of the NVU at the microvascular level. We found significant transcriptomic changes affecting endothelial cell, oligodendrocytes, neurons and smooth muscle cells that preceded the development of neurovascular coupling dysfunction, BBB leakage and cognitive impairment. Through a combination of transcriptomic investigation and *in situ* histological and functional validation, we started to delineate the most prominent functional pathways impacted in AngII HTN and their potential link to neurovascular and cognitive dysfunction. These studies represent the prelude to further mechanistic investigations to establish precisely how these cooperative transcriptomic changes contribute to the development of neurovascular dysfunction, white matter damage, and cognitive impairment in HTN. To this end, we have established an online platform to provide a user-friendly resource for further mechanistic studies and exploration of potential therapeutic targets (https://anratherlab.shinyapps.io/angii_brain/).

## Supporting information

Key Resources Table

## RESOURCE AVAILABILITY

### LEAD CONTACT

Requests for further information and resources should be directed to the lead contact, Costantino Iadecola, MD.

### MATERIAL AVAILABILITY

This study did not generate new unique reagents.

### DATA AND CODE AVAILABILITY

This paper does not report original code. All data generated by quantification of immunofluorescence/histology/behavior is provided in the supplementary materials accompanying this manuscript. Microscopy data reported in this paper will be shared by the lead contact upon request. Single-cell RNA-seq data have been deposited to GEO at GSE294795 and are publicly available as of the date of publication. Any additional information required to reanalyze the data reported is available upon request.

## ACKNOWLEDGEMENTS

This work was supported by NIH grants R01-NS095441(CI), R01NS081179 (J.A.) and by Cure Alzheimer Fund grant 243348 (C.I., G.F.). Support from the Feil Family Foundation is gratefully acknowledged.

## AUTHOR CONTRIBUTIONS

A.G.P., S.M.S., S.J.A., and M.M.S., conducted the experiments; S.M.S. and A.G.P. performed the data analysis; G.W. measured NO production and transcallosal conduction; G.R. performed the qRT-PCR; C.I., J.A., L.P., and G.F. supervised the research; S.M.S., A.G.P., CI, and J.A. wrote the manuscript; C.I., J.A., and G.F. provided funding.

---

**Figure S1.**
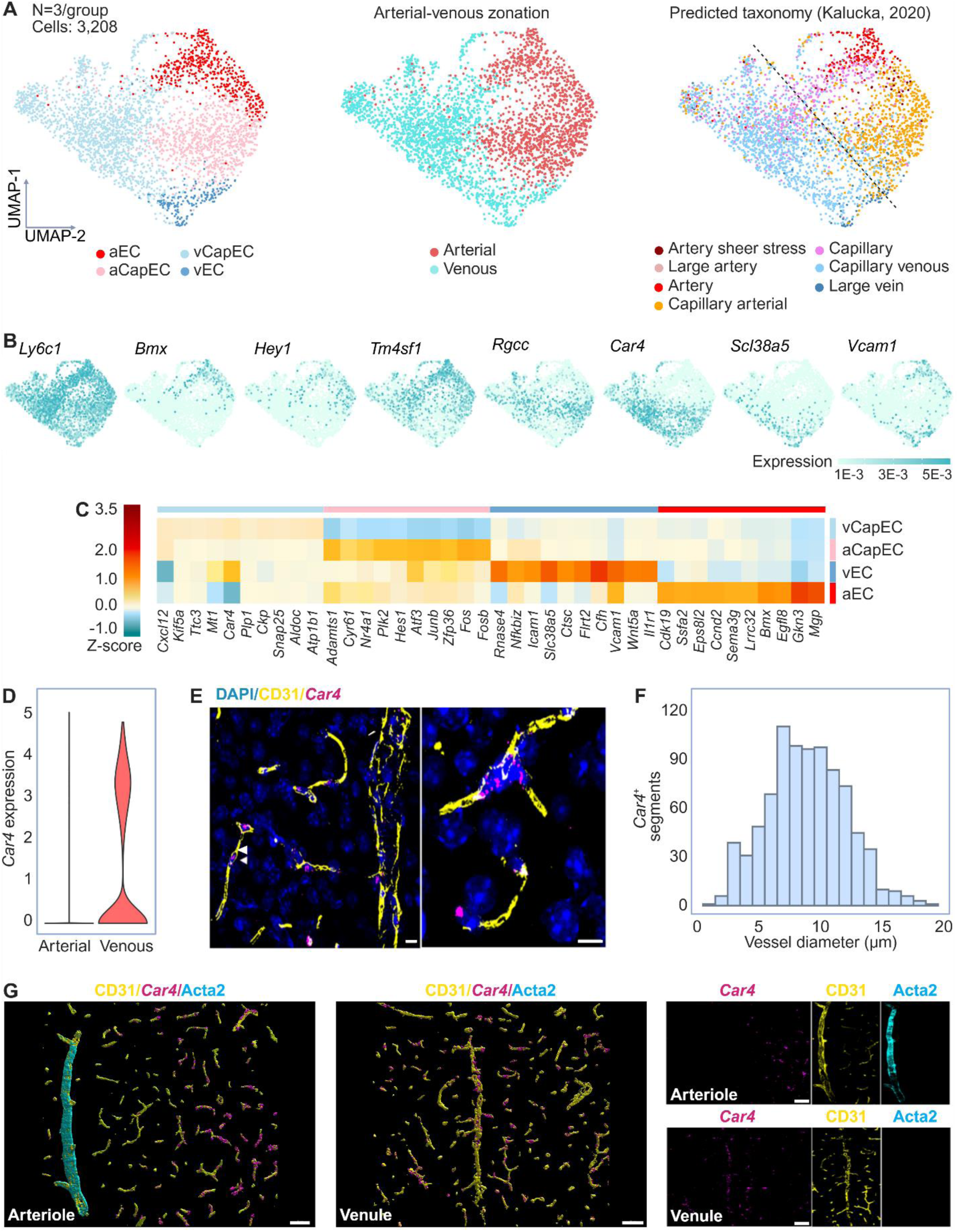
Endothelial cell subtypes align along an arterial-venous axis, related to Figures 2 and 4. **(A)** Left: UMAP analysis of 3,232 endothelial cells depicting the 4 subtypes identified in merged single cell transcriptomes: arteriolar (aEC), arteriolar capillary (aCapEC), venular (vEC), and venular capillary (vCapEC). Middle: UMAP classification of endothelial cells into arteriolar and venular zones. Right: UMAP representation of the unsupervised integration and transfer of cell-type labels from the Kalucka (2020) brain endothelial cell scRNAseq database^47^. **(B)** From left to right, feature plots showing the expression of marker genes characterizing the molecular transition along the arterial-venous axis. Scale bar represents log normalized gene expression. **(C)** Heatmap displaying the expression of the top ten upregulated genes in each sub-cluster. Scale bar represents z-score of the average log gene expression. **(D)** Violin plot showing the expression of the venular marker, *Car4*, in endothelial cells within the arteriolar and venular zones. **(E)** RNA-FISH images of *Car4* (magenta) expression in the naïve cortex, combined with IF of CD31^+^ endothelial cells and nuclear staining of DAPI. Scale bar = 20 µm. **(F)** Histogram depicting the morphometric analysis of *Car4*^+^ vessel diameter. Bin size of vessel diameters = 5 µm. **(G)** Imaris 3D reconstruction of RNA-FISH images for *Car4* (magenta) in the naïve cortex combined with IF for CD31^+^ endothelial cells (yellow), α-Acta2^+^ smooth muscle cells (cyan), and nuclear staining with DAPI. Images are labelled according to the presence of arterioles or venules (white text). Scale bar = 20 µm.

**Figure S2.**
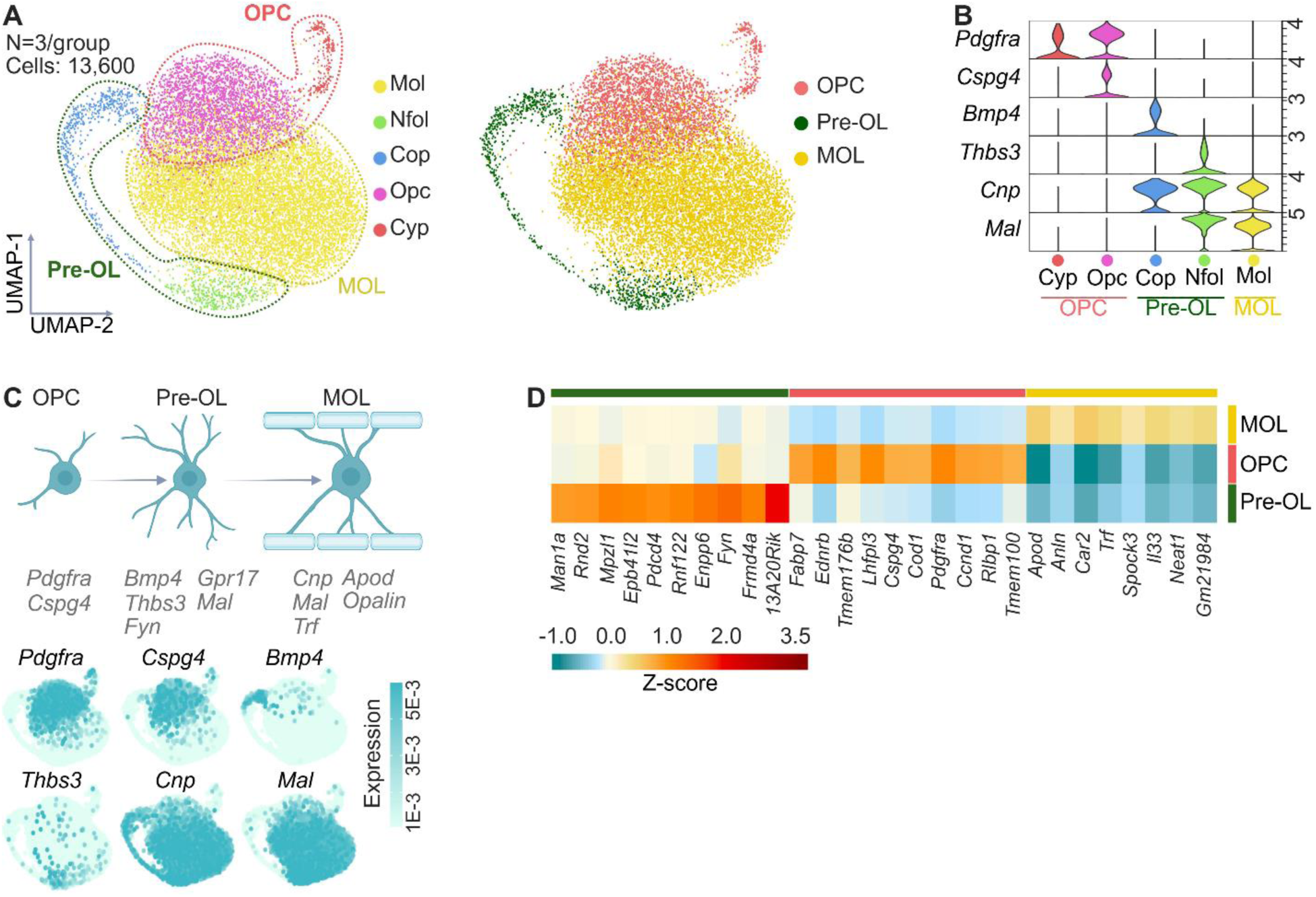
Validation of oligodendrocyte-lineage cell clustering and analysis of cell-cell communication within the oligovascular niche, related to Figures 5 and 6. **(A)** Left: UMAP representation of 13,600 oligodendrocyte-lineage cells showing the 5 subtypes identified in merged single cell transcriptomes: oligodendrocyte precursors (Opc), cycling precursors (Cyp), committed precursors (Cop), newly formed oligodendrocytes (Nfol), and mature oligodendrocytes (Mol). Right: UMAP depicting the organization of oligodendrocyte-lineage cells into 3 subtypes characterized by degree of maturation: oligodendrocyte precursors (OPC), pre-myelinating oligodendrocytes (Pre-OL), and mature oligodendrocytes (MOL). **(B)** Stacked violin plots showing the relative expression of marker genes in each subtype. **(C)** Top: Diagram of genes which characterize the maturation status of oligodendrocyte-lineage cells. Bottom: From top left to bottom right, feature plots depicting the expression of genes characterizing the transition of oligodendrocyte-lineage cells from precursors to mature oligodendrocytes. Scale bar represents log normalized gene expression. **(D)** Heatmap displaying the expression of the top ten upregulated genes in each sub-cluster. Scale bar represents z-score of the average log gene expression.

**Figure S3.**
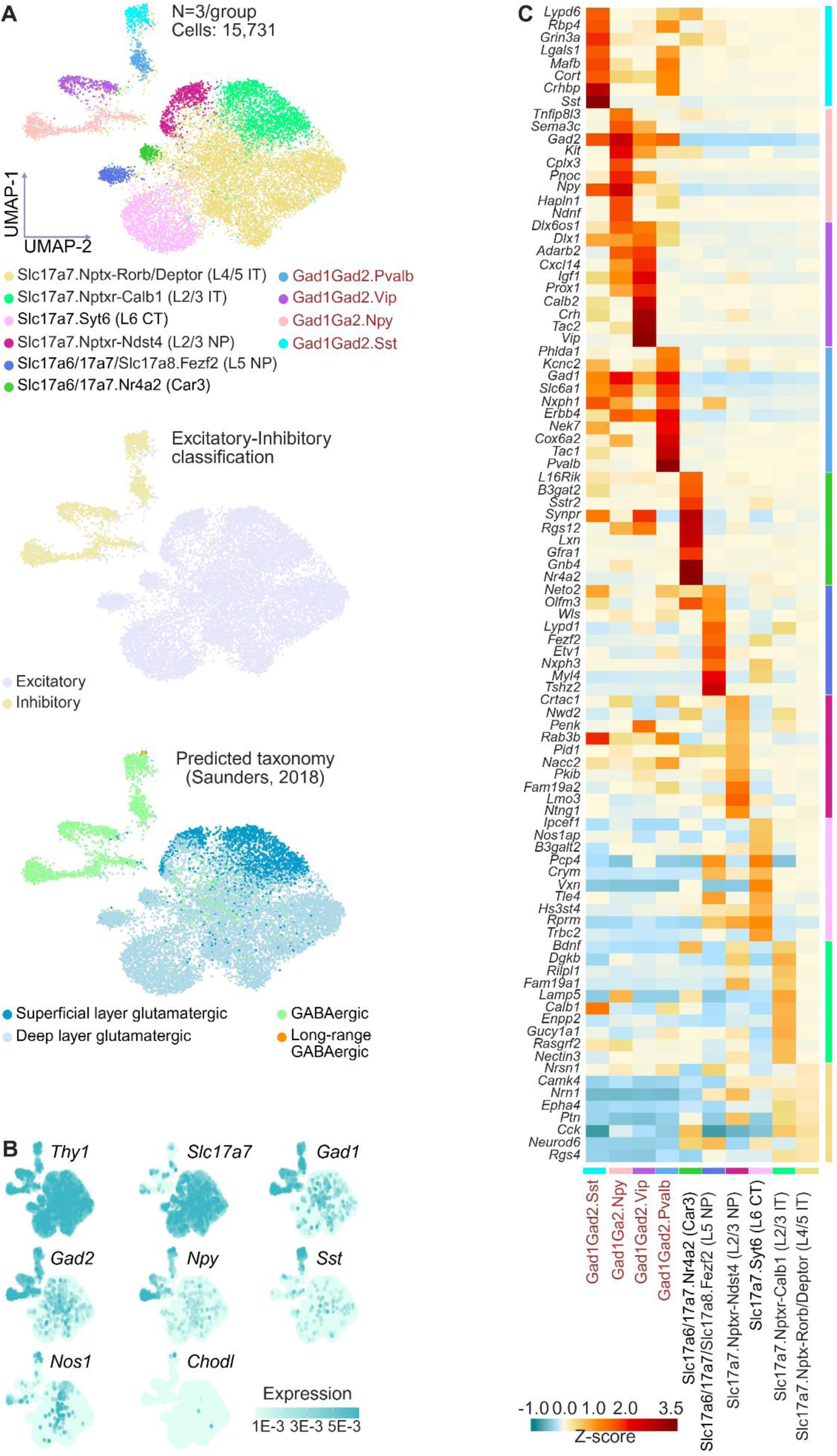
Cortical neurons align along an excitatory-inhibitory axis, related to Figure 7. **(A)** Top: UMAP representation of 15,731 neurons in merged single cell transcriptomes, revealing 4 inhibitory subtypes and 6 excitatory subtypes: Sst (Gad1Gad2.Sst), Npy (Gad1Gad2.Npy), Vip (Gad1Gad2.Vip), Pvalb (Gad1Gad2.Pvalb), L4/5 IT (Slc17a7.Nptxr-Rorb/Deptor), L2/3 IT (Slc17a7.Nptxr-Calb1), L6 CT (Slc17a7.Syt6), L2/3 NP (Slc17a7/Nptxr-Ndst4), L5 NP (Slc17a6/17a7/17a8.Fezf2), and Car3 (Slc17a6/17a7.Nr4a2). Middle: UMAP showing the excitatory-inhibitory classification of neurons. Bottom: UMAP representation of the unsupervised integration and transfer of cell-type labels from the Saunders (2018) cortical neuron scRNAseq database^42^. **(B)** From top left to bottom right, feature plots depicting the expression of genes which characterize excitatory versus inhibitory neurons. The final 2 feature plots highlights markers of nNOS^+^ neurons. Scale bar represents log normalized gene expression. **(C)** Heatmap displaying the expression of the top ten upregulated genes in each sub-cluster. Scale bar represents z-score of the average log normalized gene expression.

**Figure S4.**
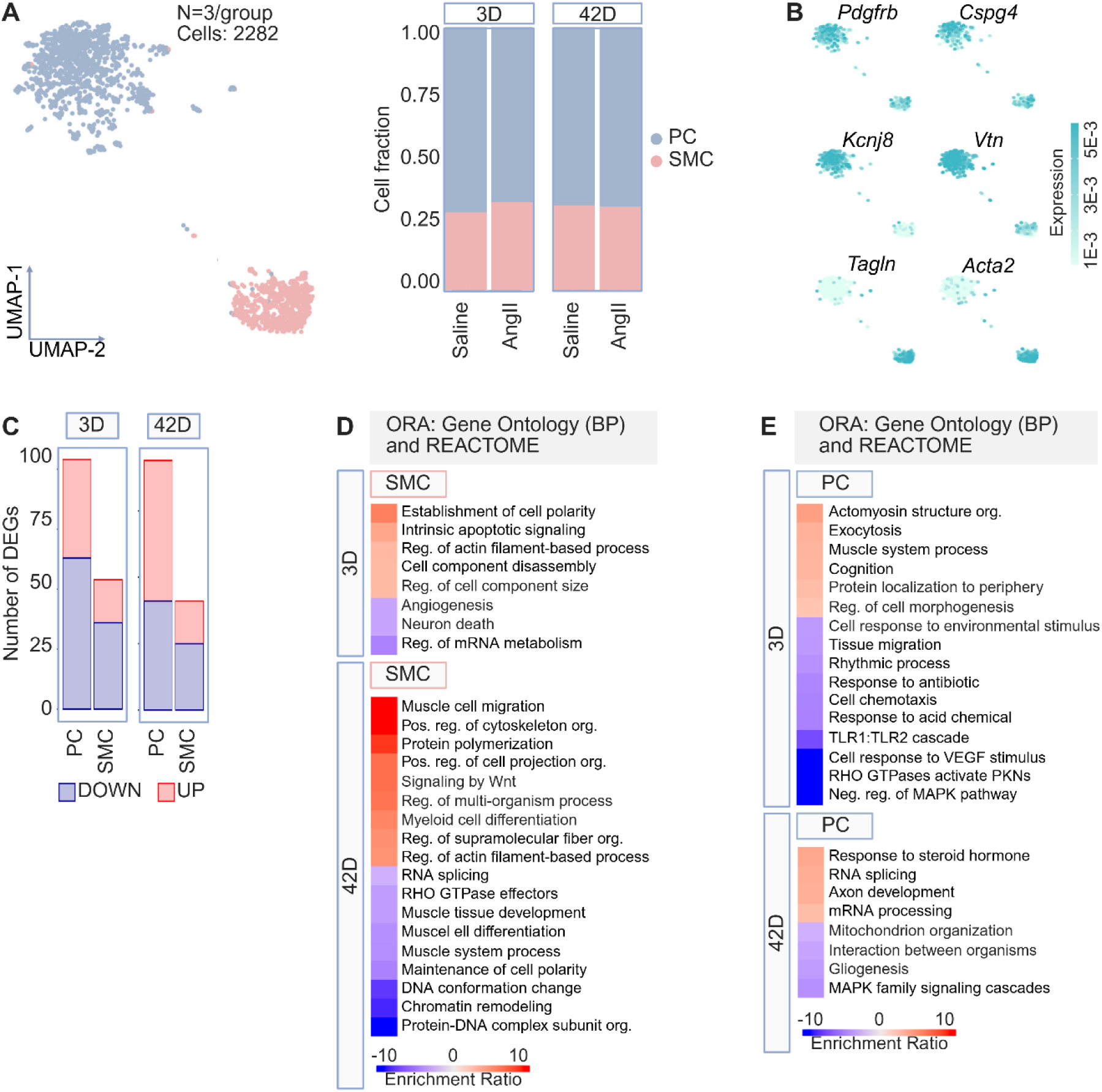
Transcriptional changes in mural cells in early and late AngII HTN. **(A)** Left: UMAP analysis of 2282 mural cells identified smooth muscle cell and pericyte subtypes in merged single cell transcriptomes: pericyte (PC), smooth muscle cell (SMC). Right: Bar graph showing the relative frequencies of smooth muscle cells and pericytes across time points and treatments. **(B)** From left to right, feature plots showing the expression of genes which characterize SMC and PC. **(C)** Bar graph depicting the number of significant upregulated and downregulated differentially expressed genes (DEG) in smooth muscle cells and pericytes in response to 3 days or 42 days of AngII treatment. **(D)** Heatmaps representing the time point-specific enriched biological pathways in AngII-treated SMC, determined through overrepresentation analysis (ORA) of upregulated and downregulated DEG (*p*<0.05, logFC>2). Scale bar represents the enrichment score of genes within a given pathway. **(E)** Heatmaps representing the time point-specific enriched biological pathways in AngII-treated PC, determined through ORA of upregulated and downregulated DEG (*p*<0.05, logFC>2). Scale bar represents the enrichment score of genes within a given pathway.

**Figure S5.**
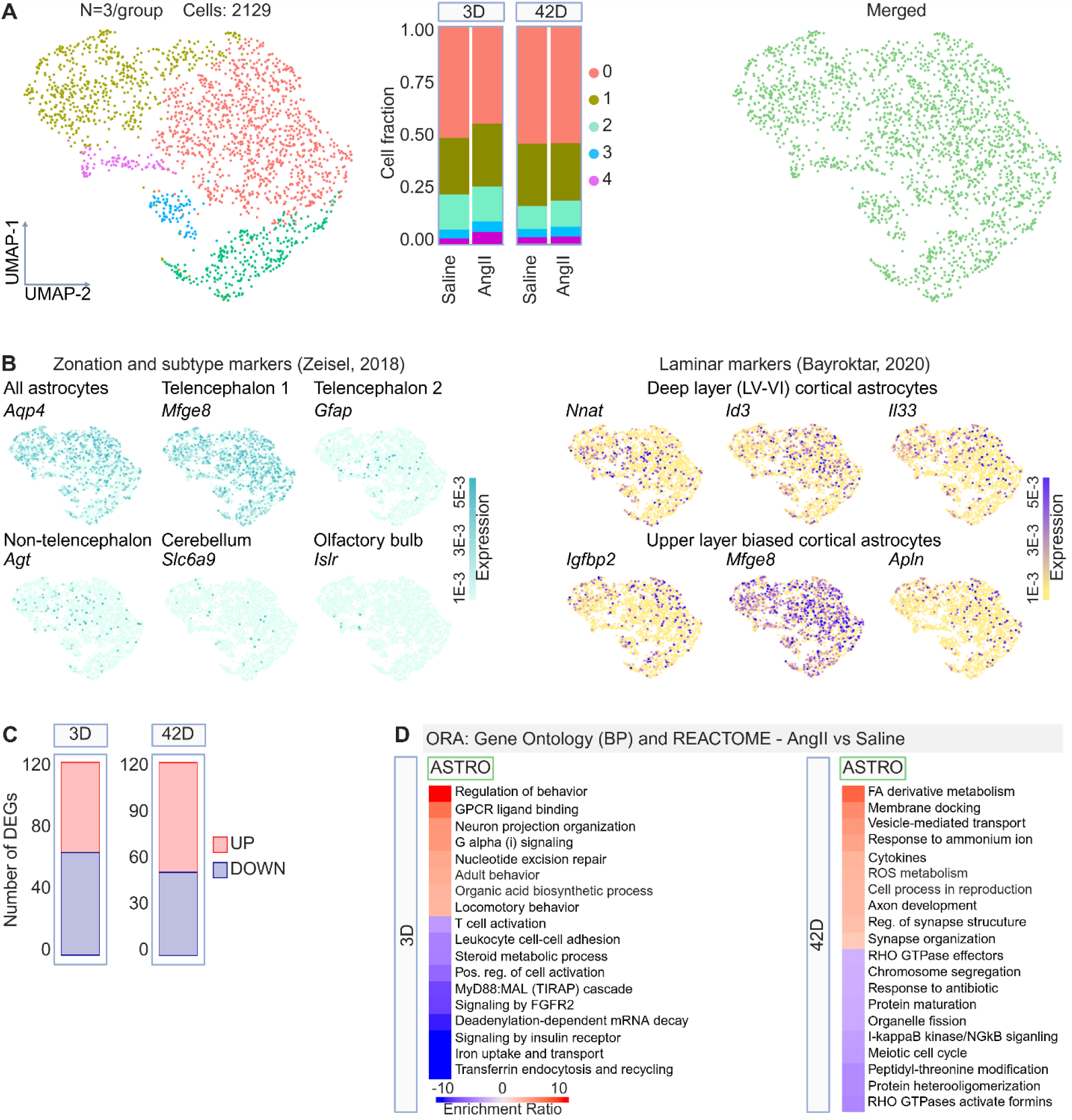
Transcriptional changes in astrocytes in early and late AngII HTN. **(A)** Left: UMAP analysis of 2129 astrocytes. Middle: Bar graph showing the relative frequencies within each sub-cluster across time points and treatments. Right: UMAP of merged astrocyte clusters. **(B)** Left: Feature plots depicting the expression of genes which characterize astrocyte subtypes by gross anatomical location. Right: Feature plots depicting the expression of genes which characterize astrocyte subtypes according to upper versus lower cortical layer zonation. **(C)** Bar graph depicting the number of significant upregulated and downregulated differentially expressed genes (DEG) in response to 3 days or 42 days of AngII. **(D)** Heatmaps representing the time point-specific enriched biological pathways in AngII-treated astrocytes determined through overrepresentation analysis (ORA) of upregulated and downregulated DEG (p<0.05, logFC>2). Pathways are derived from the Gene Ontology Biological Process and Reactome libraries. Scale bar represents the enrichment score of genes within a given pathway.

**Figure S6.**
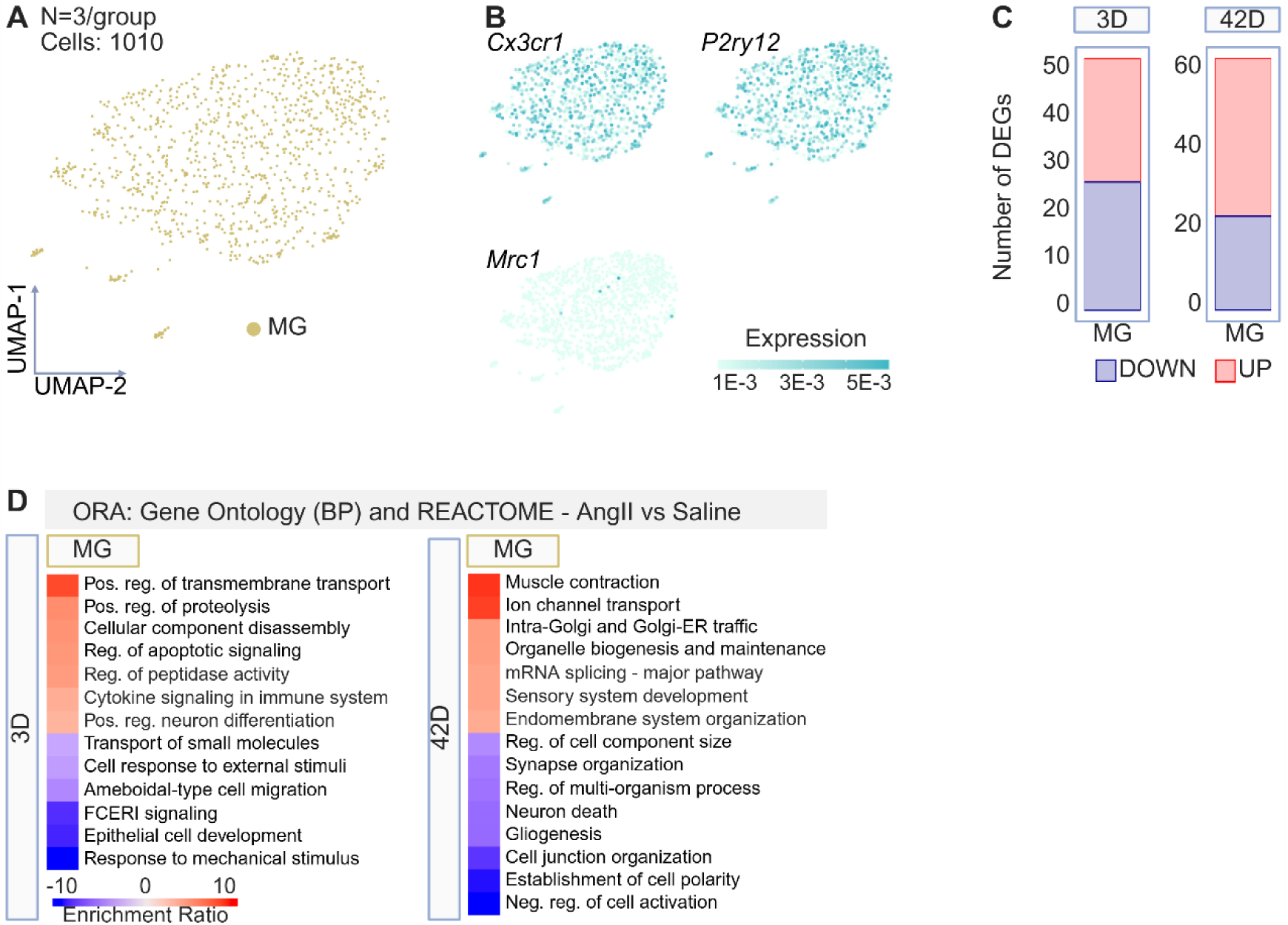
Transcriptional changes in microglia in early and late AngII HTN. **(A)** Left: UMAP analysis of 1010 microglia. **(B)** From top left to bottom right, feature plots depicting the expression of genes which characterize microglia and macrophages. The relative lack of Mrc1 expression (bottom right) indicates that macrophages are absent from the microglial population. **(C)** Bar graph showing the number of significant upregulated and downregulated differentially expressed genes (DEG) in microglia in response to 3 days or 42 days of AngII treatment. **(D)** Heatmaps of the time point-specific enriched biological pathways in AngII-treated microglia determined through overrepresentation analysis (ORA) of upregulated and downregulated DEG (p<0.05, logFC>2). Scale bar represents the enrichment score of genes within a given pathway.

**Figure S7.**
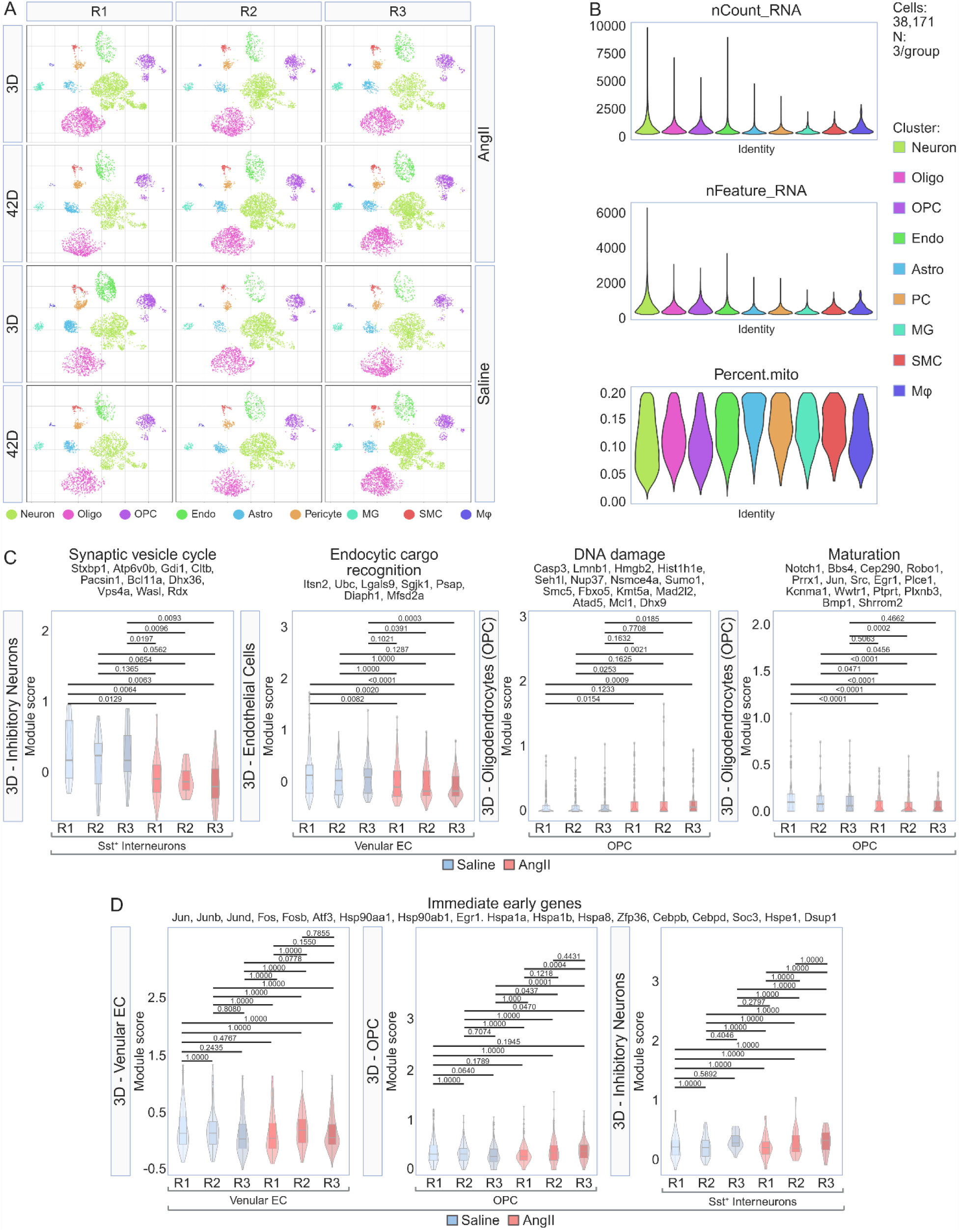
Quality control metrics for the transcriptomic data, related to STAR methods. **(A)** UMAP analysis of the identified clusters, split by replicate (replicate 1, R1; replicate 2, R2; replicate 3, R3), time point (3 days, 42 days), and treatment (saline, AngII). Notice the consistency of cell-type composition of cell cluster across biological replicates. **(B)** Violin plots showing the distribution of total UMI counts per cell (nCount_RNA), genes detected per cell (nFeature_RNA), and percentage of mitochondrial genes identified after thresholding. **(C)** Violin and box plots showing module scores, split by replicate, of the gene sets used to guide subsequent experimental validations. These analyses highlight the consistency of the treatment effect among biological replicates. *p*<0.05. Boxplots represent the median ± IQR. Statistical analysis performed using the Kruskal-Wallis test with Dunn’s correction for multiple comparisons. **(D)** Violin and box plots showing module scores of immediate early genes (IEF), split by replicate, in relevant cell types. These analyses indicate the absence of IEG transcriptomic changes, reflecting cell stress, induced by cell isolation and processing procedures. *p*<0.05. Boxplots represent the median ± IQR. Statistical analysis performed using the Kruskal-Wallis test with Dunn’s correction for multiple comparisons

## STAR METHODS

### Experimental model

#### Animal information and housing

All procedures were approved by the Institutional Animal Care and Use Committee of Weill Cornell Medicine and performed in accordance with the National Institutes of Health Guide for the Care and Use of Laboratory Animals. All studies were performed in a blinded fashion in male mice aged 3-5 months. Mice were housed socially in ventilated cages with ad libitum access to food and water (22 °C; 45-65% humidity; 12:12 hour light:dark cycle with the light phase from 07:00 to 19:00). Strains used include: C57BL/6 mice (WT, weight 25-30 g; JAX, Jackson Laboratory) and nNOS Cre driver mice (C57BL/6-*Nos1*^Cre^, JAX strain no. 017526). To enable the identification of active nNOS neurons, GCaMP7f was expressed in *Nos1*^Cre^ mice using i.v. injection of an adeno-associated virus containing a Cre responsive flip-excision switch (AAV PHP.eB Syn-FLEX-GCaMP7f)^91^. *Nos1*^cre^ mice were congenic on a C57BL/6 genetic background.

#### Experimental design

Mice were treated with slow-pressor AngII for 3 days or 42 days to contrast those transcriptional alterations which emerge prior to HTN and those which develop at the time of cognitive impairment^14,32^. To delineate the direct contributions of AngII relative to the elevated BP, mice were treated with one of saline, a non-pressor dose of AngII, or phenylephrine^33^ for 6 days. Lastly, to investigate whether the early effects of slow-pressor AngII could be reversed, mice were treated with saline or slow-pressor AngII, with or without losartan (600 mg/L, I934, AK Scientific)^58,59^, an AT1R blocker, for 8 days. Losartan was added to the drinking water on day 5 AngII treatment, once HTN had fully developed.

### Method details

#### Minipump implantation and drug delivery

Mice were randomized to treatment group and anesthetized by isoflurane inhalation (5% induction, 2% maintenance) for subcutaneous (s.c.) implantation of an osmotic minipump (ALZET model 2006, DURECT Corporation). Osmotic minipumps were loaded with saline, a slow-pressor dose of AngII (600 ng/kg/min, Sigma-Aldrich, A9525), a non-pressor dose of AngII (200 ng/kg/min, Sigma-Aldrich, A9525), or a pressor dose of phenylephrine (3 ug/kg/min, Sigma-Aldrich, P6126), as previously described^14,32^, then incubated at 37 °C for 72 hours prior to implantation, as per the manufacturer’s instructions. Systolic BP (SBP) was monitored in awake mice using tail-cuff plethysmography (Hatteras, Multi-channel BP analysis software, MC4000)^108^. Mice were acclimated to tail-cuff plethysmography for 1 week prior to surgery. The infusion of AngII at 200 ng/kg/min does not affect BP (Fig. 3D), whereas 600 ng/kg/min elicits a time-dependent rise in BP (Fig. 1A) consistent with the “slow pressor” model of AngII-induced HTN^22^. The increase in BP induced by phenylephrine follows a similar temporal progression^33^ (Fig. 3D).

#### Novel object recognition test

The novel object recognition test (NOR) task was conducted in a plastic box measuring 29 cm × 47 cm × 30 cm high, as previously described^13,138^. Stimuli consisted of plastic objects of a similar size that varied in color and shape. A video camera was used to record the testing session for offline analysis using AnyMaze software. Mice were acclimated to the testing room for 1 hour prior to habituation/testing. On day 1, mice were acclimated to the testing chamber for 5 minutes (habituation). On day 2, mice were placed into the same chamber in the presence of 2 identical sample objects and were allowed to explore for 5 minutes. After an intersession interval of 1 hour, mice were placed in the same box, but 1 of the 2 objects was replaced by a novel object. Mice were allowed to explore for 5 minutes. Between trials, the testing chamber was cleaned with 10% ethanol in water to minimize olfactory cues. Exploratory behavior was manually assessed by an experimenter blinded to the treatment group. Exploration of an object was defined as the mouse sniffing the object or touching the object while looking at it. A minimal exploration time for both objects (total exploration time) during the test phase (5 seconds) was used. The amount of time taken to explore the novel object was expressed as percentage of the total exploration time and provides an index of recognition memory.

#### Mouse brain single-cell suspensions

A total of 12 cell suspensions were generated by adapting protocols from^43^, optimized for cortical tissue, with 3 biologic replicates per treatment group. Within each replicate, 2 cortical hemispheres were pooled from 2 identically treated mice. Mice were anesthetized by isoflurane inhalation (5% induction, 2% maintenance) and perfused transcardially with ice-cold sucrose-HEPES “Cutting Buffer” containing (in mM) 110 NaCl, 2.5 KCl, 10 HEPES, 7.5 MgCl_2_, 25 glucose, and 75 sucrose (∼350 mOsm kg−1). The brain was removed, placed in ice-cold Cutting Buffer, then transferred to a slicing chamber filled with ice-cold Cutting Buffer. 400 μm thick brain slabs were cut with a vibrating microtome. Slabs containing the regions of interest were gently transferred to a dissection dish with ice-cold dissociation buffer (DB) containing (in mM): 82 Na_2_SO_4_, 30 K_2_SO_4_, 10 HEPES, 10 glucose and 5 MgCl_2_. Regions of interest were isolated, then transferred into a 15 ml falcon tube filled with 5 ml of DB + enzyme buffer (DEB) consisting of DB with 3 mg/ml of Protease XXIII (Sigma-Aldrich, P5380), 10 units/ml of Papain (Sigma-Aldrich, P4762), 0.5 mM L-Cysteine (Sigma-Aldrich, 168149), and 0.25 mM EDTA (Worthington, LK003153). Digestion was performed at 34 °C for 2 hours. Tubes containing digested tissue were transferred onto ice and the DEB was replaced with 10 ml of a DB + stop solution (DBS) containing DB and 1 mg/ml Trypsin Inhibitor (Sigma-Aldrich, T6522), 2 mg/ml BSA (Sigma-Aldrich, A2153), and 1 mg/ml Ovomucoid Protease Inhibitor (Worthington, LK003153). Digested tissues were then dissociated using a series of 4 fire-polished Pasteur pipets with successively smaller bores. Bubbles were avoided. Falcon tubes containing 10 ml of dissociated cells were then centrifuged at 300 g for 10 minutes. The supernatant was removed and discarded, taking care not to disturb the cell pellet. The pellet was resuspended in 5 ml of DBS and centrifuged again under the same conditions. The supernatant was once again removed, and the cleaned cell pellet was resuspended to a volume of 3 ml in DB containing 0.01% BSA (w/v). Suspensions were passed into a new tube through a pre-wet 40 μm filter and held on ice. 2 samples of 10 μl were drawn from the suspension and mixed 1:1 with 10 μl of 2× dye mix containing DB and 20 μM EthD-1 (Thermo Fisher Scientific, L-3224), 20 μM Calcein-AM (Thermo Fisher Scientific, L-3224), and 40 μM Hoechst 33342 (Thermo Fisher Scientific, 62249). After 5 minutes of incubation at room temperature, 10 μl from each sample were loaded onto a haemocytometer and imaged using a fluorescent Olympus IX83 microscope. For each of the 2 samples, three random locations were imaged using DIC and three fluorescent channels to capture the dyes. These images were used to calculate cell concentrations and metrics of cell intactness. Afterward, cells were resuspended in 0.01% BSA-PBS at a concentration of 100,000 cells/ml for Drop-seq processing.

#### Generation of single cell RNA libraries by Drop-seq

Single-cell transcriptomes of cortical brain cells were prepared by Drop-seq as described^42,44^, with modifications. Cells were resuspended in 0.01% BSA-PBS to a final concentration of 100 cells/μl. Barcoded capture beads (ChemGenes Corporation, Wilmington, MA) were resuspended in 1.8 ml lysis buffer consisting of 4M Guanidine HCL (Thermofisher Scientific, 24110), 6% Ficoll PM-400 (Sigma-Aldrich, F4375), 0.2% Sarkosyl (Sigma-Aldrich, 61747), 20mM EDTA (Thermofisher Scientific, 17892), 200mM Tris pH 7.5 (Sigma-Aldrich, DB0339), and 50mM DTT (Sigma-Aldrich, D0632) at a concentration of 120 beads/μl. A 5 mm diameter, 1.7 mm thick PVDF encapsulated magnetic stir disc and rotary magnetic tumble stirrer (V&P Scientific, VP772DP-N42-5-2) was used along with a 3 ml syringe that contained the beads in lysis buffer to maintain the beads in suspension. Next, single cells and beads were encapsulated in nanoliter-scale droplets using a Drop-seq microfluidic device coated with Aquapel (FlowJEM); flow rates were 15 ml/hour for the Droplet generation oil (BioRad, 1863005) and 4 ml/hour for the cells and beads. Each run lasted approximately 18 minutes. After removing the oil, droplets were resuspended in 30 ml of room temperature 6X SSC (Promega, V4261) and 1 ml perfluorooctanol (Sigma-Aldrich, 370533) and shaken vigorously 6 times vertically to break the droplets. Beads were then captured by loading them into a 20 ml syringe with an attached 0.22 μm Millex-Gv syringe filter (Millipore Sigma, SLGV033R), as previously described^139^. Beads were washed with 2 x 20 ml of ice cold 6X SCC. After washing, the syringe filter was inverted and flushed 3x with 10 ml of ice cold 6x SSC using a 10 ml syringe. Beads were collected from the resultant solution via centrifugation at 1,250 x g for 2 min at 4°C with low brake setting. The remainder of the Drop-seq protocol was performed as previously described^43^. cDNA was amplified by PCR using the following parameters: 95 °C (3 min); 4 cycles of 98 °C (20 s), 65 °C (45 s), and 72 °C (3 min); 11 cycles of 98 °C (20 s), 67 °C (20 s), and 72 °C (3 min). Libraries were quantified by quantitative PCR and checked for quality and size distribution on a Bioanalyzer (Agilent) before sequencing on an Illumina NextSeq500 instrument using the 75 cycle High Output v2 kit (Genomics Core Facility, Cornell University, Ithaca, NY). Three to four libraries were multiplexed into a single run. We loaded a 1.8 pM library and provided Drop-seq Custom Read1 Primer at 0.3 μM in position 7 of the reagent cartridge without PhiX spike-in using a read configuration of 20 bases (Read1), 8 bases (Index1), and 64 bases (Read2).

#### Data pre-processing

Demultiplexed fastq files were cleaned of reads not passing the Illumina Passing Filter with fastq_illumina_filter (version 0.1) and processed with the Drop-seq Tools (version 2.3.0) pipeline^42^. Briefly, each transcriptome Read2 was tagged with the cell barcode (bases 1 to 12) and unique molecular identifier (UMI) barcode (bases 13 to 20) obtained from Read1, trimmed for sequencing adapters and poly-A sequences, and aligned using STAR v2.7.3a^140^ to the mouse reference genome assembly (Ensembl GRCm38.94 release). Reads aligning to exons were tagged with the respective gene symbol and counts of UMI-deduplicated reads per gene within each singular cell barcode were used to build a digital gene expression (DGE) matrix. The DGE matrix contained 40,000 cell barcodes associated with the highest numbers of UMIs. Cells with fewer than 200 UMIs, more than 10,000 UMIs, or more than 20% mitochondrial genes were excluded (Fig. S7B). Doubletfinder (RRID:SCR_018771)^141^ was used to computationally detect cell doublets with an expected doublet rate of 5% as the input parameter. Cells tagged with a “Doublet” call were removed. The corrected DGE matrices were merged into a single matrix.

#### Bioinformatic analysis and statistics of RNAseq data

All bioinformatic analysis was performed using R (RRID:SCR_001905). Seurat (version 4.1.0; RRID:SCR_016341)^142^ was used for down-stream analysis: Counts were log-normalized for each cell using the natural logarithm of 1 + counts per ten thousand. The 3000 most variable genes were identified by calling FindVariableFeatures. We next standardized expression values for each gene across all cells by Z-score transformation (ScaleData). Principle Component Analysis (PCA) was performed on the scaled variable gene matrix. We used Uniform Manifold Approximation and Projection (UMAP)^143^ for dimensional reduction and visualization of merged libraries in a two-dimensional space with preset parameters by invoking the RunUMAP function in Seurat, utilizing the first 40 components of the PCA. We used the Louvain algorithm as implemented in FindClusters with a resolution setting of 0.7 to perform graph-based clustering on the neighbor graph that was constructed with the FindNeighbors function call. After clustering, we used the model-based analysis of single cell transcriptomics (MAST) algorithm^144^ in the FindAllMarkers function to find differentially expressed genes in each cluster based on the log-normalized expression matrix with the following parameters: only.pos = T, min.pct = 0.1, logfc.threshold = log2(1.5), max.cells.per.ident = 2000. We performed unsupervised cell type annotation using the Seurat Integration and FindIntegrationAnchors and TransferData functions with Dropviz^42^, Tabula Muris^92,145^, EndoDB^146^, and LinnersonLab^147^. Assignments were further manually validated by scoring the 10 most differentially expressed genes for the presence of canonical marker genes for each cell type. On these bases, we assigned the metacells to neurons, mature oligodendrocytes (OL), oligodendrocytes precursor cells (OPC), endothelial cells (EC), astrocytes, microglia (MG), mural cells, and macrophages (Mφ). UMAP analyses at the sample level confirmed uniform cell clustering between biological replicates, highlighting the lack of batch effects (Fig. S7A). In agreement with scRNAseq studies using a similar digestion protocol and microfluidics platform^42^, the number of astrocytes is less than that observed in studies using in situ histological assessments (∼20%)^148^. However, the frequency reported herein aligns with that described in the Saunders database (5.7% versus 6.7%)^42^.

To achieve further resolution of cell states, individual count matrices were generated based on the initial cluster designation and steps 1-8 were repeated with the following modifications: 2000 variable features were selected, and resolution settings of 0.5, 0.2, 0.3, 0.3, 0.3, and 0.4 were used for the FindClusters function for endothelial cells, oligodendrocytes, neurons, microglia, astrocytes, and mural cells, respectively. Differentially expressed genes with an FDR < 0.05 were ranked by their log2 fold change and z-scores were computed on the average gene expression across clusters for visualization in heatmaps. On these bases, we assigned the metacells to endothelial cells (arterial (aEC), arterial-capillary (aCapEC), capillary-venous (vCapEC), and venous (vEC)), oligodendrocyte-lineage cells (mature oligodendrocyte (Mol), newly formed oligodendrocyte (Nfol), committed oligodendrocyte precursor (Cop), cycling oligo-precursor (Cyp), and oligodendrocyte precursor (Opc)), microglia (MG), macrophages (Mφ), vascular smooth muscle cells (SMC), pericytes (PC), and neurons (excitatory and inhibitory).

#### Differential gene expression analysis

Following pseudobulk conversion of individual experiments using the aggregateAcrossCells function of the scuttle package^149^, differential gene expression for general cell clusters was computed with limma-voom using default parameters^150^. To detect the DEGs for each cell sub-type between the saline and AngII treatment groups (P < 0.05, |fold change| > 2), a pseudobulk-based method, edgeR (V4.2.2; RRID:SCR_012802), was employed^151^. Although the sample sizes employed in this study (n=3) may limit the statistical power in the sample-based (i.e. pseudobulk) analysis, this approach performs better than cell-based statistical methods that are prone to false positives by inflating the p-value^152^. Furthermore, the distribution of cells within clusters was uniform across biological replicates (Fig.S7A) and the AngII treatment effect was consistent across key gene sets (Fig. S7C), attesting to the reliability of the data.

#### Gene pathway analysis

Analysis of enriched pathways in cell clusters of the AngII treatment group was performed using WebGestaltR 2019 (RRID:SCR_024312)^153^. EdgeR DEGs were filtered (P < 0.05, fold change > 2) then evaluated for enrichment in the following functional categories: Gene Ontology Biological Process (Daily build) and Reactome (Version 66). The affinityPropagation function of the WebGestaltR package was called to reduce redundancy in the reported pathways. Computation of immediate-early gene (IEG) module scores at the sample level confirmed that that the results of the pathway analyses were not driven by cell dissociation-induced stress and subsequent IEG upregulation (Fig. S7D).

#### Module score calculation and visualization

The AddModuleScore function in the Seurat package was used to calculate the cell-type, timepoint, and treatment specific enrichment of target gene sets. Each score was calculated using gene sets derived from the GO Browser, as previously reported^154,155^. All genes used for score calculation can be found in the supplementary data (Table S2). Module scores were visualized in R using the ggplot2 (V3.3.6) package (RRID:SCR_014601). Data are expressed as mean ± SEM (violin plots) and mean with interquartile range (inlaid box plots).

#### Brain histology

Mice were anesthetized with sodium pentobarbital (100 mg/kg, i.p.) and perfused transcardially with PBS followed by 4% paraformaldehyde (PFA; Millipore-Sigma, 158237) in PBS. Brains were dissected, post-fixed in 4% PFA overnight, dehydrated in 30% PBS-sucrose solution for 1-2 days, then frozen using dry ice. Frozen brains were embedded in Epredia™ M-1 Embedding Matrix (Thermo Ficher Scientific, 1310), cut in coronal sections using a Leica CM3050S cryostat (Leica Biosystems), then mounted on slides for either immunoflorescence (IF) or RNA-fluorescence in situ hybridization (FISH).

#### Immunofluorescence

18 µm-thick coronal sections were permeabilized with 0.5% Triton X-100 (Sigma, X100) in PBS (PBST), blocked with 5% normal donkey serum (NDS; Millipore-Sigma, 566460) in 0.1% PSBT for 1 hour, then incubated overnight at 4°C with primary antibodies in 1% NDS-0.1% PBST. Senescent cells were detected using anti-P27 (1:400, rabbit, 25614-1-AP, Proteintech), anti-P21 (1:200, rabbit, ab188224, Abcam), and anti-P16 (1:100, rabbit, ab211542, Abcam). Blood vessels were stained with anti-PECAM-1, clone 2H8 (1:100, goat, MAB1398Z, Millipore-Sigma). OPC were identified using anti-Olig2 (1:200, rabbit, ab109186, Abcam) and anti-Pdgfrα (1:200, goat, AF1062, Biotechne), and double-stranded DNA breaks were detected with anti-phospho(Ser^139^)- histone H2A.X (1:100, mouse, MA1-2022, ThermoFisher Scientific). Collagen IV was stained using anti-collagen IV (1:400, rabbit, ab6586, Abcam). After overnight incubation, sections were washed 3 x 5 min with 0.1% PBST, then incubated with secondary antibodies in 1% NDS-0.1% PBST for 1 hour at room temperature. Secondary antibodies used are as follows: AF-488 donkey anti-rabbit (1:200, A-21206, ThermoFisher Scientific), AF-647 donkey anti-goat (1:200, A-21447, ThermoFisher Scientific), AF-647 goat anti-Armenian hamster (1:200, A78967, ThermoFisher Scientific), AF-488 donkey anti-mouse (1:200, A-21202, ThermoFisher Scientific) and Cy3 donkey anti-rabbit (1:200, AP182C, Sigma-Aldrich). Next, sections were washed 2 x 5 min with 0.1% PBST, followed by a 1 x 5 min incubation in 0.1% PBST-DAPI (1:100, 12.5 ng/ul). Sections were mounted with FluorSave Reagent (Millipore-Sigma, 345789) and visualized using a Leica TCS SP8 confocal microscope (Leica Biosystems). Images were analyzed using Fiji (RRID:SCR_002285), ImageJ (RRID:SCR_003070), or Imaris Software (v10.0) (RRID:SCR_007370). Senescence marker (P27, P21, and P16) colocalization with CD31^+^ endothelial cells was imaged at 20x magnification. Using ImageJ, masks of P27^+^, P21^+^, and P16^+^ areas were overlayed onto a mask of the CD31^+^ positive area in a 240 um^2^ section, then the area of overlap between the two masks was divided by the total CD31^+^ zone. To quantify double-strand DNA breaks in OPC, OPC were first identified by colocalization of DAPI, Olig2, and PDGFRA at 63x magnification (oil immersion). Then, using ImageJ, mean gray values of the γ-H2A.X channel were calculated within the DAPI+Olig2 positive area in confirmed OPC across 3 sequential 1 um thick Z-slices, with the background subtracted from the mean of each. A total of 10 cells were quantified for each animal. For collagen IV, images were captured at 20x magnification, then, using ImageJ, the collagen IV^+^ areas surrounding each CD31^+^ vessel were traced, the background subtracted, and the mean gray value measured. Collagen IV intensity was quantified in 8-10 vessels per 240 um^2^ section, with 3 sections assessed per animal.

#### FISH

RNA-FISH was performed using RNAscope Multiplex Fluorescent Kit v2 (ACD-Bio-Techne, 323110) as per manufacturer’s instructions. Briefly, 10-µm thick sections were processed using RNAscope hydrogen peroxide, followed by boiling in Target Retrieval solution, dehydration with 100% ethanol, and incubation in Protease III solution. Afterwards, tissue sections were hybridized with the target probes for 2 hours at 40 °C, followed by a series of signal amplification and washing steps. Hybridization signals were detected by fluorescent signal using peroxidase-based Tyramide Signal Amplification-Plus Fluorescein or TSA Plus Cyanine 5 (Thermo Fisher Scientific, 16423834). Finally, sections were counterstained with DAPI, and cover slips were applied using ProLong Gold Antifade Mounting agent (Thermo Fisher Scientific, P36934). Images were acquired using a Leica TCS SP8 confocal microscope and analyzed using Fiji (RRID:SCR_002285) or Imaris Software (v10.0). When FISH and IF staining were combined, the slides were washed in RNAscope wash buffer after the development of Tyramide signal, and IF was then sequentially performed as described above sans permeabilization.

#### Cellular senescence detection

SA-β-Gal staining was achieved with the Cellular Senescence Detection Kit (Cell Bio Labs, CBA-230). SA-β-Gal catalyzes the hydrolysis of X-gal, which produces a blue color in senescent cells. For tissue staining in the mouse cortex, brains were fixed in 4% PFA overnight, dehydrated in 30% PBS-sucrose solution for 1-2 days, frozen, then sectioned into 10 μm slices using a cryostat. After washing 3 times with PBS, slices were incubated in the 1x X-Gal solution overnight at 37°C in a humidifying container. The next day, sections were washed in PBS 1x and images were captured and analyzed using standard light microscopy. Quantification was performed by counting vessel-associated cells positive for SA-β-Gal in a 500 um^2^ area, with 5 total areas counted and averaged per animal.

#### Nitric oxide detection by DAF-FM

As previously described^156–159^, the brains of mice treated for 3 days with saline (sham treatment) or slow-pressor AngII, 6 days with non-pressor AngII or phenylephrine, or 8 days with slow-pressor AngII with or without losartan, were removed and immersed in artificial cerebral spinal fluid (aCSF). Coronal cortical slices (350 µm) were cut using a vibratome and then incubated for 1 hour each at 36 °C in oxygenated l-aCSF containing 0.02% Pronase (Millipore-Sigma, 10165921001) and 0.02% thermolysin (Millipore-Sigma, P1512). Next, the slices were mechanically dissociated and incubated with DAF-FM (4-amino-5-methylamino-2′,7′-difluorofluorescein) diacetate (5 µM; Thermo Fisher Scientific, D23844) in oxygenated l-aCSF for 30 minutes, then rinsed in control buffer (no DAF-FM) for 30 minutes. Target neurons were identified based on their size and presence of processes. DAF-FM is sequestered into cells and converted to DAF-2 by intracellular esterases, rendering it membrane impermeable. NO and NO-derived species *N*-nitrosylate DAF-2, leading to the formation of DAF-2T, a green fluorescent triazole^160^. Time-resolved fluorescence (FITC filter) was measured at 30 second intervals with a Nikon diaphot 300 inverted microscope equipped with CCD digital camera (Princeton Instruments) and using IPLab software (Scanalytics). The DAF-FM fluorescence intensity is expressed as Ft/F0, where F0 is the baseline fluorescence before application of NMDA or vehicle, and Ft is the fluorescence in the same cell after the application of NMDA.

#### Trans-callosal conduction velocity recordings

Trans-callosal recordings of action potential latency were performed as described^80^, with the following modifications. Mice treated with saline or AngII for 3D or 42D were anesthetized with isoflurane and decapitated. Brains were removed and immersed in ice-cold sucrose + artificial cerebrospinal fluid (s-ACSF) gassed with 95% O_2_ and 5% CO_2_. Next, brains were fixed (oriented caudal end down) to the specimen disc of a Leica VT-1000s vibratome cutting chamber using Loctite 404 glue (Henkel Adhesives, 135465). The cutting chamber was packed with ice. Brains were sectioned at 350 µm. Slices corresponding to Plates 35-45 in the Paxinos and Franklin atlas were transferred to an incubation chamber containing room temperature (RT) oxygenated lactic acid + acSF (l-ACSF). Slices were allowed to recover for one hour prior to recording. After transfer to the submersion-type recording chamber, slices were continuously perfused with oxygenated l-ACSF. A Meiji EMZ-250 stereomicroscope (Meiji Techno) was used to position the stimulating and recording electrodes: the stimulating electrode was placed on the outer edge of the corpus callosum (CC) 0.5 mm lateral to midline, while the tip of the recording electrode was placed within the CC of the opposite hemisphere 0.5 mm, 1.0 mm, or 2.0 mm away from the stimulating electrode. Field potential recording of evoked compound action potentials (CAP) were attained using the following protocol: Episodic stimulation was provided over 25 sweeps (1 sweep every 5 seconds), with sampling performed every 5 µs (200 kHz) over a 100-ms period each sweep (lowpass Bessel filter was set to 10 kHz). Recorded data were stored for offline analysis. CC conduction velocity changed linearly with the inter-electrode distance. The most positive component of the peak latency was measured and plotted as a function of inter-electrode distance. For all experiments, an Axopatch 200a amplifier (Molecular Devicesand 395 linear stimulus isolator (WPI) connected to an Axon 1332A Digidata and compute with the pClamp10 software (Molecular Devices) were used for data acquisition. Recording glass micropipettes were pulled from WPI borosilicate capillaries using a two-step protocol (P-80, Sutter Instrument), back-filled with l-ACSF, and connected to the silver/silver-chloride wire of the electrode holder. A concentric bipolar microelectrode (FHC) was used as the stimulating electrode.

### Quantification and statistical analysis

After testing for normality (Shapiro-Wilk test), intergroup differences were analyzed using the unpaired two-tailed t-test, paired two-tailed t-test, repeated measures two-way ANOVA with Šidák’s correction, one-way ANOVA with Tukey’s test, or one-way ANOVA with Dunnet’s test, as appropriate. If non-parametric testing was indicated, intergroup differences were assessed using the Wilcoxon signed-rank test (non-normal distribution) or Mann-Whitney U-test. The specific statistical method employed for a given comparison is indicated in the figure legend, as appropriate. Statistical tests throughout the manuscript were performed using either Prism 9 (GraphPad) or the rstatix (V0.7.2) package (RRID:SCR_021240). Data are expressed as mean ± SEM and differences are considered statistically significant when p<0.05. No data were excluded in the preparation of this manuscript.

### Additional information

A portal has been established to allow user friendly access to our processed single-cell RNA sequencing data: https://anratherlab.shinyapps.io/angii_brain/.

## REFERENCES

1. Forouzanfar, M.H., Liu, P., Roth, G.A., Ng, M., Biryukov, S., Marczak, L., Alexander, L., Estep, K., Hassen Abate, K., Akinyemiju, T.F., et al. (2017). Global Burden of Hypertension and Systolic Blood Pressure of at Least 110 to 115 mm Hg, 1990-2015. Jama 317, 165–182. 10.1001/jama.2016.19043.

2. Pacholko, A., and Iadecola, C. (2024). Hypertension, Neurodegeneration, and Cognitive Decline. Hypertension. 10.1161/HYPERTENSIONAHA.123.21356.

3. Abdulrahman, H., van Dalen, J.W., den Brok, M., Latimer, C.S., Larson, E.B., and Richard, E. (2022). Hypertension and Alzheimer’s disease pathology at autopsy: A systematic review. Alzheimers Dement 18, 2308–2326. 10.1002/alz.12707.

4. Lackland, D.T., Roccella, E.J., Deutsch, A.F., Fornage, M., George, M.G., Howard, G., Kissela, B.M., Kittner, S.J., Lichtman, J.H., Lisabeth, L.D., et al. (2014). Factors influencing the decline in stroke mortality: a statement from the American Heart Association/American Stroke Association. Stroke 45, 315–353. 10.1161/01.str.0000437068.30550.cf.

5. Lee, C.J., Hwang, J., Kang, C.Y., Kim, H.C., Ryu, D.R., Ihm, S.H., Kim, Y.J., Shin, J.H., Pyun, W.B., Kim, C., and Park, S. (2021). Protective effect of controlled blood pressure on risk of dementia in low-risk, grade 1 hypertension. J Hypertens 39, 1662–1669. 10.1097/hjh.0000000000002820.

6. Adesuyan, M., Jani, Y.H., Alsugeir, D., Howard, R., Wong, I.C.K., Wei, L., and Brauer, R. (2023). Trends in the incidence of dementia in people with hypertension in the UK 2000 to 2021. Alzheimers Dement (Amst) 15, e12466. 10.1002/dad2.12466.

7. Iadecola, C. (2017). The Neurovascular Unit Coming of Age: A Journey through Neurovascular Coupling in Health and Disease. Neuron 96, 17–42. 10.1016/j.neuron.2017.07.030.

8. Schaeffer, S., and Iadecola, C. (2021). Revisiting the neurovascular unit. Nat Neurosci 24, 1198–1209. 10.1038/s41593-021-00904-7.

9. Jennings, J.R., Muldoon, M.F., Ryan, C., Price, J.C., Greer, P., Sutton-Tyrrell, K., van der Veen, F.M., and Meltzer, C.C. (2005). Reduced cerebral blood flow response and compensation among patients with untreated hypertension. Neurology 64, 1358–1365. 10.1212/01.Wnl.0000158283.28251.3c.

10. Delles, C., Michelson, G., Harazny, J., Oehmer, S., Hilgers, K.F., and Schmieder, R.E. (2004). Impaired endothelial function of the retinal vasculature in hypertensive patients. Stroke 35. 10.1161/01.STR.0000126597.11534.3b.

11. Bagi, Z., Brandner, D.D., Le, P., McNeal, D.W., Gong, X., Dou, H., Fulton, D.J., Beller, A., Ngyuen, T., Larson, E.B., et al. (2018). Vasodilator dysfunction and oligodendrocyte dysmaturation in aging white matter. Annals of Neurology 83, 142–152. 10.1002/ana.25129.

12. Muñoz Maniega, S., Chappell, F.M., Valdés Hernández, M.C., Armitage, P.A., Makin, S.D., Heye, A.K., Thrippleton, M.J., Sakka, E., Shuler, K., Dennis, M.S., and Wardlaw, J.M. (2017). Integrity of normal-appearing white matter: Influence of age, visible lesion burden and hypertension in patients with small-vessel disease. Journal of cerebral blood flow and metabolism : official journal of the International Society of Cerebral Blood Flow and Metabolism 37 10.1177/0271678X16635657.

13. Faraco, G., Sugiyama, Y., Lane, D., Garcia-Bonilla, L., Chang, H., Santisteban, M.M., Racchumi, G., Murphy, M., Van Rooijen, N., Anrather, J., and Iadecola, C. (2016). Perivascular macrophages mediate the neurovascular and cognitive dysfunction associated with hypertension. J Clin Invest 126, 4674–4689. 10.1172/jci86950.

14. Santisteban, M.M., Ahn, S.J., Lane, D., Faraco, G., Garcia-Bonilla, L., Racchumi, G., Poon, C., Schaeffer, S., Segarra, S.G., Körbelin, J., et al. (2020). Endothelium-Macrophage Crosstalk Mediates Blood-Brain Barrier Dysfunction in Hypertension. Hypertension 76, 795–807. 10.1161/hypertensionaha.120.15581.

15. Kazama, K., Wang, G., Frys, K., Anrather, J., and Iadecola, C. (2003). Angiotensin II attenuates functional hyperemia in the mouse somatosensory cortex. Am J Physiol Heart Circ Physiol 285, H1890–1899. 10.1152/ajpheart.00464.2003.

16. Kazama, K., Anrather, J., Zhou, P., Girouard, H., Frys, K., Milner, T.A., and Iadecola, C. (2004). Angiotensin II impairs neurovascular coupling in neocortex through NADPH oxidase-derived radicals. Circulation research 95. 10.1161/01.RES.0000148637.85595.c5.

17. Iadecola, C., and Gottesman, R.F. (2019). Neurovascular and Cognitive Dysfunction in Hypertension. Circ Res 124, 1025–1044. 10.1161/CIRCRESAHA.118.313260.

18. Carnevale, L., D’Angelosante, V., Landolfi, A., Grillea, G., Selvetella, G., Storto, M., Lembo, G., and Carnevale, D. (2018). Brain MRI fiber-tracking reveals white matter alterations in hypertensive patients without damage at conventional neuroimaging. Cardiovascular research 114. 10.1093/cvr/cvy104.

19. Carnevale, L., Maffei, A., Landolfi, A., Grillea, G., Carnevale, D., and Lembo, G. (2020). Brain Functional Magnetic Resonance Imaging Highlights Altered Connections and Functional Networks in Patients With Hypertension. Hypertension (Dallas, Tex. : 1979) 76. 10.1161/HYPERTENSIONAHA.120.15296.

20. Siedlinski, M., Carnevale, L., Xu, X., Carnevale, D., Evangelou, E., Caulfield, M.J., Maffia, P., Wardlaw, J., Samani, N.J., Tomaszewski, M., et al. (2023). Genetic analyses identify brain structures related to cognitive impairment associated with elevated blood pressure. European heart journal 44. 10.1093/eurheartj/ehad101.

21. Romero, J.C., and Reckelhoff, J.F. (1999). State-of-the-Art lecture. Role of angiotensin and oxidative stress in essential hypertension. Hypertension 34, 943–949. 10.1161/01.hyp.34.4.943.

22. Lerman, L.O., Kurtz, T.W., Touyz, R.M., Ellison, D.H., Chade, A.R., Crowley, S.D., Mattson, D.L., Mullins, J.J., Osborn, J., Eirin, A., et al. (2019). Animal Models of Hypertension: A Scientific Statement From the American Heart Association. Hypertension 73, e87–e120. 10.1161/hyp.0000000000000090.

23. Kawada, N., Imai, E., Karber, A., Welch, W.J., and Wilcox, C.S. (2002). A mouse model of angiotensin II slow pressor response: role of oxidative stress. J Am Soc Nephrol 13, 2860–2868. 10.1097/01.asn.0000035087.11758.ed.

24. Young, C.N., Cao, X., Guruju, M.R., Pierce, J.P., Morgan, D.A., Wang, G., Iadecola, C., Mark, A.L., and Davisson, R.L. (2012). ER stress in the brain subfornical organ mediates angiotensin-dependent hypertension. J Clin Invest 122, 3960–3964. 10.1172/jci64583.

25. Dickinson, C.J., and Lawrence, J.R. (1963). A slowly developing pressor response to small concentrations of angiotensin. Its bearing on the pathogenesis of chronic renal hypertension. Lancet 1, 1354–1356. 10.1016/s0140-6736(63)91929-9.

26. Capone, C., Faraco, G., Peterson, J.R., Coleman, C., Anrather, J., Milner, T.A., Pickel, V.M., Davisson, R.L., and Iadecola, C. (2012). Central cardiovascular circuits contribute to the neurovascular dysfunction in angiotensin II hypertension. The Journal of neuroscience : the official journal of the Society for Neuroscience 32. 10.1523/JNEUROSCI.6262-11.2012.

27. Chen, S. (2012). Essential hypertension: perspectives and future directions. J Hypertens 30, 42–45. 10.1097/HJH.0b013e32834ee23c.

28. van den Kerkhof, M., de Jong, J.J.A., Voorter, P.H.M., Postma, A.A., Kroon, A.A., van Oostenbrugge, R.J., Jansen, J.F.A., and Backes, W.H. (2024). Blood-Brain Barrier Integrity Decreases With Higher Blood Pressure: A 7T DCE-MRI Study. Hypertension 81, 2162–2172. 10.1161/hypertensionaha.123.22617.

29. Junejo, R.T., Braz, I.D., Lucas, S.J., van Lieshout, J.J., Phillips, A.A., Lip, G.Y., and Fisher, J.P. (2020). Neurovascular coupling and cerebral autoregulation in atrial fibrillation. J Cereb Blood Flow Metab 40, 1647–1657. 10.1177/0271678x19870770.

30. Yang, S., and Webb, A.J.S. (2024). Reduced neurovascular coupling is associated with increased cardiovascular risk without established cerebrovascular disease: A cross-sectional analysis in UK biobank. J Cereb Blood Flow Metab, 271678x241302172. 10.1177/0271678x241302172.

31. Faraco, G., Hochrainer, K., Segarra, S.G., Schaeffer, S., Santisteban, M.M., Menon, A., Jiang, H., Holtzman, D.M., Anrather, J., and Iadecola, C. (2019). Dietary salt promotes cognitive impairment through tau phosphorylation. Nature 574, 686–690. 10.1038/s41586-019-1688-z.

32. Faraco, G., Sugiyama, Y., Lane, D., Garcia-Bonilla, L., Chang, H., Santisteban, M.M., Racchumi, G., Murphy, M., Van Rooijen, N., Anrather, J., and Iadecola, C. (2016). Perivascular macrophages mediate the neurovascular and cognitive dysfunction associated with hypertension. The Journal of Clinical Investigation 126, 4674–4689. 10.1172/JCI86950.

33. Capone, C., Faraco, G., Park, L., Cao, X., Davisson, R.L., and Iadecola, C. (2011). The cerebrovascular dysfunction induced by slow pressor doses of angiotensin II precedes the development of hypertension. American journal of physiology. Heart and circulatory physiology 300. 10.1152/ajpheart.00679.2010.

34. Girouard, H., Park, L., Anrather, J., Zhou, P., and Iadecola, C. (2007). Cerebrovascular nitrosative stress mediates neurovascular and endothelial dysfunction induced by angiotensin II. Arterioscler Thromb Vasc Biol 27, 303–309. 10.1161/01.Atv.0000253885.41509.25.

35. Iadecola, C., Smith, E.E., Anrather, J., Gu, C., Mishra, A., Misra, S., Perez-Pinzon, M.A., Shih, A.Y., Sorond, F.A., van Veluw, S.J., and Wellington, C.L. (2023). The Neurovasculome: Key Roles in Brain Health and Cognitive Impairment: A Scientific Statement From the American Heart Association/American Stroke Association. Stroke 54, e251–e271. 10.1161/str.0000000000000431.

36. Garcia, F.J., Sun, N., Lee, H., Godlewski, B., Mathys, H., Galani, K., Zhou, B., Jiang, X., Ng, A.P., Mantero, J., et al. (2022). Single-cell dissection of the human brain vasculature. Nature 603, 893–899. 10.1038/s41586-022-04521-7.

37. Wälchli, T., Ghobrial, M., Schwab, M., Takada, S., Zhong, H., Suntharalingham, S., Vetiska, S., Gonzalez, D.R., Wu, R., Rehrauer, H., et al. (2024). Single-cell atlas of the human brain vasculature across development, adulthood and disease. Nature 632, 603–613. 10.1038/s41586-024-07493-y.

38. Yang, A.C., Vest, R.T., Kern, F., Lee, D.P., Agam, M., Maat, C.A., Losada, P.M., Chen, M.B., Schaum, N., Khoury, N., et al. (2022). A human brain vascular atlas reveals diverse mediators of Alzheimer’s risk. Nature 603, 885–892. 10.1038/s41586-021-04369-3.

39. Vanlandewijck, M., He, L., Mäe, M.A., Andrae, J., Ando, K., Del Gaudio, F., Nahar, K., Lebouvier, T., Laviña, B., Gouveia, L., et al. (2018). A molecular atlas of cell types and zonation in the brain vasculature. Nature 554, 475–480. 10.1038/nature25739.

40. Girouard, H., Lessard, A., Capone, C., Milner, T.A., and Iadecola, C. (2008). The neurovascular dysfunction induced by angiotensin II in the mouse neocortex is sexually dimorphic. Am J Physiol Heart Circ Physiol 294, H156–163. 10.1152/ajpheart.01137.2007.

41. Santisteban, M.M., and Iadecola, C. (2025). The pathobiology of neurovascular aging. Neuron 113, 49–70. 10.1016/j.neuron.2024.12.014.

42. Saunders, A., Macosko, E.Z., Wysoker, A., Goldman, M., Krienen, F.M., de Rivera, H., Bien, E., Baum, M., Bortolin, L., Wang, S., et al. (2018). Molecular Diversity and Specializations among the Cells of the Adult Mouse Brain. Cell 174, 1015–1030.e1016. 10.1016/j.cell.2018.07.028.

43. Macosko, E.Z., Basu, A., Satija, R., Nemesh, J., Shekhar, K., Goldman, M., Tirosh, I., Bialas, A.R., Kamitaki, N., Martersteck, E.M., et al. (2015). Highly Parallel Genome-wide Expression Profiling of Individual Cells Using Nanoliter Droplets. Cell 161, 1202–1214. 10.1016/j.cell.2015.05.002.

44. Garcia-Bonilla, L., Shahanoor, Z., Sciortino, R., Nazarzoda, O., Racchumi, G., Iadecola, C., and Anrather, J. (2024). Analysis of brain and blood single-cell transcriptomics in acute and subacute phases after experimental stroke. Nat Immunol 25, 357–370. 10.1038/s41590-023-01711-x.

45. Chen, M.B., Yang, A.C., Yousef, H., Lee, D., Chen, W., Schaum, N., Lehallier, B., Quake, S.R., and Wyss-Coray, T. (2020). Brain Endothelial Cells Are Exquisite Sensors of Age-Related Circulatory Cues. Cell Reports 30, 4418–4432.e4414. 10.1016/j.celrep.2020.03.012.

46. Vanlandewijck, M., He, L., Mae, M.A., Andrae, J., Ando, K., Del Gaudio, F., Nahar, K., Lebouvier, T., Lavina, B., Gouveia, L., et al. (2018). A molecular atlas of cell types and zonation in the brain vasculature. Nature 554, 475–480. 10.1038/nature25739.

47. Kalucka, J., de Rooij, L., Goveia, J., Rohlenova, K., Dumas, S.J., Meta, E., Conchinha, N.V., Taverna, F., Teuwen, L.A., Veys, K., et al. (2020). Single-Cell Transcriptome Atlas of Murine Endothelial Cells. Cell 180, 764–779.e720. 10.1016/j.cell.2020.01.015.

48. Ghandour, M.S., Langley, O.K., Zhu, X.L., Waheed, A., and Sly, W.S. (1992). Carbonic anhydrase IV on brain capillary endothelial cells: a marker associated with the blood-brain barrier. Proc Natl Acad Sci U S A 89, 6823–6827. 10.1073/pnas.89.15.6823.

49. Zhao, L., Li, Z., Vong, J.S.L., Chen, X., Lai, H.-M., Yan, L.Y.C., Huang, J., Sy, S.K.H., Tian, X., Huang, Y., et al. (2020). Pharmacologically reversible zonation-dependent endothelial cell transcriptomic changes with neurodegenerative disease associations in the aged brain. Nature Communications 11, 4413. 10.1038/s41467-020-18249-3.

50. Yang, A.C., Stevens, M.Y., Chen, M.B., Lee, D.P., Stähli, D., Gate, D., Contrepois, K., Chen, W., Iram, T., Zhang, L., et al. (2020). Physiological blood–brain transport is impaired with age by a shift in transcytosis. Nature 583, 425–430. 10.1038/s41586-020-2453-z.

51. Tran, K.A., Zhang, X., Predescu, D., Huang, X., Machado, R.F., Göthert, J.R., Malik, A.B., Valyi-Nagy, T., and Zhao, Y.Y. (2016). Endothelial β-Catenin Signaling Is Required for Maintaining Adult Blood-Brain Barrier Integrity and Central Nervous System Homeostasis. Circulation 133, 177–186. 10.1161/circulationaha.115.015982.

52. Cui, Y., Wang, Y., Song, X., Ning, H., Zhang, Y., Teng, Y., Wang, J., and Yang, X. (2021). Brain endothelial PTEN/AKT/NEDD4-2/MFSD2A axis regulates blood-brain barrier permeability. Cell Rep 36, 109327. 10.1016/j.celrep.2021.109327.

53. Andreone, B.J., Chow, B.W., Tata, A., Lacoste, B., Ben-Zvi, A., Bullock, K., Deik, A.A., Ginty, D.D., Clish, C.B., and Gu, C. (2017). Blood-Brain Barrier Permeability Is Regulated by Lipid Transport-Dependent Suppression of Caveolae-Mediated Transcytosis. Neuron 94, 581–594.e585. 10.1016/j.neuron.2017.03.043.

54. Wiley, C.D., and Campisi, J. (2021). The metabolic roots of senescence: mechanisms and opportunities for intervention. Nature Metabolism 3, 1290–1301. 10.1038/s42255-021-00483-8.

55. Luo, X., Jiang, X., Li, J., Bai, Y., Li, Z., Wei, P., Sun, S., Liang, Y., Han, S., Li, X., and Zhang, B. (2019). Insulin-like growth factor-1 attenuates oxidative stress-induced hepatocyte premature senescence in liver fibrogenesis via regulating nuclear p53–progerin interaction. Cell Death & Disease 10, 451. 10.1038/s41419-019-1670-6.

56. Xiang, Y., You, Z., Huang, X., Dai, J., Zhang, J., Nie, S., Xu, L., Jiang, J., and Xu, J. (2022). Oxidative stress-induced premature senescence and aggravated denervated skeletal muscular atrophy by regulating progerin-p53 interaction. Skelet Muscle 12, 19. 10.1186/s13395-022-00302-y.

57. López-Otín, C., Blasco, M.A., Partridge, L., Serrano, M., and Kroemer, G. (2023). Hallmarks of aging: An expanding universe. Cell 186, 243–278. 10.1016/j.cell.2022.11.001.

58. Hemming, M.L., Selkoe, D.J., and Farris, W. (2007). Effects of prolonged angiotensin-converting enzyme inhibitor treatment on amyloid beta-protein metabolism in mouse models of Alzheimer disease. Neurobiology of Disease 26, 273–281. 10.1016/j.nbd.2007.01.004.

59. Cuddy, L.K., Prokopenko, D., Cunningham, E.P., Brimberry, R., Song, P., Kirchner, R., Chapman, B.A., Hofmann, O., Hide, W., Procissi, D., et al. (2020). Aβ-accelerated neurodegeneration caused by Alzheimer’s-associated ACE variant R1279Q is rescued by angiotensin system inhibition in mice. Sci Transl Med 12. 10.1126/scitranslmed.aaz2541.

60. Yousef, H., Czupalla, C.J., Lee, D., Chen, M.B., Burke, A.N., Zera, K.A., Zandstra, J., Berber, E., Lehallier, B., Mathur, V., et al. (2019). Aged blood impairs hippocampal neural precursor activity and activates microglia via brain endothelial cell VCAM1. Nat Med 25, 988–1000. 10.1038/s41591-019-0440-4.

61. Chamling, X., Kallman, A., Fang, W., Berlinicke, C.A., Mertz, J.L., Devkota, P., Pantoja, I.E.M., Smith, M.D., Ji, Z., Chang, C., et al. (2021). Single-cell transcriptomic reveals molecular diversity and developmental heterogeneity of human stem cell-derived oligodendrocyte lineage cells. Nature Communications 12, 652. 10.1038/s41467-021-20892-3.

62. Marques, S., Zeisel, A., Codeluppi, S., van Bruggen, D., Mendanha Falcão, A., Xiao, L., Li, H., Häring, M., Hochgerner, H., Romanov, R.A., et al. (2016). Oligodendrocyte heterogeneity in the mouse juvenile and adult central nervous system. Science 352, 1326–1329. 10.1126/science.aaf6463.

63. Thorburne, S.K., and Juurlink, B.H.J. (1996). Low Glutathione and High Iron Govern the Susceptibility of Oligodendroglial Precursors to Oxidative Stress. Journal of Neurochemistry 67, 1014–1022. 10.1046/j.1471-4159.1996.67031014.x.

64. Tse, K.H., and Herrup, K. (2017). DNA damage in the oligodendrocyte lineage and its role in brain aging. Mech Ageing Dev 161, 37–50. 10.1016/j.mad.2016.05.006.

65. French, H.M., Reid, M., Mamontov, P., Simmons, R.A., and Grinspan, J.B. (2009). Oxidative stress disrupts oligodendrocyte maturation. J Neurosci Res 87, 3076–3087. 10.1002/jnr.22139.

66. Kotter, M.R., Stadelmann, C., and Hartung, H.-P. (2011). Enhancing remyelination in disease—can we wrap it up? Brain 134, 1882–1900. 10.1093/brain/awr014.

67. Tawk, M., Makoukji, J., Belle, M., Fonte, C., Trousson, A., Hawkins, T., Li, H., Ghandour, S., Schumacher, M., and Massaad, C. (2011). Wnt/beta-catenin signaling is an essential and direct driver of myelin gene expression and myelinogenesis. J Neurosci 31, 3729–3742. 10.1523/jneurosci.4270-10.2011.

68. Weider, M., Starost, L.J., Groll, K., Küspert, M., Sock, E., Wedel, M., Fröb, F., Schmitt, C., Baroti, T., Hartwig, A.C., et al. (2018). Nfat/calcineurin signaling promotes oligodendrocyte differentiation and myelination by transcription factor network tuning. Nature Communications 9, 899. 10.1038/s41467-018-03336-3.

69. Kinner, A., Wu, W., Staudt, C., and Iliakis, G. (2008). Gamma-H2AX in recognition and signaling of DNA double-strand breaks in the context of chromatin. Nucleic Acids Res 36, 5678–5694. 10.1093/nar/gkn550.

70. Beiter, R.M., Rivet-Noor, C., Merchak, A.R., Bai, R., Johanson, D.M., Slogar, E., Sol-Church, K., Overall, C.C., and Gaultier, A. (2022). Evidence for oligodendrocyte progenitor cell heterogeneity in the adult mouse brain. Sci Rep 12, 12921. 10.1038/s41598-022-17081-7.

71. Hughes, A.N., and Appel, B. (2019). Oligodendrocytes express synaptic proteins that modulate myelin sheath formation. Nat Commun 10, 4125. 10.1038/s41467-019-12059-y.

72. Dimas, P., Montani, L., Pereira, J.A., Moreno, D., Trötzmüller, M., Gerber, J., Semenkovich, C.F., Köfeler, H.C., and Suter, U. (2019). CNS myelination and remyelination depend on fatty acid synthesis by oligodendrocytes. eLife 8, e44702. 10.7554/eLife.44702.

73. Sandra, G., Jan, H.O., Robert, K., Ingo, H., Ingo, B., Susanne, W., Sven, P.W., Wiebke, M., Xin, L., Corinna, L.-S., et al. (2010). Elevated Phosphatidylinositol 3,4,5-Trisphosphate in Glia Triggers Cell-Autonomous Membrane Wrapping and Myelination. The Journal of Neuroscience 30, 8953. 10.1523/JNEUROSCI.0219-10.2010.

74. Karlie, N.F.-S., and Bruce, A. (2021). The Akt-mTOR Pathway Drives Myelin Sheath Growth by Regulating Cap-Dependent Translation. The Journal of Neuroscience 41, 8532. 10.1523/JNEUROSCI.0783-21.2021.

75. Liu, A., Li, J., Marin-Husstege, M., Kageyama, R., Fan, Y., Gelinas, C., and Casaccia-Bonnefil, P. (2006). A molecular insight of Hes5-dependent inhibition of myelin gene expression: old partners and new players. Embo j 25, 4833–4842. 10.1038/sj.emboj.7601352.

76. Petersen, M.A., Ryu, J.K., Chang, K.-J., Etxeberria, A., Bardehle, S., Mendiola, A.S., Kamau-Devers, W., Fancy, S.P.J., Thor, A., Bushong, E.A., et al. (2017). Fibrinogen Activates BMP Signaling in Oligodendrocyte Progenitor Cells and Inhibits Remyelination after Vascular Damage. Neuron 96, 1003–1012.e1007. papers3://publication/doi/10.1016/j.neuron.2017.10.008.

77. Snaidero, N., and Simons, M. (2017). The logistics of myelin biogenesis in the central nervous system. Glia 65, 1021–1031. 10.1002/glia.23116.

78. Young, K.M., Psachoulia, K., Tripathi, R.B., Dunn, S.J., Cossell, L., Attwell, D., Tohyama, K., and Richardson, W.D. (2013). Oligodendrocyte dynamics in the healthy adult CNS: evidence for myelin remodeling. Neuron 77, 873–885. 10.1016/j.neuron.2013.01.006.

79. Gallego-Delgado, P., James, R., Browne, E., Meng, J., Umashankar, S., Tan, L., Picon, C., Mazarakis, N.D., Faisal, A.A., Howell, O.W., and Reynolds, R. (2020). Neuroinflammation in the normal-appearing white matter (NAWM) of the multiple sclerosis brain causes abnormalities at the nodes of Ranvier. PLoS Biol 18, e3001008. 10.1371/journal.pbio.3001008.

80. Crawford, D.K., Mangiardi, M., and Tiwari-Woodruff, S.K. (2009). Assaying the functional effects of demyelination and remyelination: revisiting field potential recordings. J Neurosci Methods 182, 25–33. 10.1016/j.jneumeth.2009.05.013.

81. Südhof, T.C., and Rizo, J. (2011). Synaptic vesicle exocytosis. Cold Spring Harb Perspect Biol 3. 10.1101/cshperspect.a005637.

82. Clayton, E.L., Anggono, V., Smillie, K.J., Chau, N., Robinson, P.J., and Cousin, M.A. (2009). The phospho-dependent dynamin-syndapin interaction triggers activity-dependent bulk endocytosis of synaptic vesicles. J Neurosci 29, 7706–7717. 10.1523/jneurosci.1976-09.2009.

83. Eto, K., Ishibashi, H., Yoshimura, T., Watanabe, M., Miyamoto, A., Ikenaka, K., Moorhouse, A.J., and Nabekura, J. (2012). Enhanced GABAergic activity in the mouse primary somatosensory cortex is insufficient to alleviate chronic pain behavior with reduced expression of neuronal potassium-chloride cotransporter. J Neurosci 32, 16552–16559. 10.1523/jneurosci.2104-12.2012.

84. Sukenik, N., Vinogradov, O., Weinreb, E., Segal, M., Levina, A., and Moses, E. (2021). Neuronal circuits overcome imbalance in excitation and inhibition by adjusting connection numbers. Proc Natl Acad Sci U S A 118. 10.1073/pnas.2018459118.

85. Pfeiffer, P., Egorov, A.V., Lorenz, F., Schleimer, J.H., Draguhn, A., and Schreiber, S. (2020). Clusters of cooperative ion channels enable a membrane-potential-based mechanism for short-term memory. Elife 9. 10.7554/eLife.49974.

86. Vivas, O., Moreno, C.M., Santana, L.F., and Hille, B. (2017). Proximal clustering between BK and Ca(V)1.3 channels promotes functional coupling and BK channel activation at low voltage. Elife 6. 10.7554/eLife.28029.

87. Kann, O., and Kovács, R. (2007). Mitochondria and neuronal activity. Am J Physiol Cell Physiol 292, C641–657. 10.1152/ajpcell.00222.2006.

88. Wang, H., Hitron, I.M., Iadecola, C., and Pickel, V.M. (2005). Synaptic and vascular associations of neurons containing cyclooxygenase-2 and nitric oxide synthase in rat somatosensory cortex. Cereb Cortex 15, 1250–1260. 10.1093/cercor/bhi008.

89. Buerk, D.G., Ances, B.M., Greenberg, J.H., and Detre, J.A. (2003). Temporal dynamics of brain tissue nitric oxide during functional forepaw stimulation in rats. Neuroimage 18, 1–9. 10.1006/nimg.2002.1314.

90. Lourenço, C.F., Santos, R.M., Barbosa, R.M., Cadenas, E., Radi, R., and Laranjinha, J. (2014). Neurovascular coupling in hippocampus is mediated via diffusion by neuronal-derived nitric oxide. Free Radic Biol Med 73, 421–429. 10.1016/j.freeradbiomed.2014.05.021.

91. Ahn, S.J., Anfray, A., Anrather, J., and Iadecola, C. (2023). Calcium transients in nNOS neurons underlie distinct phases of the neurovascular response to barrel cortex activation in awake mice. J Cereb Blood Flow Metab 43, 1633–1647. 10.1177/0271678x231173175.

92. Tasic, B., Menon, V., Nguyen, T.N., Kim, T.K., Jarsky, T., Yao, Z., Levi, B., Gray, L.T., Sorensen, S.A., Dolbeare, T., et al. (2016). Adult mouse cortical cell taxonomy revealed by single cell transcriptomics. Nat Neurosci 19, 335–346. 10.1038/nn.4216.

93. Paul, A., Crow, M., Raudales, R., He, M., Gillis, J., and Huang, Z.J. (2017). Transcriptional Architecture of Synaptic Communication Delineates GABAergic Neuron Identity. Cell 171, 522–539.e520. 10.1016/j.cell.2017.08.032.

94. Sagi, Y., Heiman, M., Peterson, J.D., Musatov, S., Scarduzio, M., Logan, S.M., Kaplitt, M.G., Surmeier, D.J., Heintz, N., and Greengard, P. (2014). Nitric oxide regulates synaptic transmission between spiny projection neurons. Proc Natl Acad Sci U S A 111, 17636–17641. 10.1073/pnas.1420162111.

95. Gomes, J.R., Lobo, A.C., Melo, C.V., Inácio, A.R., Takano, J., Iwata, N., Saido, T.C., de Almeida, L.P., Wieloch, T., and Duarte, C.B. (2011). Cleavage of the vesicular GABA transporter under excitotoxic conditions is followed by accumulation of the truncated transporter in nonsynaptic sites. J Neurosci 31, 4622–4635. 10.1523/jneurosci.3541-10.2011.

96. Armulik, A., Genové, G., Mäe, M., Nisancioglu, M.H., Wallgard, E., Niaudet, C., He, L., Norlin, J., Lindblom, P., Strittmatter, K., et al. (2010). Pericytes regulate the blood-brain barrier. Nature 468, 557–561. 10.1038/nature09522.

97. Dalkara, T., Østergaard, L., Heusch, G., and Attwell, D. (2025). Pericytes in the brain and heart: functional roles and response to ischaemia and reperfusion. Cardiovasc Res 120, 2336–2348. 10.1093/cvr/cvae147.

98. Ramamoorthy, P., and Whim, M.D. (2008). Trafficking and fusion of neuropeptide Y-containing dense-core granules in astrocytes. J Neurosci 28, 13815–13827. 10.1523/jneurosci.5361-07.2008.

99. Aronica, E., Gorter, J.A., Ijlst-Keizers, H., Rozemuller, A.J., Yankaya, B., Leenstra, S., and Troost, D. (2003). Expression and functional role of mGluR3 and mGluR5 in human astrocytes and glioma cells: opposite regulation of glutamate transporter proteins. Eur J Neurosci 17, 2106–2118. 10.1046/j.1460-9568.2003.02657.x.

100. Ehmsen, J.T., Ma, T.M., Sason, H., Rosenberg, D., Ogo, T., Furuya, S., Snyder, S.H., and Wolosker, H. (2013). D-serine in glia and neurons derives from 3-phosphoglycerate dehydrogenase. J Neurosci 33, 12464–12469. 10.1523/jneurosci.4914-12.2013.

101. Cheli, V.T., Correale, J., Paez, P.M., and Pasquini, J.M. (2020). Iron Metabolism in Oligodendrocytes and Astrocytes, Implications for Myelination and Remyelination. ASN Neuro 12, 1759091420962681. 10.1177/1759091420962681.

102. Dai, Y., Bi, M., Jiao, Q., Du, X., Yan, C., and Jiang, H. (2024). Astrocyte-derived apolipoprotein D is required for neuronal survival in Parkinson’s disease. NPJ Parkinsons Dis 10, 143. 10.1038/s41531-024-00753-8.

103. Pascua-Maestro, R., González, E., Lillo, C., Ganfornina, M.D., Falcón-Pérez, J.M., and Sanchez, D. (2018). Extracellular Vesicles Secreted by Astroglial Cells Transport Apolipoprotein D to Neurons and Mediate Neuronal Survival Upon Oxidative Stress. Front Cell Neurosci 12, 526. 10.3389/fncel.2018.00526.

104. Thomas, A.L., Lehn, M.A., Janssen, E.M., Hildeman, D.A., and Chougnet, C.A. (2022). Naturally-aged microglia exhibit phagocytic dysfunction accompanied by gene expression changes reflective of underlying neurologic disease. Sci Rep 12, 19471. 10.1038/s41598-022-21920-y.

105. Ungvari, Z., Tarantini, S., Donato, A.J., Galvan, V., and Csiszar, A. (2018). Mechanisms of Vascular Aging. Circ Res 123, 849–867. 10.1161/CIRCRESAHA.118.311378.

106. Gorgoulis, V., Adams, P.D., Alimonti, A., Bennett, D.C., Bischof, O., Bishop, C., Campisi, J., Collado, M., Evangelou, K., Ferbeyre, G., et al. (2019). Cellular Senescence: Defining a Path Forward. Cell 179, 813–827. 10.1016/j.cell.2019.10.005.

107. Kaiser, D., Weise, G., Möller, K., Scheibe, J., Pösel, C., Baasch, S., Gawlitza, M., Lobsien, D., Diederich, K., Minnerup, J., et al. (2014). Spontaneous white matter damage, cognitive decline and neuroinflammation in middle-aged hypertensive rats: an animal model of early-stage cerebral small vessel disease. Acta Neuropathol Commun 2, 169. 10.1186/s40478-014-0169-8.

108. Capone, C., Faraco, G., Park, L., Cao, X., Davisson, R.L., and Iadecola, C. (2010). The cerebrovascular dysfunction induced by slow pressor doses of angiotensin II precedes the development of hypertension. American Journal of Physiology-Heart and Circulatory Physiology 300, H397–H407. 10.1152/ajpheart.00679.2010.

109. Bloom, S.I., Islam, M.T., Lesniewski, L.A., and Donato, A.J. (2023). Mechanisms and consequences of endothelial cell senescence. Nat Rev Cardiol 20, 38–51. 10.1038/s41569-022-00739-0.

110. Mogi, M. (2020). Effect of renin-angiotensin system on senescence. Geriatr Gerontol Int 20, 520–525. 10.1111/ggi.13927.

111. Khan, I., Schmidt, M.O., Kallakury, B., Jain, S., Mehdikhani, S., Levi, M., Mendonca, M., Welch, W., Riegel, A.T., Wilcox, C.S., and Wellstein, A. (2021). Low Dose Chronic Angiotensin II Induces Selective Senescence of Kidney Endothelial Cells. Front Cell Dev Biol 9, 782841. 10.3389/fcell.2021.782841.

112. Guo, S., Deng, W., Xing, C., Zhou, Y., Ning, M., and Lo, E.H. (2019). Effects of aging, hypertension and diabetes on the mouse brain and heart vasculomes. Neurobiology of Disease 126, 117–123. 10.1016/j.nbd.2018.07.021.

113. Scalera, F., Martens-Lobenhoffer, J., Bukowska, A., Lendeckel, U., Täger, M., and Bode-Böger, S.M. (2008). Effect of telmisartan on nitric oxide--asymmetrical dimethylarginine system: role of angiotensin II type 1 receptor gamma and peroxisome proliferator activated receptor gamma signaling during endothelial aging. Hypertension 51, 696–703. 10.1161/hypertensionaha.107.104570.

114. Afsar, B., and Afsar, R.E. (2023). Hypertension and cellular senescence. Biogerontology 24, 457–478. 10.1007/s10522-023-10031-4.

115. Saavedra, J.M. (2005). Brain angiotensin II: new developments, unanswered questions and therapeutic opportunities. Cell Mol Neurobiol 25, 485–512. 10.1007/s10571-005-4011-5.

116. Csik, B., Nyúl-Tóth, Á., Gulej, R., Patai, R., Kiss, T., Delfavero, J., Nagaraja, R.Y., Balasubramanian, P., Shanmugarama, S., Ungvari, A., et al. (2025). Senescent Endothelial Cells in Cerebral Microcirculation Are Key Drivers of Age-Related Blood-Brain Barrier Disruption, Microvascular Rarefaction, and Neurovascular Coupling Impairment in Mice. Aging Cell, e70048. 10.1111/acel.70048.

117. Groh, J., and Simons, M. (2025). White matter aging and its impact on brain function. Neuron 113, 127–139. 10.1016/j.neuron.2024.10.019.

118. Pak, K., Chan, S.L., and Mattson, M.P. (2003). Presenilin-1 mutation sensitizes oligodendrocytes to glutamate and amyloid toxicities, and exacerbates white matter damage and memory impairment in mice. Neuromolecular Med 3, 53–64. 10.1385/nmm:3:1:53.

119. Desai, M.K., Sudol, K.L., Janelsins, M.C., Mastrangelo, M.A., Frazer, M.E., and Bowers, W.J. (2009). Triple-transgenic Alzheimer’s disease mice exhibit region-specific abnormalities in brain myelination patterns prior to appearance of amyloid and tau pathology. Glia 57, 54–65. 10.1002/glia.20734.

120. Desai, M.K., Guercio, B.J., Narrow, W.C., and Bowers, W.J. (2011). An Alzheimer’s disease-relevant presenilin-1 mutation augments amyloid-beta-induced oligodendrocyte dysfunction. Glia 59, 627–640. 10.1002/glia.21131.

121. Reimer, M.M., McQueen, J., Searcy, L., Scullion, G., Zonta, B., Desmazieres, A., Holland, P.R., Smith, J., Gliddon, C., Wood, E.R., et al. (2011). Rapid disruption of axon-glial integrity in response to mild cerebral hypoperfusion. J Neurosci 31, 18185–18194. 10.1523/jneurosci.4936-11.2011.

122. Maillard, P., Seshadri, S., Beiser, A., Himali, J.J., Au, R., Fletcher, E., Carmichael, O., Wolf, P.A., and DeCarli, C. (2012). Effects of systolic blood pressure on white-matter integrity in young adults in the Framingham Heart Study: a cross-sectional study. Lancet Neurol 11, 1039–1047. 10.1016/s1474-4422(12)70241-7.

123. Won, J., Maillard, P., Shan, K., Ashley, J., Cardim, D., Zhu, D.C., and Zhang, R. (2024). Association of Blood Pressure With Brain White Matter Microstructural Integrity Assessed With MRI Diffusion Tensor Imaging in Healthy Young Adults. Hypertension 81, 1145–1155. 10.1161/hypertensionaha.123.22337.

124. Petrea, R.E., Pinheiro, A., Demissie, S., Ekenze, O., Aparicio, H.J., Satizabal, C.L., Maillard, P., DeCarli, C., Beiser, A.S., Seshadri, S., et al. (2024). Hypertension Trends and White Matter Brain Injury in the Offspring Framingham Heart Study Cohort. Hypertension 81, 87–95. 10.1161/hypertensionaha.123.21264.

125. Dufouil, C., de Kersaint-Gilly, A., Besançon, V., Levy, C., Auffray, E., Brunnereau, L., Alpérovitch, A., and Tzourio, C. (2001). Longitudinal study of blood pressure and white matter hyperintensities: the EVA MRI Cohort. Neurology 56, 921–926. 10.1212/wnl.56.7.921.

126. Marcus, J., Gardener, H., Rundek, T., Elkind, M.S., Sacco, R.L., Decarli, C., and Wright, C.B. (2011). Baseline and longitudinal increases in diastolic blood pressure are associated with greater white matter hyperintensity volume: the northern Manhattan study. Stroke 42, 2639–2641. 10.1161/strokeaha.111.617571.

127. Bruno, C., Xin-Kang, T., Armelle, R., Nella, S., Bertrand, L., Jean, R., and Edith, H. (2004). Cortical GABA Interneurons in Neurovascular Coupling: Relays for Subcortical Vasoactive Pathways. The Journal of Neuroscience 24, 8940. 10.1523/JNEUROSCI.3065-04.2004.

128. Ayata, C., Ma, J., Meng, W., Huang, P., and Moskowitz, M.A. (1996). L-NA-sensitive rCBF augmentation during vibrissal stimulation in type III nitric oxide synthase mutant mice. Journal of Cerebral Blood Flow & Metabolism 16, 539–541.

129. Ng, S.K., Hauser, W.A., Brust, J.C., and Susser, M. (1993). Hypertension and the risk of new-onset unprovoked seizures. Neurology 43, 425–428. 10.1212/wnl.43.2.425.

130. Stefanidou, M., Himali, J.J., Devinsky, O., Romero, J.R., Ikram, M.A., Beiser, A.S., Seshadri, S., and Friedman, D. (2022). Vascular risk factors as predictors of epilepsy in older age: The Framingham Heart Study. Epilepsia 63, 237–243. 10.1111/epi.17108.

131. Xia, M., Wang, T., Wang, Y., Hu, T., Chen, D., and Wang, B. (2024). A neural perspective on the treatment of hypertension: the neurological network excitation and inhibition (E/I) imbalance in hypertension. Front Cardiovasc Med 11, 1436059. 10.3389/fcvm.2024.1436059.

132. Acosta, J.C., Banito, A., Wuestefeld, T., Georgilis, A., Janich, P., Morton, J.P., Athineos, D., Kang, T.-W., Lasitschka, F., Andrulis, M., et al. (2013). A complex secretory program orchestrated by the inflammasome controls paracrine senescence. Nature Cell Biology 15, 978–990. 10.1038/ncb2784.

133. Coppé, J.P., Desprez, P.Y., Krtolica, A., and Campisi, J. (2010). The senescence-associated secretory phenotype: the dark side of tumor suppression. Annu Rev Pathol 5, 99–118. 10.1146/annurev-pathol-121808-102144.

134. Wang, B., Han, J., Elisseeff, J.H., and Demaria, M. (2024). The senescence-associated secretory phenotype and its physiological and pathological implications. Nat Rev Mol Cell Biol 25, 958–978. 10.1038/s41580-024-00727-x.

135. Catt, K.J., Cran, E., Zimmet, P.Z., Best, J.B., Cain, M.D., and Coghlan, J.P. (1971). Angiotensin II blood-levels in human hypertension. Lancet 1, 459–464. 10.1016/s0140-6736(71)91085-3.

136. Jones, D.W., Ferdinand, K.C., Taler, S.J., Johnson, H.M., Shimbo, D., Abdalla, M., Altieri, M.M., Bansal, N., Bello, N.A., Bress, A.P., et al. (2025). 2025 AHA/ACC/AANP/AAPA/ABC/ACCP/ACPM/AGS/AMA/ASPC/NMA/PCNA/SGIM Guideline for the Prevention, Detection, Evaluation and Management of High Blood Pressure in Adults: A Report of the American College of Cardiology/American Heart Association Joint Committee on Clinical Practice Guidelines. Hypertension. 10.1161/hyp.0000000000000249.

137. Barton, M., and Meyer, M.R. (2009). Postmenopausal hypertension: mechanisms and therapy. Hypertension 54, 11–18. 10.1161/hypertensionaha.108.120022.

138. Faraco, G., Brea, D., Garcia-Bonilla, L., Wang, G., Racchumi, G., Chang, H., Buendia, I., Santisteban, M.M., Segarra, S.G., Koizumi, K., et al. (2018). Dietary salt promotes neurovascular and cognitive dysfunction through a gut-initiated TH17 response. Nat Neurosci 21, 240–249. 10.1038/s41593-017-0059-z.

139. Datlinger, P., Rendeiro, A.F., Schmidl, C., Krausgruber, T., Traxler, P., Klughammer, J., Schuster, L.C., Kuchler, A., Alpar, D., and Bock, C. (2017). Pooled CRISPR screening with single-cell transcriptome readout. Nature Methods 14, 297–301. 10.1038/nmeth.4177.

140. Dobin, A., Davis, C.A., Schlesinger, F., Drenkow, J., Zaleski, C., Jha, S., Batut, P., Chaisson, M., and Gingeras, T.R. (2013). STAR: ultrafast universal RNA-seq aligner. Bioinformatics 29, 15–21. 10.1093/bioinformatics/bts635.

141. McGinnis, C.S., Murrow, L.M., and Gartner, Z.J. (2019). DoubletFinder: Doublet Detection in Single-Cell RNA Sequencing Data Using Artificial Nearest Neighbors. Cell Syst 8, 329–337.e324. 10.1016/j.cels.2019.03.003.

142. Stuart, T., Butler, A., Hoffman, P., Hafemeister, C., Papalexi, E., Mauck, W.M., 3rd, Hao, Y., Stoeckius, M., Smibert, P., and Satija, R. (2019). Comprehensive Integration of Single-Cell Data. Cell 177, 1888–1902.e1821. 10.1016/j.cell.2019.05.031.

143. Melville}, L.M.a.J.H.a.J. (2018). {UMAP: Uniform Manifold Approximation and Projection for Dimension Reduction}. {1802.03426} (stat.ML).

144. Finak, G., McDavid, A., Yajima, M., Deng, J., Gersuk, V., Shalek, A.K., Slichter, C.K., Miller, H.W., McElrath, M.J., Prlic, M., et al. (2015). MAST: a flexible statistical framework for assessing transcriptional changes and characterizing heterogeneity in single-cell RNA sequencing data. Genome Biology 16, 278. 10.1186/s13059-015-0844-5.

145. Almanzar, N., Antony, J., Baghel, A.S., Bakerman, I., Bansal, I., Barres, B.A., Beachy, P.A., Berdnik, D., Bilen, B., Brownfield, D., et al. (2020). A single-cell transcriptomic atlas characterizes ageing tissues in the mouse. Nature 583, 590–595. 10.1038/s41586-020-2496-1.

146. Kalucka, J.,, de Rooij, L.P.M.H.,, Goveia, J.,, Rohlenova, K.,, Dumas, S.J.,, et al. (2020). Single-Cell Transcriptome Atlas of Murine Endothelial Cells. 180, 764–779.e720. 10.1016/j.cell.2020.01.015.

147. Marques, S., Zeisel, A., Codeluppi, S., van Bruggen, D., Mendanha Falcão, A., Xiao, L., Li, H., Häring, M., Hochgerner, H., Romanov, R.A., et al. (2016). Oligodendrocyte heterogeneity in the mouse juvenile and adult central nervous system. Science 352, 1326–1329. 10.1126/science.aaf6463.

148. Li, Q., Zhou, B., Su, M., Liao, P., Lei, F., Li, X., Liao, D., Zhang, X., and Jiang, R. (2023). Visualization and Characterization of the Brain Regional Heterogeneity of Astrocyte-Astrocyte Structural Interactions by Using Improved Iontophoresis with Dual-Fluorescent Dyes. Brain Sci 13. 10.3390/brainsci13121644.

149. McCarthy, D.J., Campbell, K.R., Lun, A.T., and Wills, Q.F. (2017). Scater: pre-processing, quality control, normalization and visualization of single-cell RNA-seq data in R. Bioinformatics 33, 1179–1186. 10.1093/bioinformatics/btw777.

150. Ritchie, M.E., Phipson, B., Wu, D., Hu, Y., Law, C.W., Shi, W., and Smyth, G.K. (2015). limma powers differential expression analyses for RNA-sequencing and microarray studies. Nucleic Acids Res 43, e47. 10.1093/nar/gkv007.

151. Robinson, M.D., McCarthy, D.J., and Smyth, G.K. (2010). edgeR: a Bioconductor package for differential expression analysis of digital gene expression data. Bioinformatics 26, 139–140. 10.1093/bioinformatics/btp616.

152. Squair, J.W., Gautier, M., Kathe, C., Anderson, M.A., James, N.D., Hutson, T.H., Hudelle, R., Qaiser, T., Matson, K.J.E., Barraud, Q., et al. (2021). Confronting false discoveries in single-cell differential expression. Nat Commun 12, 5692. 10.1038/s41467-021-25960-2.

153. Zhang, B., Kirov, S., and Snoddy, J. (2005). WebGestalt: an integrated system for exploring gene sets in various biological contexts. Nucleic Acids Res 33, W741–748. 10.1093/nar/gki475.

154. Hayamizu, T.F., Mangan, M., Corradi, J.P., Kadin, J.A., and Ringwald, M. (2005). The Adult Mouse Anatomical Dictionary: a tool for annotating and integrating data. Genome Biol 6, R29. 10.1186/gb-2005-6-3-r29.

155. Kim, H., de Jesus, A.A., Brooks, S.R., Liu, Y., Huang, Y., VanTries, R., Montealegre Sanchez, G.A., Rotman, Y., Gadina, M., and Goldbach-Mansky, R. (2018). Development of a Validated Interferon Score Using NanoString Technology. J Interferon Cytokine Res 38, 171–185. 10.1089/jir.2017.0127.

156. Park, L., Hochrainer, K., Hattori, Y., Ahn, S.J., Anfray, A., Wang, G., Uekawa, K., Seo, J., Palfini, V., Blanco, I., et al. (2020). Tau induces PSD95-neuronal NOS uncoupling and neurovascular dysfunction independent of neurodegeneration. Nat Neurosci 23, 1079–1089. 10.1038/s41593-020-0686-7.

157. Coleman, C.G., Wang, G., Park, L., Anrather, J., Delagrammatikas, G.J., Chan, J., Zhou, J., Iadecola, C., and Pickel, V.M. (2010). Chronic intermittent hypoxia induces NMDA receptor-dependent plasticity and suppresses nitric oxide signaling in the mouse hypothalamic paraventricular nucleus. J Neurosci 30, 12103–12112. 10.1523/jneurosci.3367-10.2010.

158. Wang, G., Coleman, C.G., Chan, J., Faraco, G., Marques-Lopes, J., Milner, T.A., Guruju, M.R., Anrather, J., Davisson, R.L., Iadecola, C., and Pickel, V.M. (2013). Angiotensin II slow-pressor hypertension enhances NMDA currents and NOX2-dependent superoxide production in hypothalamic paraventricular neurons. Am J Physiol Regul Integr Comp Physiol 304, R1096–1106. 10.1152/ajpregu.00367.2012.

159. Koizumi, K., Hattori, Y., Ahn, S.J., Buendia, I., Ciacciarelli, A., Uekawa, K., Wang, G., Hiller, A., Zhao, L., Voss, H.U., et al. (2018). Apoε4 disrupts neurovascular regulation and undermines white matter integrity and cognitive function. Nat Commun 9, 3816. 10.1038/s41467-018-06301-2.

160. von Bohlen und Halbach, O. (2003). Nitric oxide imaging in living neuronal tissues using fluorescent probes. Nitric Oxide 9, 217–228. 10.1016/j.niox.2004.01.001.

