## Supplementary material for "Hypertension-Induced Neurovascular and Cognitive Dysfunction at Single-Cell Resolution": Key Resources Table

| REAGENT or RESOURCE | SOURCE | IDENTIFIER |
|---|---|---|
| **Antibodies** | | |
| anti-P27 (1:400, rabbit host) – used for detection of endothelial senescence | Proteintech | Cat# 25614-1-AP |
| anti-P21 (1:200, rabbit host) – used for detection of endothelial senescence | Abcam | Cat# ab188224 |
| anti-P16 (1:100, rabbit host) – used for detection of endothelial senescence | Abcam | Cat# ab211542 |
| anti-PECAM-1, clone 2H8 (1:100, goat host) – used to identify endothelial cells | Millipore-Sigma | Cat# MAB1398Z |
| anti-Olig2 (1:200, rabbit host) – used to label oligodendrocyte precursor cells | Abcam | Cat# ab109186 |
| anti-Pdgfrα (1:200, goat host) – used to label oligodendrocyte precursor cells | Biotechne | Cat# AF1062 |
| anti-phospho(Ser$^{139}$)-histone H2A.X (1:100, mouse host) – used to label double-stranded DNA breaks | ThermoFisher Scientific | Cat# MA1-2022 |
| anti-collagen IV (1:400, rabbit host) – used to label collagen IV associated with the endothelium | Abcam | Cat# ab6586 |
| AF-488 donkey anti-rabbit (1:200) | ThermoFisher Scientific | Cat# A-21206 |
| AF-647 donkey anti-goat (1:200) | ThermoFisher Scientific | Cat# A-21447 |
| AF-647 goat anti-Armenian hamster (1:200) | ThermoFisher Scientific | Cat# A78967 |
| AF-488 donkey anti-mouse (1:200) | ThermoFisher Scientific | Cat# A-21202 |
| Cy3 donkey anti-rabbit (1:200) | Sigma-Aldrich | Cat# AP182C |
| **Biological samples** | | |
| C56BL/6J mouse brain samples (cortex) – live cells for single-cell RNA sequencing | This paper | N/A |
| C56BL/6J mouse brain samples (cortex) – frozen tissue for immunofluorescence and histology | This paper | N/A |
| C56BL/6J mouse brain samples (cortex) – live dissociated neurons for NO determination | This paper | N/A |
| C56BL/6J mouse brain samples (cortex) – live slices for trans-callosal recordings | This paper | N/A |
| **Chemicals, peptides, and recombinant proteins** | | |
| Angiotensin II human | Sigma-Aldrich | Cat# A9525; CAS 4474-91-3 |
| Phenylephrine hydrochloride | Millipore-Sigma | Cat# P6126; CAS 61-76-7 |
| Losartan potassium | AK Scientific | Cat# I934; CAS 124750-99-8 |
| Protease VIII | Sigma-Aldrich | Cat# P5380; CAS Cat# 9014-01-1 |
| Papain from papaya latex | Millipore-Sigma | Cat# P4762; CAS 9001-73-4 |
| L-Cysteine | Sigma-Aldrich | Cat# 168149; CAS 52-90-4 |
| Trypsin inhibitor | Sigma-Aldrich | Cat# T6522; CAS 9035-81-8 |
| EDTA and ovomucoid protease inhibitor | Worthington | Cat# LK003153; CAS 9048-46-8 and CAS 9001-73-4 |

| | | |
|---|---|---|
| EthD-1 | Thermo Fisher Scientific | Cat# L-3224 |
| Calcein-AM | Thermo Fisher Scientific | Cat# L-3224 |
| Hoechst 33342 | Thermo Fisher Scientific | Cat# 62249 |
| Guanidine HCL | Thermofisher Scientific | Cat# 24110; |
| Ficoll PM-400 | Sigma-Aldrich | Cat# F4375; CAS 26873-85-8 |
| Sarkosyl | Sigma-Aldrich | Cat# 61747; CAS 137-16-6 |
| EDTA | Thermofisher Scientific | Cat# 17892 |
| DTT | Sigma-Aldrich | Cat# D0632; CAS 3483-12-3 |
| Tris pH 7.5 | Sigma-Aldrich | Cat# DB0339 |
| Droplet generation oil | Bio-Rad | Cat# 1863005 |
| SSC Buffer, 20X, Molecular Grade | Promega | Cat# V4261 |
| perfluorooctanol | Sigma-Aldrich | Cat# 370533; CAS 647-42-7 |
| Epredia™ M-1 Embedding Matrix | Fisher Scientific | Cat# 1310 |
| Pronase | Millipore-Sigma | Cat# 10165921001 |
| Thermolysin | Millipore-Sigma | Cat# P1512; CAS 9073-78-3 |
| **Critical commercial assays** | | |
| RNAscope Multiplex Fluorescent Kit v2 | ACD-Bio-Techne | Cat# 323110 |
| Cellular Senescence Detection Kit | Cell Bio Labs | Cat# CBA-230 |
| DAF-FM (4-amino-5-methylamino-2′,7′-difluorofluorescein) diacetate | Thermo Fisher Scientific | Cat# D23844 |
| **Deposited data** | | |
| Raw scRNA data | This paper | GEO: GSE294795 |
| Analyzed scRNA data | This paper | Tables S1-3 |
| Raw immunofluorescence/histology/behavior data | This paper | Tabel S4 |
| **Experimental models: Organisms/strains** | | |
| Mouse: C57BL/6J: strain no. 000664 | Jackson | IMSR_JAX:000664 |
| Mouse: C57BL/6-Nos1$^{Cre}$: B6.129-*Nos1*$^{tm1(cre)Mgmj}$/J: strain no. 017526 | Jackson | IMSR_JAX:017526 |
| **Software and algorithms** | | |
| R (version 4.5.1) | R Project for Statistical Computing | http://www.r-project.org/; RRID:SCR_001905 |
| Seurat (version 4.1.0) | Stuart et al.[1] | https://satijalab.org/seurat/get_started.html; RRID:SCR_016341 |
| Model-based analysis of single cell transcriptomics (MAST) | Finak et al.[2] | https://bioconductor.org/packages/release/bioc/html/MAST.html; RRID:SCR_016340 |
| Uniform Manifold Approximation and Projection (UMAP) | Melville et al.[3], 2018 #404} | https://github.com/lmcinnes/umap; RRID:SCR_018217 |
| edgeR (version 4.2.2) | Robinson et al.[4] | http://bioconductor.org/packages/edgeR/; RRID:SCR_012802 |

| | | |
|---|---|---|
| WebGestaltR 2019 | Zhang et al.[5] | https://cran.r-project.org/package=WebGestaltR; RRID:SCR_024312 |
| ggplot2 (version 3.5.2) | Wickham[6] | https://cran.r-project.org/web/packages/ggplot2/index.html; RRID:SCR_014601 |
| ImageJ | Schneider et al.[7] | https://imagej.net/; RRID:SCR_003070 |
| Imaris (version 10.0) | Oxford Instruments | http://www.bitplane.com/imaris/imaris; RRID:SCR_007370 |
| Axon™pCLAMP™ 10 Electrophysiology Data Acquisition & Analysis | Molecular Devices | http://www.moleculardevices.com/products/software/pclamp.html; RRID:SCR_011323 |
| Rstatix (version 0.7.2) | Kassambara et al.[8] | https://CRAN.R-project.org/package=rstatix; RRID:SCR_021240 |
| **Other** | | |
| Resource website for analyzed single-cell RNA sequencing data | This paper | https://anratherlab.shinyapps.io/angii_brain/ |


1. Stuart, T., Butler, A., Hoffman, P., Hafemeister, C., Papalexi, E., Mauck, W.M., 3rd, Hao, Y., Stoeckius, M., Smibert, P., and Satija, R. (2019). Comprehensive Integration of Single-Cell Data. Cell *177*, 1888-1902.e1821. 10.1016/j.cell.2019.05.031.
2. Finak, G., McDavid, A., Yajima, M., Deng, J., Gersuk, V., Shalek, A.K., Slichter, C.K., Miller, H.W., McElrath, M.J., Prlic, M., et al. (2015). MAST: a flexible statistical framework for assessing transcriptional changes and characterizing heterogeneity in single-cell RNA sequencing data. Genome Biology *16*, 278. 10.1186/s13059-015-0844-5.
3. Melville}, L.M.a.J.H.a.J. (2018). {UMAP: Uniform Manifold Approximation and Projection for Dimension Reduction}. *{1802.03426}* (*stat.ML*).
4. Robinson, M.D., McCarthy, D.J., and Smyth, G.K. (2010). edgeR: a Bioconductor package for differential expression analysis of digital gene expression data. Bioinformatics *26*, 139-140. 10.1093/bioinformatics/btp616.
5. Zhang, B., Kirov, S., and Snoddy, J. (2005). WebGestalt: an integrated system for exploring gene sets in various biological contexts. Nucleic Acids Res *33*, W741-748. 10.1093/nar/gki475.
6. Wickham, H. (2016). Elegant Graphics for Data Analysis.
7. Schneider, C.A., Rasband, W.S., and Eliceiri, K.W. (2012). NIH Image to ImageJ: 25 years of image analysis. Nat Methods *9*, 671-675. 10.1038/nmeth.2089.
8. Kassambara, A. (2023). rstatix: Pipe-Friendly Framework for Basic Statistical Tests.
